# An Azapeptide Platform in Conjunction with Covalent Warheads to Uncover High-Potency Inhibitors for SARS-CoV-2 Main Protease

**DOI:** 10.1101/2023.04.11.536467

**Authors:** Kaustav Khatua, Yugendar R. Alugubelli, Kai S. Yang, Veerabhadra R. Vulupala, Lauren R. Blankenship, Demonta D. Coleman, Sandeep Atla, Sankar P. Chaki, Zhi Zachary Geng, Xinyu R. Ma, Jing Xiao, Peng-Hsun Chase Chen, Chia-Chuan Dean Cho, Erol C. Vatansever, Yuying Ma, Ge Yu, Benjamin W. Neuman, Shiqing Xu, Wenshe Ray Liu

## Abstract

Main protease (M_Pro_) of SARS-CoV-2, the viral pathogen of COVID-19, is a crucial nonstructural protein that plays a vital role in the replication and pathogenesis of the virus. Its protease function relies on three active site pockets to recognize P1, P2, and P4 amino acid residues in a substrate and a catalytic cysteine residue for catalysis. By converting the P1 Cα atom in an M_Pro_ substrate to nitrogen, we showed that a large variety of azapeptide inhibitors with covalent warheads targeting the M_Pro_ catalytic cysteine could be easily synthesized. Through the characterization of these inhibitors, we identified several highly potent M_Pro_ inhibitors. Specifically, one inhibitor, MPI89 that contained an aza-2,2-dichloroacetyl warhead, displayed a 10 nM EC_50_ value in inhibiting SARS-CoV-2 from infecting ACE2_+_ A549 cells and a selectivity index of 875. The crystallography analyses of M_Pro_ bound with 6 inhibitors, including MPI89, revealed that inhibitors used their covalent warheads to covalently engage the catalytic cysteine and the aza-amide carbonyl oxygen to bind to the oxyanion hole. MPI89 represents one of the most potent M_Pro_ inhibitors developed so far, suggesting that further exploration of the azapeptide platform and the aza-2,2-dichloroacetyl warhead is needed for the development of potent inhibitors for the SARS-CoV-2 M_Pro_ as therapeutics for COVID-19.

## Introduction

Azapeptides are a class of molecules that contain a nitrogen atom, named as aza-nitrogen, in an original Cα position in the peptide backbone.^1^ This modification alters the electronic and steric configurations of the peptide, which can lead to improved features such as bioactivity and stability. In addition, the conversion of the Cα atom to a more nucleophilic nitrogen atom makes a large variety of chemical functionalities including many covalent warheads readily attached to it. One area where azapeptides have been frequently explored and shown promise is protease inhibitors.^2-4^ Azapeptides can inhibit protease activity by binding to the active site of the enzyme, often through the formation of a covalent bond with the catalytic residue. The presence of the aza-nitrogen atom in the peptide backbone can increase the stability of this covalent bond, leading to improved potency and selectivity. Azapeptide-based protease inhibitors have been previously developed and explored as potential therapeutics. Notable examples include several azapeptides developed for the HIV protease that have shown promising results in preclinical studies and those developed for human cathepsins that have been evaluated for their potential therapeutic applications.^5-8^ Azapeptides have also been previously explored as inhibitors for the SARS-CoV-2 main protease (M^Pro^) albeit with a relatively small scope.^9^ In this work, we wish to report an azapeptide platform for the exploration of a variety of covalent warheads to target the M^Pro^ catalytic cysteine and the identification of a potent azapeptide inhibitor, MPI89, with an antiviral EC_50_ value as low as 10 nM.

## Results and Discussion

M^Pro^, also known as 3C-like protease (3CL^Pro^), is a key enzyme in the replication of the SARS-CoV-2 virus that causes COVID-19.^10, 11^ It cleaves two large viral polypeptides into functional nonstructural proteins, which are necessary for the replication and transcription of the viral genome. Therefore, M^Pro^ is a crucial target for the development of antiviral drugs against COVID-19. Two drugs approved for emergency use that target M^Pro^ are nirmatrelvir and ensitrelvir.^12, 13^ These two molecules are also representative covalent and noncovalent M^Pro^ inhibitors, respectively. Although available, both nirmatrelvir and ensitrelvir have limitations. Nirmatrelvir is not a standalone drug and has to be used in combination with ritonavir that inhibits human cytochrome P450 3A4 to prolong nirmatrelvir’s half-life in the human body. Ensitrelvir has been shown to decrease the virus load but has not been effective in alleviating symptoms in patients. A looming prospect for both drugs is the acquired viral resistance to them.^14-17^ Therefore, while nirmatrelvir and ensitrelvir represent a significant step forward in the treatment of COVID-19, further development of M^Pro^ inhibitors is necessary to address their limitations and to stay ahead of the ongoing pandemic.

M^Pro^ is a 306aa cysteine protease. It possesses an active site comprising of four small pockets that bind to specific residues in a protein/peptide substrate, namely P1, P2, P4, and P1-3’ (Figure 1A). These pockets are denoted as S1, S2, S4, and S1’-3’ in accordance with the Schechter-Berger nomenclature (Figure 1B).^18^ The enzyme lacks a well-defined S3 pocket for accommodating the P3 side chain of the substrate. In most M^Pro^-peptide ligand structures, the P3 side chains were found to be exposed to the solvent.^12, 19-22^ The catalytic dyad of the enzyme comprises of two active site residues, C145 and H41. The catalysis involves C145 thiolate attacking the hydrolytic amide bond in the substrate, leading to the formation of a thioester intermediate that further undergoes hydrolysis to complete the catalytic cycle. To date, the primary focus of M^Pro^ inhibitor development has been directed towards the active site of M^Pro^. A vast array of these inhibitors, including nirmatrelvir, are peptide-based covalent inhibitors.^23-34^ We opted to systematically investigate the potential of azapeptides as M^Pro^ inhibitors, owing to their unique advantages in terms of synthesis and the ease of accessing a wide range of covalent warheads. Numerous attempts have been made to identify optimal amino acid sequences in M^Pro^ substrates.^35^ By replacing the P1 Cα atom with aza-nitrogen in a peptide substrate possessing a low K_m_ value, it is possible to obtain an azapeptide exhibiting potent binding to M^Pro^. In terms of synthesis, the hydrazide formed on the P2 C-terminal side provides high nucleophilicity, making it easy to attach a P1 side chain and a covalent warhead. In addition, the substitution of Cα with aza-nitrogen eliminates a chiral center, which simplifies the separation process of the resulting azapeptides.

**Figure 1.**
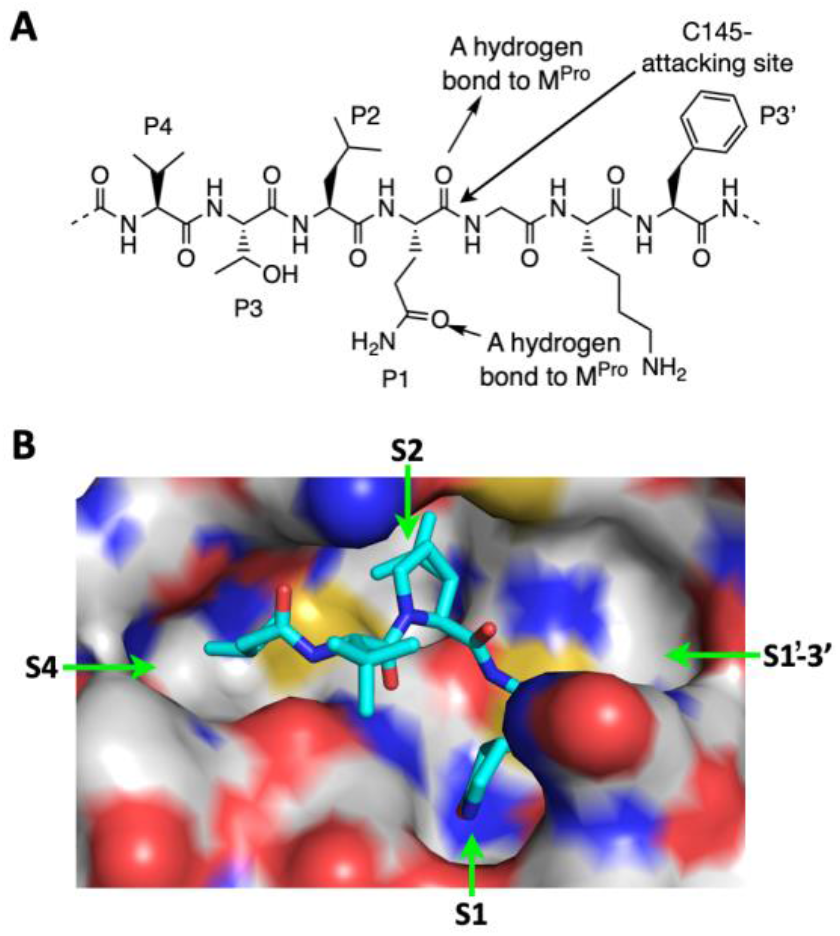
(**A**) A schematic illustration depicting the interactions between M^Pro^ and a protein/peptide substrate. The residues with respect to the hydrolytic site are labeled in accordance with the Schechter-Berger nomenclature. (**B**) A contoured M^Pro^ active site, highlighting S1, S2, S4, and S1-3’ pockets that bind a protein/peptide substrate. The configuration of the structure is derived from the PDB entry 7S75, wherein the ligand MPI42 is present as a covalent inhibitor of M^Pro^.

Our azapeptide design followed a general scheme as depicted in Figure 2. Previous research has explored the direct conjugation of nitrile to the aza nitrogen atom, resulting in a thioimidate adduct that can potentially be reversed (Figure 2A).^9^ This was integrated into our designs as a control. The U.S. FDA has approved multiple Michael acceptor-containing covalent inhibitors for therapeutic intervention in cancer,^36, 37^ and we incorporated this strategy into our design to explore these established Michael acceptors as covalent warheads for M^Pro^ (Figure 2B). The Michael addition adducts of M^Pro^ are also potentially reversible. 2-Haloacetyl groups are another group of covalent warheads for cysteine proteases (Figure 2C). Wang *et al*. have recently reported even 2,2-di- and 2,2,2-trihaloacetyl groups as covalent warheads for M^Pro^.^38^ We adopted some of their designs for our azapeptide inhibitor designs as well. Depending on the 2,2-haloacetyl groups used, covalent M^Pro^ adducts could be either reversible or irreversible.^38^ Azapeptide esters were originally developed as inhibitors for serine proteases and later found to serve as potent inhibitors for cysteine proteases as well.^4^ For cysteine proteases, azaglycine esters formed irreversible adducts, but covalent adducts with other azapeptide esters were slowly hydrolyzed (Figure 2D).^2^ We incorporated azapeptide esters into our design to evaluate whether this slow hydrolysis could be adjusted to achieve prolonged inhibition endurance. Another covalent warhead used for designing M^Pro^ inhibitors is α-ketoacyl. Boceprevir, which contains an α-ketoacyl moiety, served as a starting point for the development of multiple M^Pro^ inhibitors, including nirmatrelvir.^39^ The aza nitrogen’s availability allows for easy installation of an α-ketoacyl moiety, although the resulting α-ketoacyl will take a reversed configuration compared to that in boceprevir. We believed that this one-carbon switch of the ketone group would still allow reacting with the M^Pro^ catalytic cysteine to form a reversible covalent adduct, as depicted in Figure 2E. Additionally, we explored two other designs, as shown in Figures 2F and 2G, to investigate the likelihood of the M^Pro^ catalytic cysteine undergoing an S^N^Ar reaction with an electron-deficient halogenated benzene group and an S^N^2 reaction with an acetoxymethylcarbonyl compound, respectively. Similar reactions have been observed with M^Pro^.^32, 40^ We expected that both reactions would result in irreversible covalent adducts with M^Pro^.

**Figure 2.**
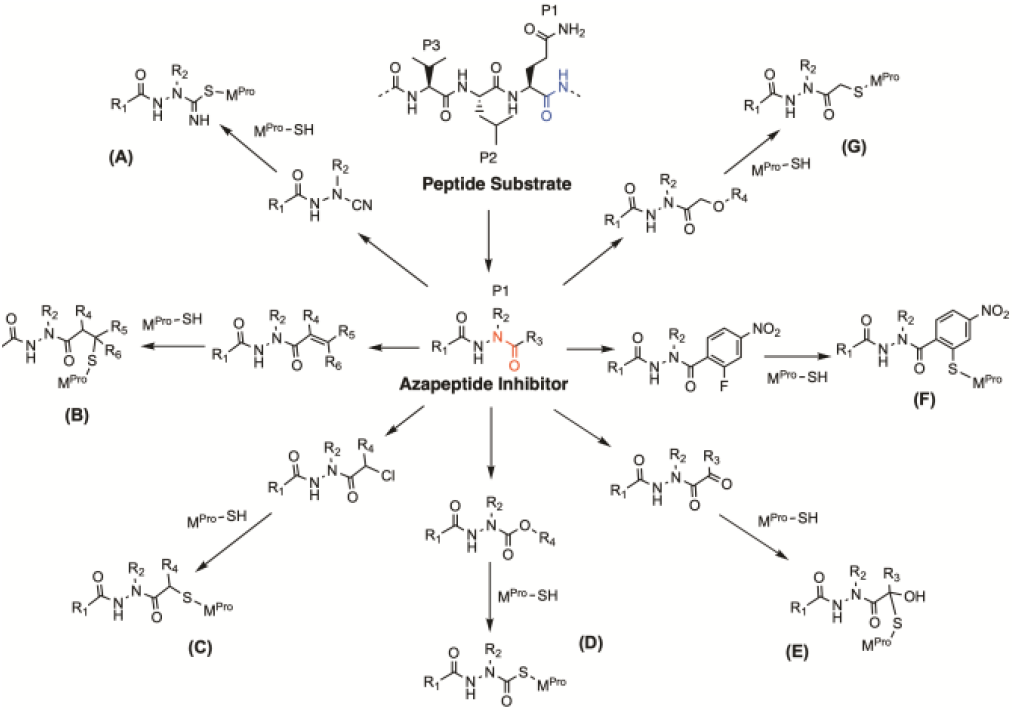
A graphic representation illustrating the transformation of an M^Pro^ substrate into an azapeptide for accessing various covalent warheads that can interact with the catalytic cysteine in the M^Pro^ active site.

GC376, which has been investigated as a treatment for feline infectious peritonitis, was also explored as a potential cure for COVID-19 during the early stages of the pandemic.^41, 42^ Our initial design and synthesis of azapeptides were based on GC376, which contains a P1 β-(*S*-2-oxopyrrolidin-3-yl)-alanal, a P2 leucine, and an *N*-terminal CBZ group. Our first group of azapeptides shown in Figure 3A all contained a P2 leucine, with a majority featuring an *N*-terminal CBZ group. The P1 side chain was selected as propanamide, similar to the native P1 glutamine in a protein substrate, for easy synthesis. Indole-2-carboxyl and 4-methoxy-indole-2-carboxyl groups, frequently used as *N*-terminal protection groups for improved potency and metabolic stability in dipeptidyl inhibitors of M^Pro^, were integrated into our overall design as well but with relatively minimal representation.^23^ We first synthesized a negative control molecule, AzaPep, that contains an aza-propanoyl group and continued to generate other molecules. Molecules in this group with potential covalent warheads for reacting with the M^Pro^ catalytic cysteine include MPI68 and MPI84 containing an aza-2-chloroacetyl warhead, MPI69 and MPI83 with aza-acryloyl, MPI70 with aza-*R*-2-chloropropanoyl, MPI71 with aza-*S*-2-chloropropanoyl, MPI72 with aza-2-fluoro-4-nitrobenzoyl, MPI73 with aza-2-benzylacryloyl, MPI74 with aza-(*E*)-4-(dimethylamino)but-2-enoyl, MPI75 with aza-2-(4-nitrophenoxy)acetyl, MPI76 with aza-4-fluorophenylcarbamoyl, MPI77 with aza-2,2,2-trifluoroethylcarbamoyl, MPI78 with aza-2-(4-fluorophenoxy)acetyl, MPI90 with aza-pyruvoyl, MPI92 with aza-2,2-dichloroacetyl, and MPI93 with aza-nitrile.

**Figure 3.**
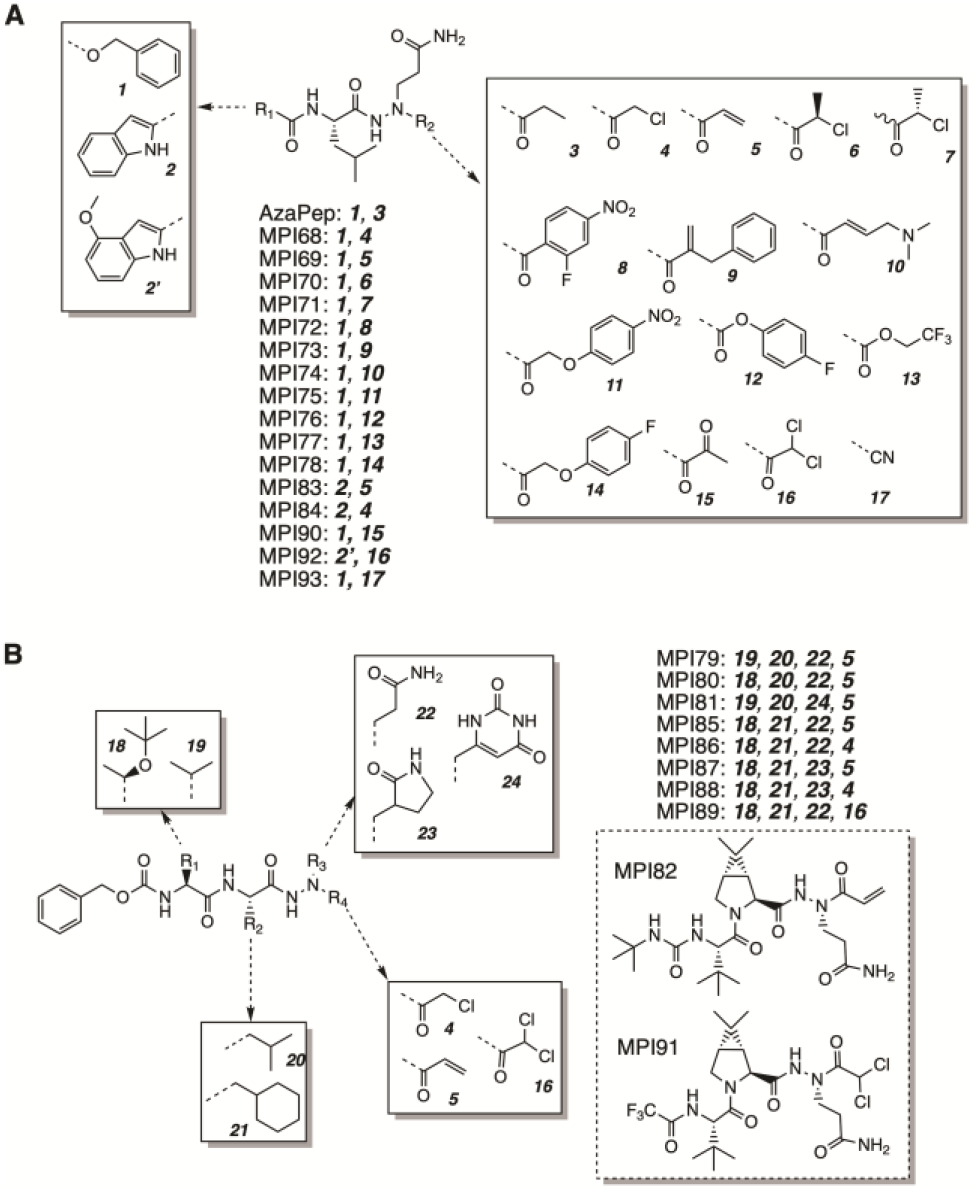
Structures of all azapeptide compounds that were designed and synthesized.

Following the identification of the three most effective covalent warheads, namely aza-2-chloroacetyl, aza-acryloyl, and aza-2,2-dichloroacetyl, in the first group of azapeptide molecules, we developed more azapeptide compounds based on the results of multiple structure-activity relationship (SAR) studies of tripeptidyl inhibitors and nirmatrelvir.^21, 22^ The structures of these molecules are shown in Figure 3B. MPI79 and MPI81 contain an aza-acryloyl warhead and were derived from MPI3.^43^ Specifically, MPI81 has a 6-methyluracil moiety at P1 to evaluate potential interactions between the uracil moiety that was used in some M^Pro^ inhibitors and the S1 pocket of M^Pro^.^44^ MPI80, which has an aza-acryloyl group, was developed from MPI6.^43^ MPI85-MPI89 are all based on MPI8, a covalent inhibitor of M^Pro^ that utilizes an aldehyde as a covalent warhead and exhibits significant antiviral activity.^43, 45^ To better mimic the structure of MPI8, a P1 3-methylpyrrolidin-2-one was incorporated into MPI87 and MPI88. MPI89 maintained a P1 propanamide. Unlike MPI87 and MPI88, which contain aza-acryloyl and aza-2-chloroacetyl respectively, MPI89 features an aza-2,2-dichloroacetyl group. MPI46 is a potent aldehyde-based inhibitor of M^Pro^.^22^ MPI82, which contains an aza-acryloyl warhead, was designed based on the structure of MPI46. When the structure of nirmatrelvir was released, MPI91 was developed based on it. MPI91 has an an aza-2,2-dichloroacetyl as the warhead group. All structures are presented in Figure S1.

To characterize the synthesized compounds, we determined their enzymatic inhibition IC_50_ values, cellular EC_50_ values in inhibiting M^Pro^ recombinantly expressed in 293T cells, antiviral EC_50_ values in preventing SARS-CoV-2 from infecting ACE2^+^ A549 cells, and cytotoxic CC^50^ values in killing 293T cells, following previously established practices. In a previous drug repurposing project, we established a protocol to characterize IC_50_ values for M^Pro^ inhibitors.^46^ The protocol involves using a fluorogenic substrate called Sub3, which is hydrolyzed by M^Pro^ to produce a fluorogenic product that can be detected by a plate reader. By adding an inhibitor to M^Pro^, we could inhibit this process of fluorescence release. We have previously used this protocol to characterize over 100 M^Pro^ inhibitors,^47^ and we applied the same protocol to determine the IC_50_ values for all azapeptide inhibitors developed in this study. To conduct the assay, we preincubated 20 nM M^Pro^ with varying concentrations of an inhibitor for 30 min before adding 10 µM Sub3 and recorded the formation of fluorescent products. We then used the product formation rates at different inhibitor concentrations to calculate the IC_50_ values by fitting the data to a three-parameter inhibition equation using GraphPad (Figure S2). All the IC_50_ values determined are presented in Table 1.

**Table 1.** Determined enzymatic IC_50_ and cellular/antiviral EC_50_ values of M^Pro^ inhibitors

| ID | Enzymatic IC<sub>50</sub> (μM) | Cellular EC<sub>50</sub> (μM) | Antiviral EC<sub>50</sub> (μM) | CC<sub>50</sub> (μM) | ID | Enzymatic IC<sub>50</sub> (μM) | Cellular EC<sub>50</sub> (μM) | Antiviral EC<sub>50</sub> (μM) | CC<sub>50</sub> (μM) |
| --- | --- | --- | --- | --- | --- | --- | --- | --- | --- |
| AzaPep | > 10 |  |  |  | MPI81 | 3.2 | > 10 |  |  |
| MPI68 | 0.088 | > 10 |  |  | MPI82 | 0.13 | 4.2 |  |  |
| MPI69 | 0.51 | > 10 |  |  | MPI83 | 0.049 | > 10 |  |  |
| MPI70 | 5.4 | > 10 |  |  | MPI84 | 0.041 | 0.22 | 0.53 | 78.2 |
| MPI71 | 1.6 | > 5 |  |  | MPI85 | 0.063 | 0.33 | 1.8 | 41.8 |
| MPI72 | > 10 |  |  |  | MPI86 | 0.28 | 0.15 | 0.38 | 59.4 |
| MPI73 | > 10 | 0.15 |  |  | MPI87 | 0.19 | 2.8 |  |  |
| MPI74 | > 10 |  |  |  | MPI88 | 0.29 | 0.21 | 0.33 | 52 |
| MPI75 | > 10 |  |  |  | MPI89 | 0.068 | 0.035 | 0.010 | 59.5 |
| MPI76 | 0.070 | > 10 |  |  | MPI90 | > 5 |  |  |  |
| MPI77 | 1.56 | > 10 |  |  | MPI91 | 0.024 | > 5 |  |  |
| MPI78 | > 10 |  |  |  | MPI92 | 0.14 | > 10 |  | 54.6 |
| MPI79 | 0.053 | 3.5 |  |  | MPI93 | > 10 |  |  |  |
| MPI80 | 0.034 | 0.84 | 0.70 | 54.6 | Nirmatrelvir | 0.066 | 3.4 | 0.068 | 31 |

As anticipated, AzaPep, which was the negative control molecule, did not exhibit any inhibition of M^Pro^. MPI68 and MPI69 both demonstrated IC_50_ values below 1 µM, with MPI68 being more potent than MPI69. During the M^Pro^-catalyzed hydrolysis of a substrate, the M^Pro^ C145 thiolate attacks the P1 *C*-terminal carbonyl carbon to produce a thioester covalent intermediate. In MPI68 and MPI69, the covalent attacking site for the M^Pro^ C145 thiolate changes by one and two carbon positions, respectively. This deviation from the original attacking site may account for the difference in potency between MPI68 and MPI69. MPI70 and MPI71 were derived from MPI68, but with an additional methyl group added to the aza-2-chloroacetyl 2-carbon. Both compounds displayed lower M^Pro^ inhibition potency than MPI68 due to the increased steric hindrance at the covalent attacking site. However, MPI71, which has an *S* configuration at the aza-chloroacetyl 2-carbon, was more potent than MPI70, which has an *R* configuration at the same carbon. The proposed covalent adduct between M^Pro^ and MPI71 would have the added methyl group pointing towards the empty S1’-3’ pocket, likely explaining why MPI71 is more potent than MPI70. This finding suggests that a bigger chemical moiety can be added to the aza-chloroacetyl α-carbon, that is in its *S* configuration, to enhance the interaction with the S1’-3’ pocket of M^Pro^ for improved inhibition strength. MPI72-MPI75 had very low inhibition potency towards M^Pro^. For MPI73 and MPI74, the low inhibition potency may be due to the added steric hindrance and clash with M^Pro^ caused by the addition of large chemical moieties to the aza-acryloyl warhead. MPI72 was created to test a possible S^N^Ar reaction with M^Pro^, but its low potency may be due to no S^N^Ar reaction occurring between M^Pro^ and MPI72. The 2-fluoro-4-nitrobenzoyl group is relatively large, so careful tuning is necessary for binding to the active site while simultaneously reaching the M^Pro^ C145 thiolate for reaction. MPI75 was designed to test a possible reaction between the M^Pro^ C145 thiolate and aza-2-(4-nitrophenoxy)acetyl group, but it did not inhibit M^Pro^. MPI78 is similar to MPI75 but has a different leaving group and was also inactive. Acetoxymethylketone is frequently used for covalent inhibition of serine and cysteine proteases.^32, 48^ The currently accepted mechanism involves the addition of the ketone to generate a hemiacetal or thiohemiacetal, which then rearranges to replace the acetoxy group by the active site serine or cysteine for irreversible covalent adduct formation. In both MPI75 and MPI78, the presence of the aza-nitrogen atom leads to an inactive amide rather than active ketone. This may explain that both compounds are inactive against M^Pro^.

MPI76 and MPI77 are two compounds that contain an azapeptide ester group. Both compounds displayed significant inhibition activity, with MPI76 being 20 times more potent than MPI77 as determined by their respective IC_50_ values. The high potency of MPI76 may be attributed to the fact that 4-fluorophenol is an easy leaving group. MPI83 and MPI84 were subsequently synthesized for comparison to MPI68 and MPI69, and both contain an *N*-indole-2-carboxyl protection group. They exhibited remarkable potency, with IC_50_ values lower than 50 nM. *N*-Indole-2-carboxyl is more rigid than CBZ group. Earlier crystallography research indicated that the *N*-indole-2-carboxyl protection group in dipeptide M^Pro^ inhibitors produced a hydrogen bond between the indole nitrogen and the M^Pro^ E166 backbone carbonyl oxygen.^49^ Both factors may contribute to the strong potency of MPI83 and MPI84. MPI90 was also developed to investigate if an aza-conjugated α-ketoacyl, specifically here as a pyruvoyl group, could serve as a covalent warhead to react with the M^Pro^ catalytic cysteine. Unfortunately, this substance was not very effective. The pyruvoyl ketone group in MPI90 may not bind to the enzyme active site at an optimal configuration to react with the M^Pro^ C145 thiolate. MPI92, an analogue of MPI84, was developed, containing an aza-2,2-dichloroacetyl warhead compared to an aza-2-chloroacetyl warhead in MPI84. It has 10 times less inhibition potency than MPI84, as the aza-2,2-dichloroacetyl warhead is theoretically less reactive towards cysteine than the aza-2-chloroacetyl warhead. Based on IC_50_ values of group A compounds, it can be concluded that aza-2-chloroacetyl, aza-acryloyl, and azapeptide ester warheads can result in potent inhibitors. The aza-2-chloroacetyl group can be further modified to reach the S1’-3’ pocket in M^Pro^ for better binding. Since non-azaglycine esters generate slowly hydrolysable adducts with a cysteine protease and MPI76 is not significantly superior to MPI68, we excluded azapeptide esters from our further designed molecules. Therefore, in group B compounds, as shown in Figure 3B, we retained aza-2-chloroacetyl, aza-acryloyl, and aza-2,2-dichloroacetyl warheads.

The compounds in Group B were created using tripeptidyl M^Pro^ inhibitors with high enzymatic inhibition or antiviral potency as a basis for design. Both MPI79 and MPI80 demonstrated potent M^Pro^ inhibition, with very similar IC_50_ values. The only difference between the two inhibitors was at the P3 site, and since M^Pro^ lacks a defined S3 pocket for the binding of the P3 side chain, the similar IC_50_ values for the two compounds are likely due to this reason. MPI81 differs from MPI79 only at the P1 side chain, with a measurable IC_50_ value that was 60-fold higher than MPI79, indicating that the uracil side chain requires careful turning for optimal binding to the S1 pocket. MPI85 and MPI86 were based on MPI8 but contain a P1 propanamide side chain. Both demonstrated high M^Pro^ inhibition potency, with MPI85 being about four times more potent than MPI86. MPI87 differs from MPI85 at the P1 side chain, containing 3-methylpyrrodin-2-one that is present in nirmatrelvir and many other potent M^Pro^ inhibitors. It displayed reduced potency compared to MPI85, likely because the aza-nitrogen forces 3-methylpyrrolidin-2-one to adopt a configuration different from that in nirmatrelvir. MPI88, an MPI86 analogue, contains a P1 3-methylpyrrolidin-2-one side chain and demonstrated slightly lower potency than MPI86, likely for the same reason discussed for MPI87. MPI89 is an MPI86 analogue, containing an aza-2,2-dichloroacetyl warhead. Although it was anticipated that MPI89 would display lower potency due to the less reactive warhead, its IC_50_ value of 68 nM was four times lower than MPI86. This may be because the additional chloride on the warhead participates in unique interactions with M^Pro^ for improved binding and reaction. MPI82 and MPI91 were based on MPI46 and nirmatrelvir, respectively, and both demonstrated strong M^Pro^ inhibition potency. MPI91 was slightly better, with a determined IC_50_ value below 100 nM. All Group B compounds were based on optimized binding moieties for M^Pro^ active site pockets and covalent warheads.

We went further to characterize compounds with measurable M^Pro^ inhibition potency by testing their inhibition of M^Pro^ recombinantly expressed in 293T cells. M^Pro^ causes acute toxicity and drives host cells to undergo apoptosis when expressed in a human cell host. Based on this unique feature, we previously developed a cell-based assay to characterize cellular potency of M^Pro^ inhibitor.^50^ In this assay, an inhibitor with cellular potency suppresses cytotoxicity from an M^Pro^-eGFP fusion protein ectopically expressed in 293T cells and consequently leads to host cell survival and enhanced overall expression of M^Pro^-eGFP that can be characterized using flow cytometry. Using this novel cellular assay system, we have been prioritizing M^Pro^ inhibitors for antiviral assays that need to be conducted in a BSL3 facility. Using the same assay, we characterized all azapeptide compounds that exhibited measurable IC_50_ values below 5 µM (Figure S3). The determined cellular EC_50_ values are presented in Table 1. For Group A compounds, except for MPI84, they all displayed very low cellular potency. This is consistent with other dipeptidyl M^Pro^ inhibitors and is likely due to low cellular permeability and stability of this group of compounds.^43^ MPI84 showed a determined cellular EC_50_ value of 220 nM, indicating that the *N*-terminal Indole-2-carboxyl group helped with cellular permeability or stability. For Group B compounds that were tested, except for MPI81 and MPI91, they all showed measurable cellular EC_50_ values. MPI80, MPI85, MPI86, MPI88, and MPI89 exhibited determined cellular EC_50_ values below 1 µM, with MPI89 having the lowest determined EC_50_ value of 35 nM, equivalent to MPI8, the inhibitor with the lowest cellular EC_50_ value among all inhibitors (> 200) that we have tested so far.^50^ Overall, triazapeptidyl inhibitors outperformed diazapeptidyl inhibitors in inhibiting M^Pro^ recombinantly expressed in 293T cells.

We went further to conduct antiviral potency tests for six azapeptide inhibitors, namely MPI80, MPI84, MPI85, MPI88, and MPI89, in ACE2^+^ A549 cells infected with hCoV-19/USA/HP05647/2021, an early Delta variant that grows well in cell culture and produces strong cytopathic effect. The inhibitors were added at varying concentrations, and the cells were grown for three days before quantifying SARS-CoV-2 mRNAs using RT-PCR to determine antiviral EC_50_ values. Our findings, presented in Table 1 and Figure S4, indicate that all six compounds demonstrated measurable antiviral EC_50_ values, comparable to their cellular EC_50_ values in inhibiting recombinantly expressed M^Pro^ in 293T cells. MPI89 emerged as the most potent, with a determined antiviral EC_50_ value of 10 nM, while nirmatrelvir, the active M^Pro^ inhibitor component in Paxlovid, had a determined antiviral EC_50_ value of 68 nM in the same experimental setup. Of the remaining five tested azapeptide compounds, MPI86 was the most potent, with a determined antiviral EC_50_ value of 0.38 µM, which is 38 times higher than that for MPI89. The high antiviral potency of both MPI86 and MPI89 was expected, given that they were designed based on MPI8, an aldehyde-based inhibitor with high antiviral potency. Compared to MPI86, MPI89 contains an additional chloride, which is likely involved in both the binding of M^Pro^ and improving the stability and/or cellular permeability of MPI89. We also evaluated the cytotoxicity of the six compounds in 293T cells using the MTT assay and found their determined CC^50^ values (presented in Table 1 and Figure S5) to be similar to that of nirmatrelvir. Notably, MPI89’s CC^50^ value of 59.5 µM leads to a selectivity index of 875, indicating its potential safety for use as an antiviral. In addition, we assessed the stability of MPI80, MPI85, MPI86, and MPI89 in human liver microsomes (Figure S6). The inhibitors were incubated in presence or absence of nicotinamide adenine dinucleotide phosphate (NADPH) at 37 °C. LC-MS/MS analyses have been used to study the degradation of the inhibitors. While MPI80 had a determined half-life of 62 minutes, the other three compounds had a half-life of approximately 20 minutes. The relatively short half-life of MPI89 in human liver microsomes suggests that further improvements are necessary before it can be considered for an animal efficacy study.

In order to investigate how different azapeptides interact with the M^Pro^ active site, we utilized X-ray protein crystallography to characterize their binding. Initially, apo M^Pro^ was crystalized and then soaked with several azapeptides, including MPI79, and MPI85-MPI89. The resulting crystals were then subjected to X-ray diffraction, data collection, and structural refinement. The determined structures of M^Pro^ with all six azapeptide inhibitors bound at its active site are shown in Figure 4. Since all six azapeptide inhibitors are tripeptide-based, their M^Pro^ complexes can be compared to those of other tripeptide inhibitors, such as MPI3 and MPI8. All six azapeptide inhibitors formed 9 or 10 hydrogen bonds with M^Pro^ active site residues, similar to what were observed for M^Pro^ complexed with other tripeptide inhibitors.^43^ Specifically, in all structures, the aza-amide carbonyl oxygen formed three hydrogen bonds with the backbone amines from M^Pro^ residues G143, S144, and C145 that form the oxyanion hole. Moreover, a hydrogen bond was generated between the P1 backbone amine and the backbone carbonyl oxygen of M^Pro^ H164. The P2 residue of each azapeptide inhibitor utilized its backbone amine and carbonyl oxygen to form two hydrogen bonds with the backbone carbonyl oxygen and amine, respectively, of M^Pro^ E166. Additionally, the P1 side chain amide of all azapeptides was involved in three hydrogen bonds: one through its own amide carbonyl oxygen with an imidazole nitrogen from M^Pro^ H163, and two through its amide amine with the side chain carboxylate of M^Pro^ E166 and the backbone carbonyl oxygen of M^Pro^ F140. Furthermore, except for MPI79, all other azapeptides formed a hydrogen bond with the side chain amide of M^Pro^ Q189 using their P2 backbone amine.

**Figure 4.**
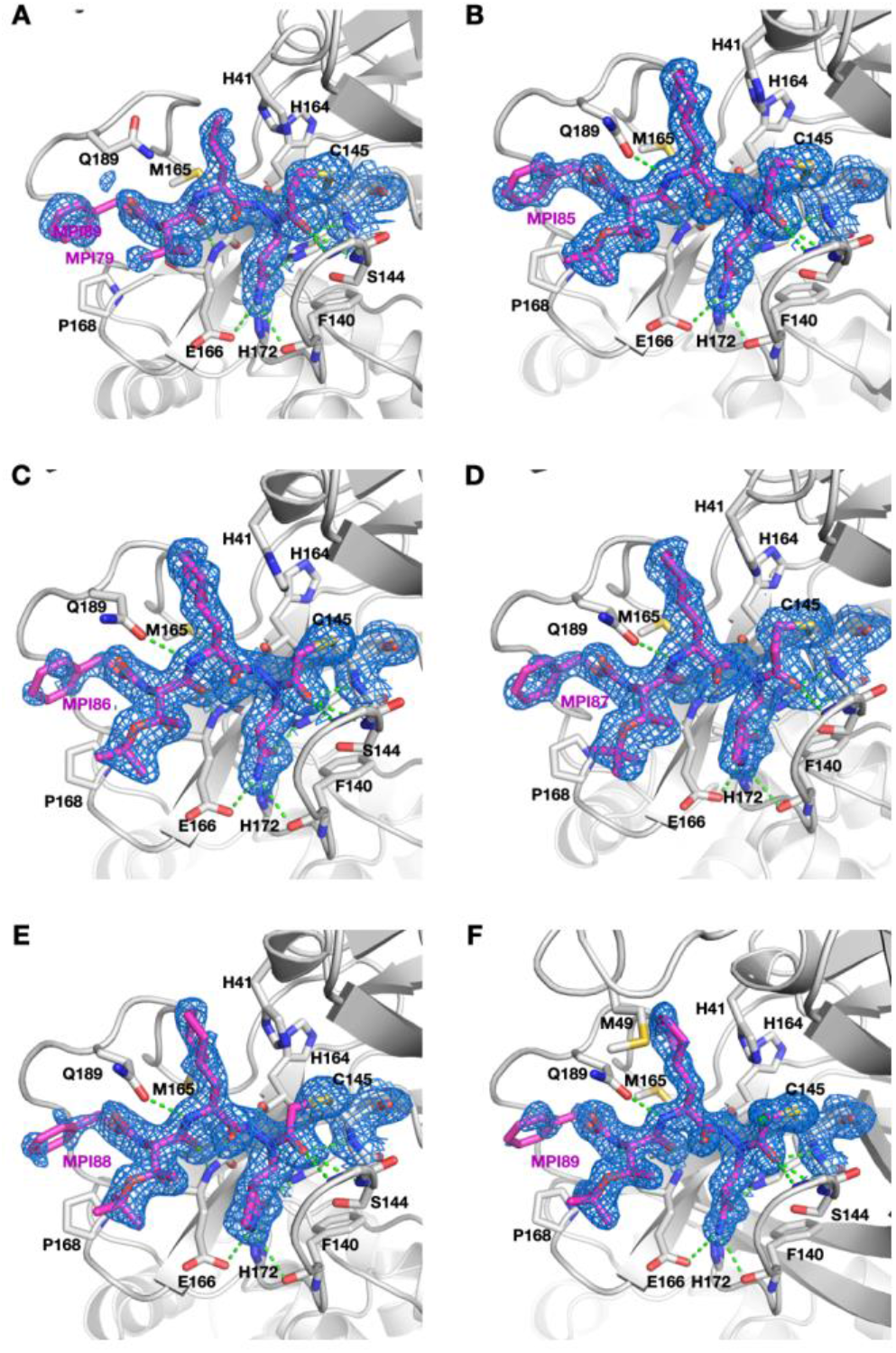
Crystal structures of M^Pro^ bound with MPI79 (**A**), MPI85 (**B**), MPI86 (**C**), MPI87 (**D**), MPI88 (**E**), and MPI89 (**F**).

MPI79 was designed based on MPI3. Its complex with M^Pro^ is structurally similar to the M^Pro^-MPI3 complex except around the P1 residue and warhead (Figure S7A). The electron density around the aza-nitrogen atom clearly showed three coplanar bonds, eliminating the chiral center at the original P1 Cα position (Figure 4A). MPI79 has an aza-acrylamide warhead, and in the refined structure, there was strong continuous electron density connecting the aza-acrylamide warhead and the M^Pro^ C145 thiolate, indicating the formation of a covalent bond between the two. In most previously determined M^Pro^ structures in complex with aldehyde and nitrile-based inhibitors, the C145 thiolate formed a covalent bond with the warhead but still pointed towards H41. However, in M^Pro^-MPI79, the C145 thiolate rotated about 180° and pointed towards the solvent, to accommodate the vinyl group added to the aza-amide of the inhibitor for the formation of a covalent bond. Compared to MPI3, the aza-amide carbonyl group that is equivalent to the P1 carbonyl group in MPI3 was also pushed away from M^Pro^ C145, likely to accommodate the added vinyl group for the covalent bond formation. These observations explain how M^Pro^ covalently engages an azapeptide inhibitor with an aza-acryloyl warhead, even though the covalent interaction site in the inhibitor is two carbons away from the original P1 carbonyl carbon in a substrate.

MPI85 has an aza-acryloyl warhead and a binding mode to the M^Pro^ active site like that for MPI79 (Figure 4B), except that their P2 and P3 residues are different. MPI85 is also a structure analogue of MPI8. When comparing the M^Pro^-MPI85 and M^Pro^-MPI8 complexes (Figure S7B), the binding patterns for P2, P3, and the *N*-terminal CBZ groups are almost identical, with differences found only at the P1 and warhead positions. Specifically, the three bonds to the aza-nitrogen adopted a coplanar structure, and the M^Pro^ C145 thiolate rotated 180° from its original position in the apo enzyme, pointing towards the solvent to create enough space for forming a covalent bond with the aza-acryloyl warhead. MPI86 is similar to MPI85, differing only at the warhead, which is aza-2-chloroacetyl. As shown in Figure 4C, the electron density at the M^Pro^ active site revealed the loss of the chloride atom and the formation of a covalent bond between the M^Pro^ C145 thiolate and the aza-acetyl group in MPI86. When comparing the M^Pro^-MPI86 and M^Pro^-MPI8 complexes (Figure S7C), the two ligands have almost identical binding patterns at the P2, P3, and the *N*-terminal groups. However, the M^Pro^ C145 thiolate rotated about 180° from its position in M^Pro^-MPI8 to form a solvent-exposed covalent bond with the aza-acetyl group in MPI86. Although MPI86 and MPI85 have almost identical interaction patterns with M^Pro^, MPI85 has one more carbon, which forced the aza-amide group to tilt more towards the solvent (Figure S7D).

As shown in Figure 4D, MPI87 binds to M^Pro^ in a similar manner as MPI85, with the only difference being the P1 side chain (Figure S7E). While MPI85 has a propanamide side chain, MPI87 has a lactam side chain. When the P1 propanamide side chain in other peptide-based M^Pro^ inhibitors was replaced with a lactam side chain, most inhibitors showed better enzyme inhibition potency. However, this was not the case for azapeptide-based M^Pro^ inhibitors, as MPI87 is a weaker inhibitor than MPI85. The superposition of M^Pro^-MPI87 and M^Pro^-MPI8 (Figure S7F) shows similar binding patterns for the two P1 lactam side chains, with the aza-nitrogen in MPI87 being slightly further from the oxyanion hole of M^Pro^ compared to the P1 Cα in MPI8. The deviation may contribute to the less favored binding of the P1 lactam side chain in MPI87. MPI88 is a structural homolog of MPI87, but with an aza-2-chloroacetyl warhead. Its structure, shown in Figure 4E, reveals a binding pattern in the M^Pro^ active site similar to that of MPI87. When superimposed with M^Pro^-MPI87, the only difference between the two is the warhead position. The aza-amide moiety in MPI87 is more tilted towards the solvent compared to MPI88, due to its one additional carbon in the warhead. On the other hand, MPI89 has one additional chloride atom compared to MPI86. Its covalent complex with M^Pro^, shown in Figure 4F, displays electron density at the C2 carbon of the aza-acetyl group that points towards the solvent, indicating the presence of an unreacted chloride atom in the final covalent complex. Aside from this additional chloride atom, the structure of M^Pro^-MPI89 is highly similar to that of M^Pro^-MPI86, as shown by their superposition (Figure S7H).

In a previous report, we demonstrated that three M^Pro^ residues (C22, C41, and Y61) used two oxygen atoms to mediate the formation of an S-O-N-O-S crosslink of their side chain heteroatoms, which serves as a potential redox mechanism to regulate M^Pro^ activity.^51^ In all M^Pro^-azapeptide inhibitor complexes, except M^Pro^-MPI89, the same Y-shaped S-O-N-O-S crosslink was observed (Figure S8). Furthermore, the aa46-50 region had a poor configuration in all five structures with this crosslink, leading to an open S2 pocket, which is similar to what has been observed in all other M^Pro^ structures with this crosslink.^52^ As discussed in our previous report, we believe this open S2 pocket will allow the design and development of a brand-new series of M^Pro^ inhibitors.

## Conclusion

In summary, we systematically explored azapeptides in combination with various covalent warheads as a platform for developing inhibitors for SARS-CoV-2 M^Pro^. Through a search of azadipeptide compounds with warheads potentially undergoing Michael addition, S^N^2, S^N^Ar, nucleophilic acyl substitution, and thiohemiacetal formation reactions with the M^Pro^ C145 thiolate, we identified aza-acryloyl, aza-2-chloroacetyl, aza-2,2-dichloroacetyl, and azapeptide ester as active warheads to covalently engage the M^Pro^ active site cysteine. Focusing on the first three covalent warheads that led to strong enzymatic inhibition potency for azadipeptides, we developed azatripeptide inhibitors and found MPI89, with an aza-2,2-dichloroacetyl warhead, to have high antiviral potency with an EC_50_ value of 10 nM and a selective index of 875. MPI89 represents one of the most potent anti-SARS-CoV-2 compounds developed. Although MPI89 has a relatively short *in vitro* metabolic half-life that hinders its practical use as an antiviral, its high antiviral potency and selectivity index indicate that the azapeptide platform combined with the aza-2,2-dichloroacetyl warhead is a practical route to generate M^Pro^ inhibitors with high antiviral potency. Further development in this direction will likely lead to compounds that can be practically used as antivirals. Additionally, our study revealed an interesting observation that the aza-2,2-dichloroacetyl warhead showed higher potency than the aza-2-chloroacetyl warhead, contrary to what was predicted from the reactivities of the two warheads. The underlying reason for the aza-2,2-dichloroacetyl warhead’s high potency requires further investigation.

## EXPERIMENTAL SECTION

### Materials

We purchased yeast extract from Thermo Fisher Scientific, tryptone from Gibco, Sub3 from Bachem, HEK 293T/17 cells from ATCC, DMEM with GlutaMax from Gibco, FBS from Gibco, polyethyleneimine from Polysciences, and the trypsin-EDTA solution from Gibco. Chemicals used in this work were acquired from Sigma Aldrich, Chem Impex, Ambeed, and A2B. Pooled human liver microsome (1910096) was obtained from Xenotech.

### M^Pro^ Expression and Purification

The expression plasmid pET28a-His-SUMO-M^Pro^ was constructed in a previous study. We used this construct to transform *E. coli* BL21(DE3) cells. A single colony grown on an LB plate containing 50 µg/mL kanamycin was picked and grown in 5 mL LB media supplemented with 50 µg/mL kanamycin overnight. We inoculated this overnight culture to 6 L 2YT media with 50 µg/mL kanamycin. Cells were grown to OD^600^ as 0.8. At this point, we added 1 mM IPTG to induce the expression of His-SUMO-M^Pro^. Induced cells were let grown for 3 h and then harvested by centrifugation at 12,000 rpm, 4 °C for 30 min. We resuspended cell pellets in 150 mL lysis buffer (20 mM Tris-HCl, 100 mM NaCl, 10 mM imidazole, pH 8.0) and lysed the cells by sonication on ice. We clarified the lysate by centrifugation at 16,000 rpm, 4 °C for 30 min. We decanted the supernatant and mixed with Ni-NTA resins (GenScript). We loaded the resins to a column, washed the resins with 10 volumes of lysis buffer, and eluted the bound protein using elution buffer (20 mM Tris-HCl, 100 mM NaCl, 250 mM imidazole, pH 8.0). We exchanged buffer of the elute to another buffer (20 mM Tris-HCl, 100 mM NaCl, 10 mM imidazole, 1 mM DTT, pH 8.0) using a HiPrep 26/10 desalting column (Cytiva) and digested the elute using 10 units SUMO protease overnight at 4 °C. The digested elute was subjected to Ni-NTA resins in a column to remove His-tagged SUMO protease, His-tagged SUMO tag, and undigested His-SUMO-M^Pro^. We loaded the flow-through onto a Q-Sepharose column and purified M^Pro^ using FPLC by running a linear gradient from 0 to 500 mM NaCl in a buffer (20 mM Tris-HCl, 1 mM DTT, pH 8.0). Fractions eluted from the Q-Sepharose column was concentrated and loaded onto a HiPrep 16/60 Sephacryl S-100 HR column and purified using a buffer containing 20 mM Tris-HCl, 100 mM NaCl, 1 mM DTT, and 1 mM EDTA at pH 7.8. The final purified was concentrated and stored in a -80 °C freezer.

### *In Vitro* M^Pro^ Inhibition Potency Characterizations of MPIs

For most MPIs, we conducted the assay using 20 nM M^Pro^ and 10 µM Sub3. We dissolved all inhibitors in DMSO as 10 mM stock solutions. Sub3 was dissolved in DMSO as a 1 mM stock solution and diluted 100 times in the final assay buffer containing 10 mM Na^x^H^y^PO^4^, 10 mM NaCl, 0.5 mM EDTA, and 1.25% DMSO at pH 7.6. We incubated M^Pro^ and an inhibitor in the final assay buffer for 30 min before adding the substrate to initiate the reaction catalyzed by M^Pro^. The production format was monitored in a fluorescence plate reader with excitation at 336 nm and emission at 490 nm.

### Ray Crystallography Analysis of M^Pro^-Inhibitor Complexes

The production of crystals of Mpro -inhibitor complexes was done following the previous protocol.^53^ The data of Mpro with MPI79, MPI85, MPI86, MPI87, MPI88, and MPI89 were collected on a Bruker Photon II detector. The diffraction data were indexed, integrated, and scaled with PROTEUM3. All the structures were determined by molecular replacement using the structure model of the free enzyme of the SARS-CoV-2 Mpro [Protein Data Bank (PDB) ID code 7JPY] as the search model using Phaser in the Phenix package. JLigand and Sketcher from the CCP4 suite were employed for the generation of PDB and geometric restraints for the inhibitors. The inhibitors were built into the 2Fo-Fc density by using Coot. Refinement of all the structures was performed with Real-space Refinement in Phenix. Details of data quality and structure refinement are summarized in Table S1. All structural figures were generated with PyMOL.

### *In cellulo* M^Pro^ Inhibition Potency Characterizations of MPIs

HEK 293T/17 cells were grown in high-glucose DMEM with GlutaMAX Supplement and 10% fetal bovine serum in 10 cm culture plates under 37 °Cand 5% CO^2^ to 80%∼90%, followed by transfecting the cells with the pLVX-M^Pro^-eGFP-2 plasmid. 30 µg/mL polyethyleneimine was used for each transfection and the total of 8 μg of the plasmid in 500 μL of the opti-MEM medium. The cells were incubated with transfecting reagents overnight. On the second day, the medium was removed, cells were washed with a PBS buffer, and digested with 0.05% trypsin-EDTA, followed by resuspending the cells in the original growth media. The cell density was adjusted to 5 × 10^5^ cells/mL. 500 μL of suspended cells were provided in the growth media to each well of a 48-well plate, and then 100 μL of a drug solution was added in the growth media. These cells were then incubated under the aforementioned growth conditions with varying concentrations of drug 37°C and 5% CO^2^ for 72h before flow cytometry analysis. Following the drug incubation, the cells resuspended in 500 µL of PBS before being spun down at 800 rpm for 5 min spun down at 800 rpm for 5 min. The supernatant was removed, and the cell pellets were suspended in 200 µL of PBS. The fluorescence of each cell sample was collected by Cytoflex Beckman Flow Cytometer based on the size scatters (SSC-A and SSC-H) and forward scatter (FSC-A). The cells were gated based on SSC-A and FSC-A, then with SSC-A and SSC-H. The eGFP fluorescence was excited by blue laser (488 nm) and cells were collected at FITC-A (525 nm). All processed data were plotted and fitted to a four-parameter Hill equation in GraphPad 9.0 to obtain determined EC_50_ values.

### Live virus antiviral testing

SARS-CoV-2 delta variant hCoV-19/USA/MD-HP05647/2021 (BEI Resources, NR-55672) was propagated in A549-hACE2 cells (BEI Resources NR-53522) for antiviral testing, at 37ºC in an air-jacketed incubator, with 5% CO^2^ and >90% relative humidity, under BSL-3 conditions at the Texas A&M Global Health Research Complex. A low-dose, multi-step growth protocol was used for live virus EC_50_ assays. Briefly, A549-hACE2 cells were cultured in DMEM supplemented with 10% fetal bovine serum overnight. Approximately 5×10^4^ A549-hACE2 cells were inoculated by adding 10^3^ infectious units of SARS-CoV-2, as determined by tissue culture infectious dose 50% (TCID50) assay, and incubated at 37ºC for 1 hour. Cells were then aspirated and rinsed three times with room temperature phosphate buffered saline, to remove residual inoculum, before replacing DMEM with 10% FBS. Serial three-fold dilutions of candidate antivirals were made in DMEM with 10% FBS and added to three replicate wells per treatment condition. Infected, treated cells were then cultured for 72h at 37ºC, 5% CO^2^. At 48h, and 72h after inoculation, 50 microliters of tissue culture medium were removed from each sample, for quantitative SARS-CoV-2 reverse transcriptase quantitative PCR (RT-qPCR). After 72h, cell culture supernatant was removed by aspiration, cells were fixed in 10% formalin, 1× phosphate buffered saline for at least 30 min, then stained with crystal violet in order to qualitatively assess cytopathic effect. EC_50_ values were calculated from the slope and intercept of log-transformed RT-qPCR results, at the point where the linear portion of the transformed dose-response curve showed 50% reduced growth compared to infected, untreated controls. Samples were processed and RT-qPCR was performed as per protocol established previously.^54^ Samples were diluted 1:1 in 2× Tris-borate-EDTA [TBE] containing 1% Tween-20 and heated at 95°C for 15 min to lyse and inactivate virions. RT-qPCR screening was performed using the CDC N1 oligonucleotide pair/FAM probe (CDC N1-F, 5′-GACCCCAAAATCAGCGAAAT-3′; CDC N1-R, 5′-TCTGGTTACTGCCAGTTGAATCTG-3′; and Probe CDC N1, 5′ FAM-ACCCCGCATTACGTTTGGTGGACC-BHQ1 3′) and the Luna Universal Probe one-step RT-qPCR kit (catalog no. E3006; New England Biolabs). A 20-μL RT-qPCR mixture contained 7 μL of sample, 0.8 μL each of forward and reverse oligonucleotides (10 μM), 0.4 μL of probe (10 μM), and 11 μL of NEB Luna one-step RT-qPCR 2× master mix. Samples were incubated at 55°C for 10 min for cDNA synthesis, followed by 95°C for 1 min (1 cycle), then 41 cycles of 95°C for 10 s and 60°C for 30 s. Genome copies were quantitated relative to quantitative PCR control RNA from heat-inactivated SARS-Related Coronavirus 2, Isolate USA-WA1/2020 (BEI Resources, NR-52347).

### Cytotoxicity Assay of the Mpro Inhibitors

To assess the half-maximal cytotoxic concentration (CC^50^), stock solutions of the tested compounds were dissolved in DMSO (10mM) and diluted further to the working solutions with DMEM. HEK293T cells were seeded in 96 well-plates and incubated at 37°C and 5% CO^2^ for 24h. The cells were then treated with different concentrations (200 µM, 100 µM, 50 µM, 25 µM, 12.5 µM, 6.25 µM, 3.125 µM, 1.5625 µM, 0.78125 µM, 0 µM) of the tested compounds in triplicates for 48h. Cell viability was assessed by MTT assay to determine CC^50^. 20 µL MTT (5mg/mL) was incubated per well for 4h and then after removing supernatant, 200 µL DMSO was added per well. The absorbance was recorded at 490 nm to determine the CC^50^. The CC^50^ values were obtained by plotting the normalization % cell viability versus log^10^ sample concentration.

### *In Vitro* Metabolic Stability in Human Liver Microsomes

This metabolic stability measurements were based on previous publications and modified as described below.^55^ The metabolic stability profile of the inhibitor, including CL_int, pred_ and *in vitro* t_1/2_ was determined by the estimation of the remaining compound concentration after incubation with human liver microsome, NADPH (cofactor), EDTA, and MgCl^2^ in a 0.1 M phosphate buffer (pH 7.4). 5 μM of each inhibitor was preincubated with 40 μL of human liver microsome (0.5 mg/mL) in 0.1 M phosphate buffer (pH 7.4) at 37°C for 5 min to set optimal a condition for metabolic reactions. After pre incubation, NADPH (5 mM, 10 μL) or 0.1 M PB (10 μL) was added to initiate metabolic reaction at 37°C. The reactions were conducted in triplicate. At 0, 5, 15, 30, 45, 60 min, 200 μL acetonitrile (with internal standard Diclofenac, 10 µg/mL) was added in order to quench the reaction. The samples were subjected to centrifugation at 4°C for 20 min at 4000 rpm. Then 50 μL of clear supernatants were analyzed by HPLC-MS/MS. The percentage of test compound remaining was determined by following formula: % remaining = (Area at t_x_ /Average area at t^0^) × 100. The half-life (t_1/2_) was calculated using the slope (k) of the log-linear regression from the % remaining parent compound versus time (min) relationship: t_1/2_ (min) = –ln 2/k. CL_int, pred_ (mL/min/kg) was calculated through following formula CLi_nt, pred_ = (0.693/t_1/2_) × (1/ (microsomal protein concentration (0.5 mg/mL)) × Scaling Factor (1254.16 for human liver microsome).

### The Synthesis of MPIs

All reagents and solvents for the synthesis were purchased from commercial sources and used without purification. All glassware was flame-dried prior to use. Thin-layer chromatography (TLC) was carried out on aluminum plates coated with 60 F254 silica gel. TLC plates were visualized under UV light (254 or 365 nm) or stained with 5% phosphomolybdic acid. Normal phase column chromatography was carried out using a Yamazen Small Flash AKROS system. NMR spectra were recorded on a Bruker AVANCE Neo 400 MHz spectrometer in specified deuterated solvents. Analytical liquid chromatography-mass spectrometry was performed on a PHENOMENEX C18 Column (150 × 2.00 mm 5u micron, gradient from 10% to 100% B [A = 10 mmol/L HCOONH4 /H2O, B = MeOH], flow rate: 0.3 mL/min) using a Thermo Scientific Ultimate 3000 with a UV-detector (detection at 215 nm), equipped with a Thermo Scientific Orbitrap Q Exactive Focus System.

## ASSOCIATED CONTENT

### Supporting Information

The Supporting Information is available free of charge on the ACS Publications website.

Supplementary methods, figures, and tables (PDF)

## AUTHOR INFORMATION

### Author Contributions

K.K.‡, Y.R.A.‡, K.S.Y.‡, V.R.V.‡, L.R.B., D.D.C., S.A., S.P.C., Z.Z.G., X.R.M., J.X., P.-H.C.C., C.-C.D.C., E.C.V., Y.M., and G.Y. conducted the experiments and the refinement of crystal structures. K.K., B.W.N., S.X., and W.R.L. interpreted data and drafted the manuscript with input from all other authors. ‡These authors contributed equally.

### Notes

The authors declare no competing financial interest.

## ACKNOWLEDGMENT

This work was supported by the Welch Foundation (grant A-1715), National Institutes of Health (grants R35GM145351 to W.R.L., R21AI164088 to S.X., and R21EB032983 to W.R.L.), Texas A&M X Grants, and the Texas A&M EDGES Fellowship Program. While we regret any inadvertent omissions, we acknowledge the substantial body of existing literature on M^Pro^ inhibitor development. The authors thank Dr. Yohannes Rezenom, Mass Spectrometry Facility, TAMU, for helping with running the HPLC of the inhibitors.

## ABBREVIATIONS

COVID-19: coronavirus disease 2019
SARS-CoV-2: severe acute respiratory syndrome coronavirus 2
M^Pro^: main protease.

---

SYNOPSIS TOC (Word Style “SN_Synopsis_TOC”). If you are submitting your paper to a journal that requires a synopsis graphic and/or synopsis paragraph, see the Instructions for Authors on the journal’s homepage for a description of what needs to be provided and for the size requirements of the artwork.

To format double-column figures, schemes, charts, and tables, use the following instructions:

Place the insertion point where you want to change the number of columns

From the **Insert** menu, choose **Break**

Under **Sections**, choose **Continuous**

Make sure the insertion point is in the new section. From the **Format** menu, choose **Columns**

In the **Number of Columns** box, type **1**

Choose the **OK** button

Now your page is set up so that figures, schemes, charts, and tables can span two columns. These must appear at the top of the page. Be sure to add another section break after the table and change it back to two columns with a spacing of 0.33 in.

**Table 1.**
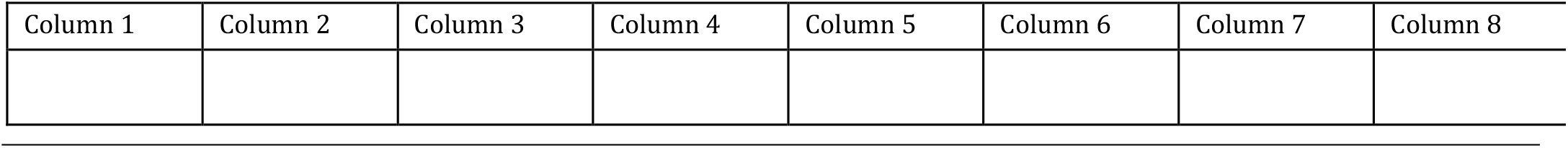
Example of a Double-Column Table.

Authors are required to submit a graphic entry for the Table of Contents (TOC) that, in conjunction with the manuscript title, should give the reader a representative idea of one of the following: A key structure, reaction, equation, concept, or theorem, etc., that is discussed in the manuscript. Consult the journal’s Instructions for Authors for TOC graphic specifications.

## Insert Table of Contents artwork here

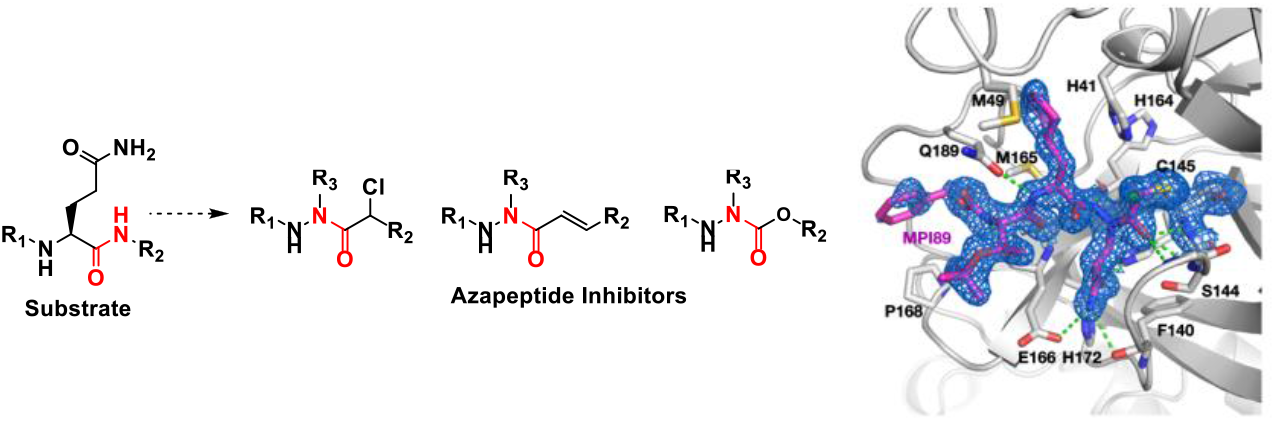

