## Supplemental information for "An Azapeptide Platform in Conjunction with Covalent Warheads to Uncover High-Potency Inhibitors for SARS-CoV-2 Main Protease"

### Supplementary Figures

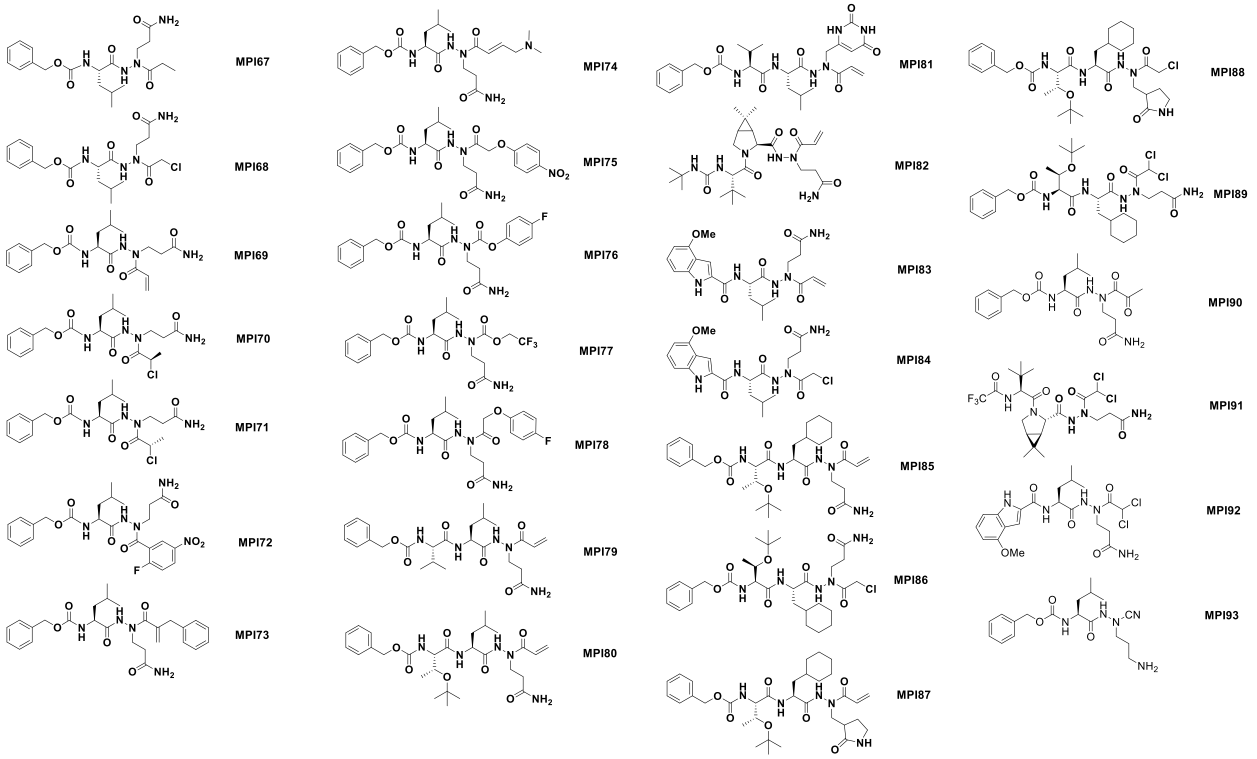


**Figure S1**. Structures of azapeptide M$^{Pro}$ inhibitors.

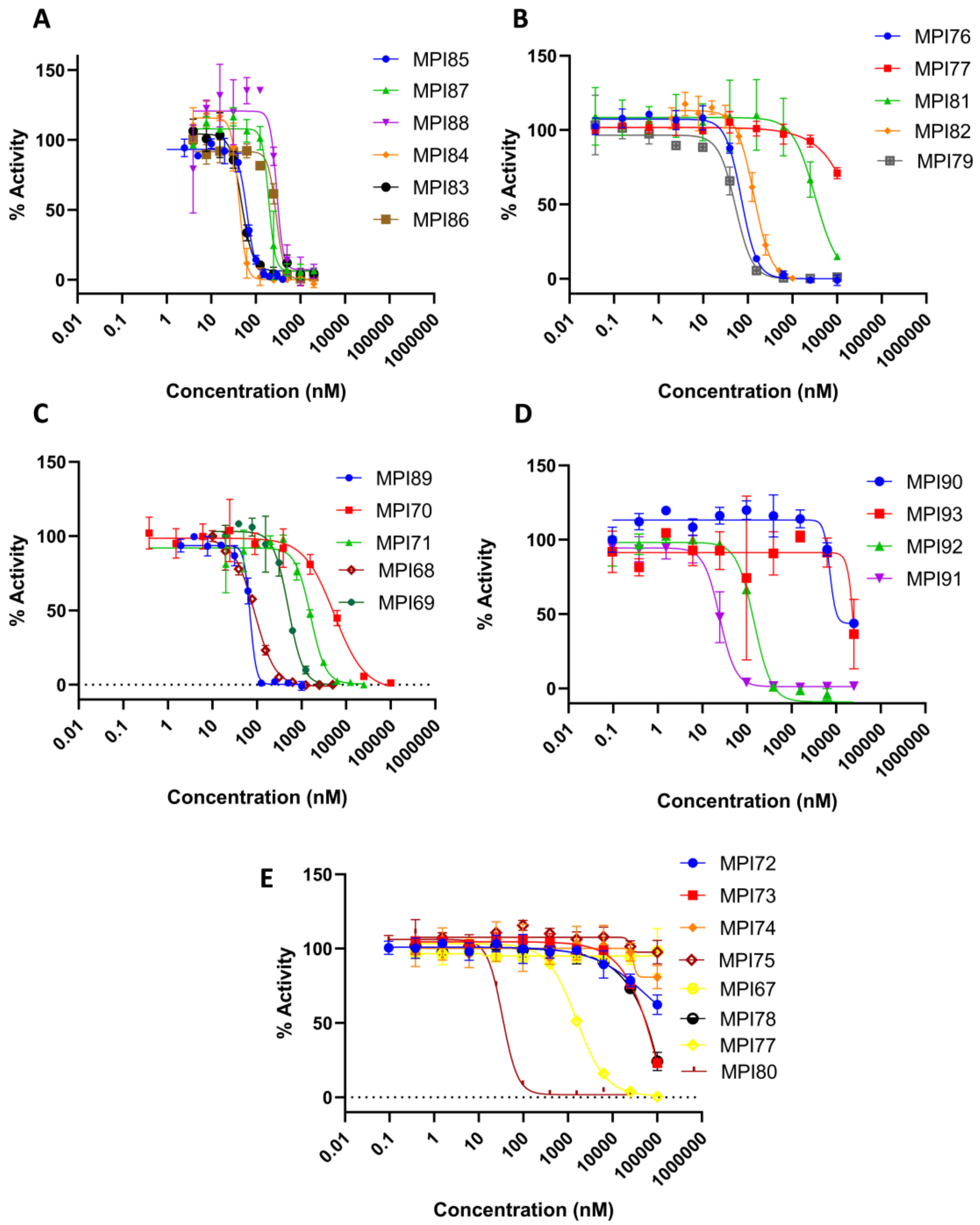


**Figure S2**. Inhibition curves of compounds on $M^{Pro}$. Triplicate experiments were performed for each compound. For all experiments, 20 $M^{Pro}$ was incubated with an inhibitor for 30 min before 10 μM Sub3 was added. The $M^{Pro}$-catalyzed Sub3 hydrolysis rate was determined by measuring linear increase of product fluorescence (Ex: 336 nm/Em: 455 nm) for 5 min.

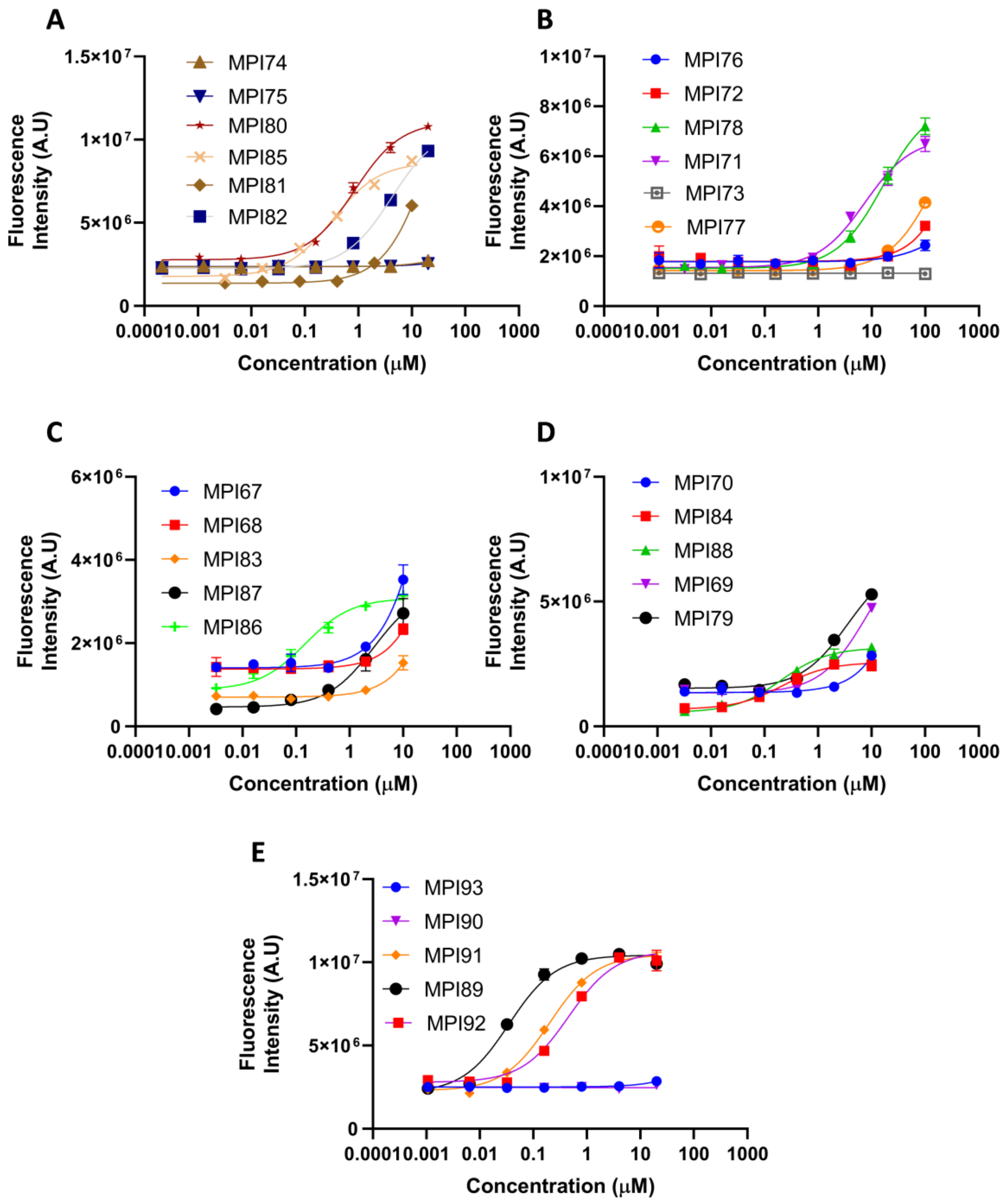


**Figure S3.** Cellular potency of inhibitors in their inhibition of $M^{Pro}$ to drive host 293T cell survival and overall $M^{Pro}$-eGFP expression.

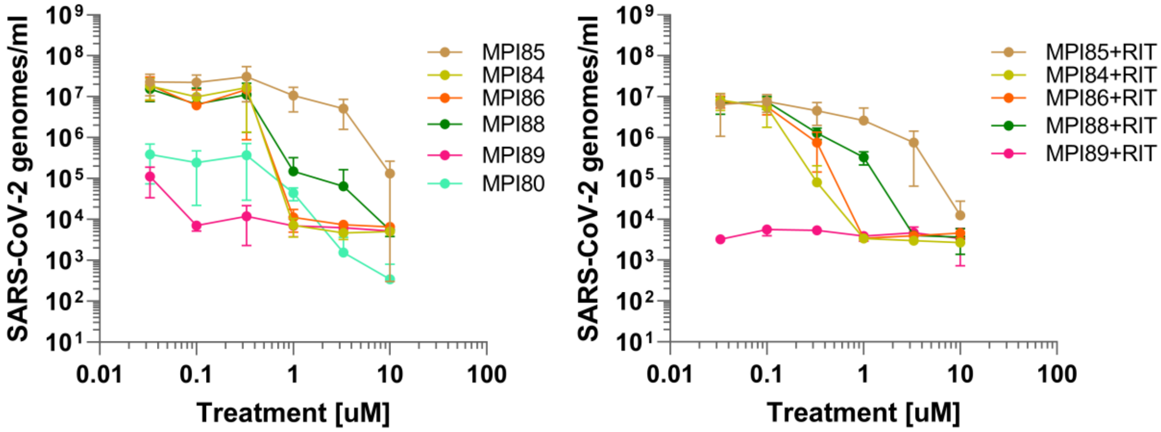


**Figure S4.** Antiviral potency tests for the inhibitors in ACE2+ A549 cells infected with hCoV-19/USA/HP05647/2021, an early Delta variant that grows well in cell culture and produces strong cytopathic effect.

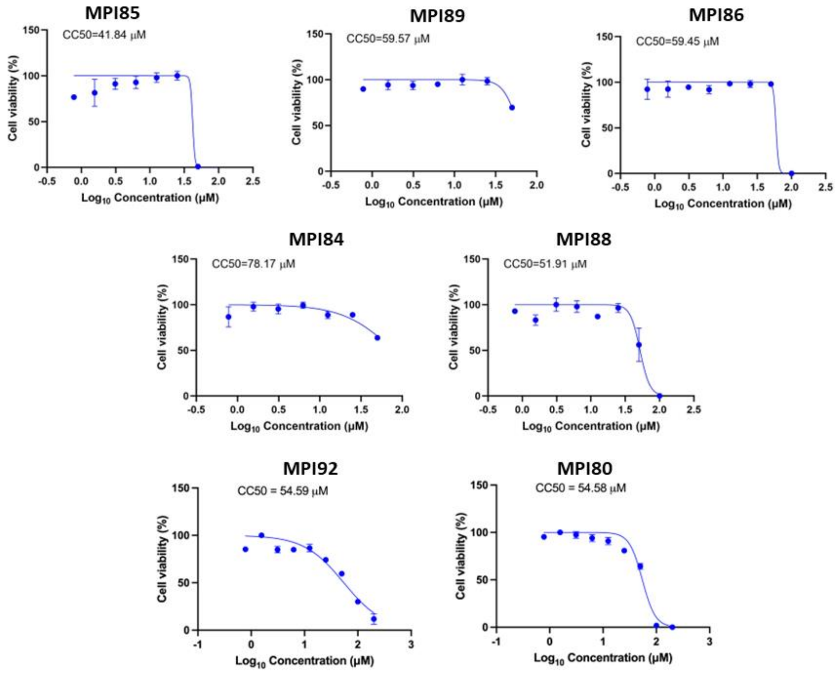


**Figure S5.** Cytotoxicity of the inhibitors in 293T cells using the MTT assay.

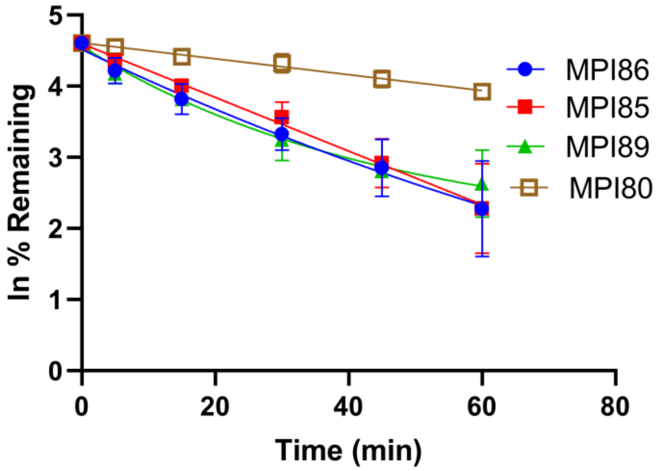


| Sample ID | Half life (min) | Intrinsic clearance (mL/min/kg) |
|---|---|---|
| MPI85 | 19.2 ± 4.8 | 94.7 ± 25.7 |
| MPI86 | 19.7 ± 5.2 | 92.1 ± 24.4 |
| MPI89 | 21.4 ± 3.4 | 82.43 ± 12 |
| MPI80 | 62.14 ± 5 | 28.1 ± 2.2 |

**Figure S6.** Stability of the inhibitors in human liver microsomes.

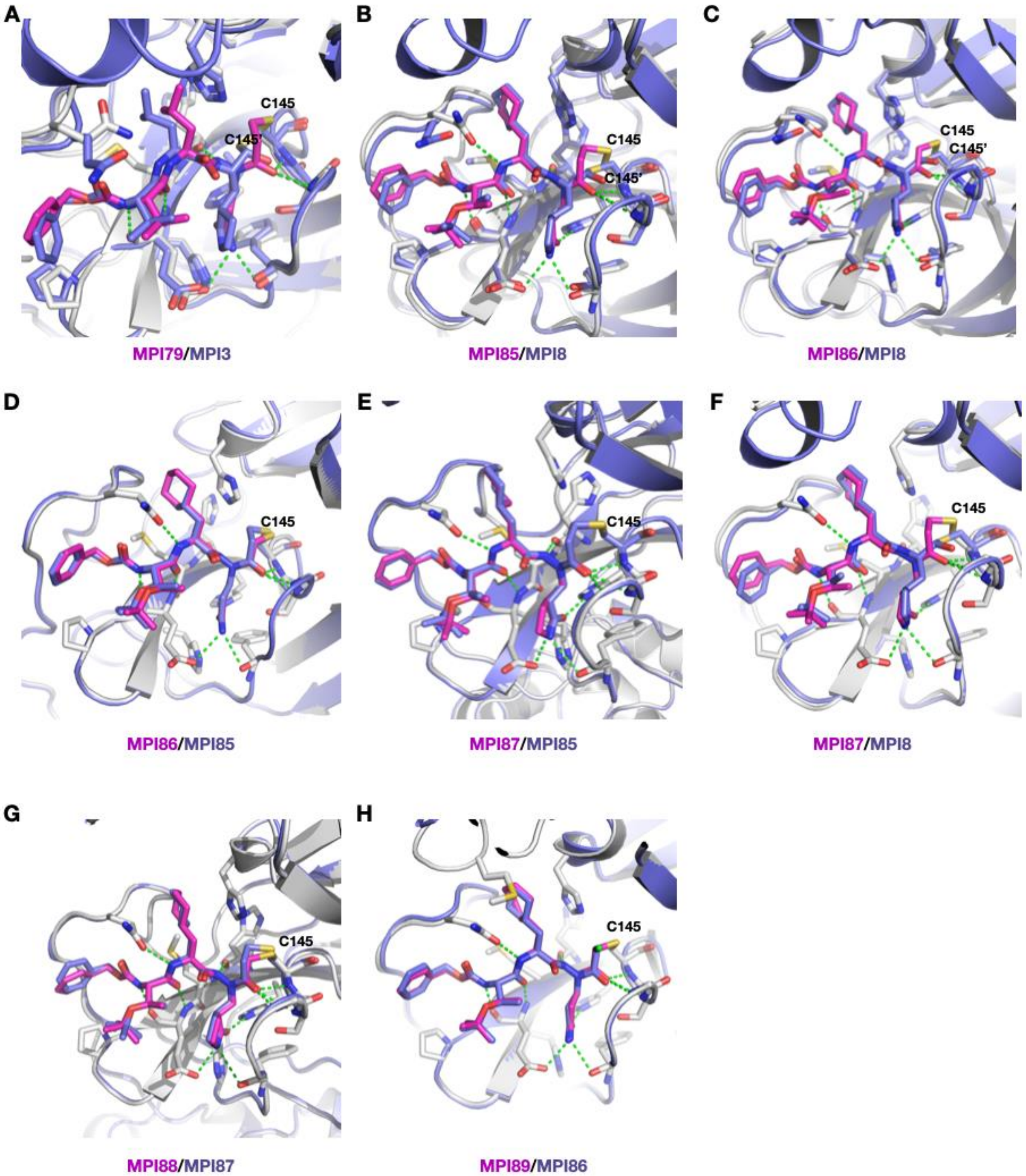


**Figure S7**: The structures superpositions of different M$^{Pro}$-inhibitor complexes. Ligands are colored according to their name colors shown in the figure. M$^{Pro}$-MPI3 is based on the pdb entry: 7JQ0 and M$^{Pro}$-MPI8 is based on the pdb entry: 7JQ5.

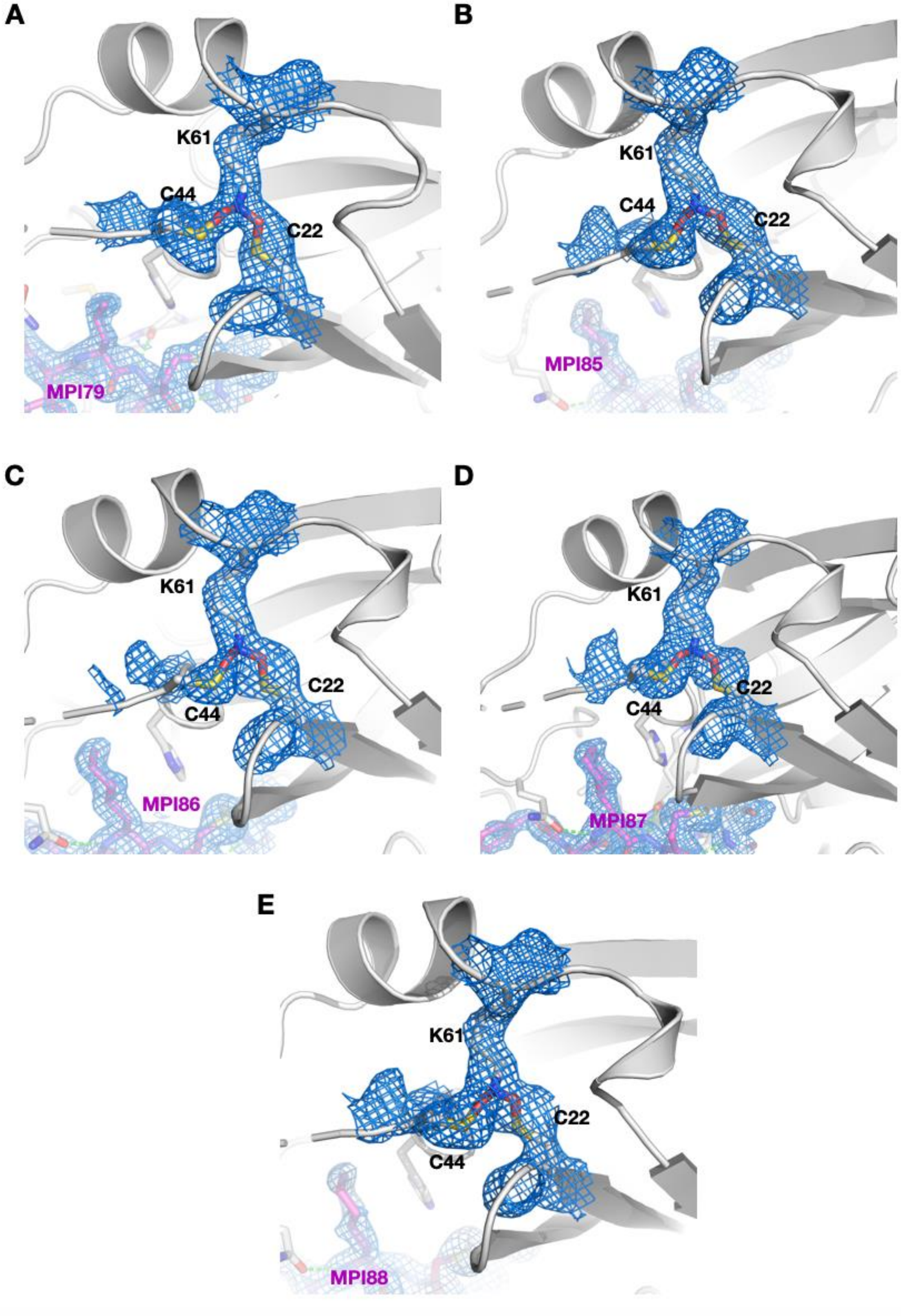


**Figure S8**: The S-O-N-O-S bridge between C22, C41, and Y61 that were observed in five M$^{Pro}$-azapeptide ligand complexes.

**Table S1.** Data Collection and Refinement Statistics

| Ligand (PDB Entry) | MPI79 (7SHB) | MPI85 (7SHA) | MPI86 (7SH9) | MPI87 (7SH7) | MPI88 (7SH8) | MPI89 (8S9Z) |
|---|---|---|---|---|---|---|
| Data Collection | | | | | | |
| Space Group | I 1 2 1 | I121 | I121 | I 1 2 1 | I 1 2 1 | I 1 2 1 |
| cell dimensions | | | | | | |
| *a, b, c* (Å) | 51.7395 81.4496 89.2227 | 54.6222 83.3566 85.8644 | 54.3812 82.7831 85.4791 | 54.2046 83.3917 86.0613 | 54.7038 83.4127 86.4164 | 54.2669 82.3178 86.4408 |
| $\alpha, \beta, \gamma$ (°) | 90 96.3983 90 | 90 96.1054 90 | 90 96.9608 90 | 90 96.7163 90 | 90 96.7048 90 | 90 96.1703 90 |
| Resolution Range (Å) | 24.3 - 2.452 (2.54 - 2.452) | 24.73-1.793 (1.832-1.793) | 24.57-1.59 (1.74-1.592) | 24.7 - 1.85 (1.916 - 1.85) | 24.18 - 1.8 (1.864 - 1.8) | 24.46 - 1.6 (1.657 - 1.6) |
| Unique Reflections | 12804 (865) | 31786 (3122) | 32008 (3200) | 31535 (3128) | 35544 (3514) | 48692 (4877) |
| Completeness (%) | 94.35 (63.92) | 97.4 (92.7) | 99.6 (94.2) | 96.94 (96.39) | 97.14 (96.40) | 97.73 (98.68) |
| Wilson B-factor | 15.79 | 15.6 | 10.1 | 17.25 | 22.45 | 15.28 |
| Refinement | | | | | | |
| No. Reflections | 12798 (861) | 66884 | 77993 | 31520 (3124) | 34728 (3429) | 48659 (4873) |
| No. Reflections *R*-free | 647 (32) | 3548 (277) | 2388 (271) | 1589 (189) | 1761 (193) | 2405 (224) |
| *R*-work | 0.2385 (0.3487) | 0.3163 (0.4348) | 0.3047 (0.4346) | 0.2795 (0.3827) | 0.2436 (0.3396) | 0.2398 (0.3995) |
| R-free | 0.2348 (0.3644) | 0.3121 (0.4410) | 0.3087 (0.4776) | 0.2740 (0.3897) | 0.2820 (0.3532) | 0.2591 (0.4047) |
| No. atoms | | | | | | |
| Non-Hydrogen | 2534 | 2631 | 2623 | 2652 | 2628 | 2723 |
| Macromolecules | 2367 | 2367 | 2367 | 2367 | 2367 | 2363 |
| Ligands | 36 | 43 | 42 | 45 | 44 | 43 |
| Water | 131 | 221 | 214 | 240 | 217 | 317 |
| Protein Reisues | 306 | 306 | 306 | 306 | 306 | 306 |
| RMS | | | | | | |
| Bond Length (Å) | 0.01 | 0.009 | 0.009 | 0.009 | 0.019 | 0.008 |
| Bond Angles (°) | 1.65 | 1.11 | 1.1 | 1.06 | 1.96 | 1.03 |

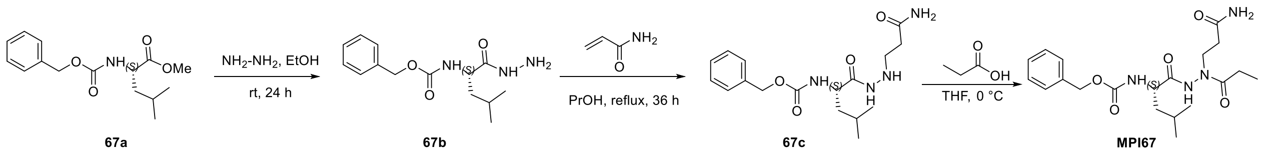


**Scheme 1:** The synthesis of compound **MPI67 (AzaPep)**

**Synthesis of (S)-benzyl (1-hydrazinyl-4-methyl-1-oxopentan-2-yl)carbamate (67b).** To a stirring solution of the compound **67a** (3.1 g, 11.0 mmol) in 15 mL of ethanol was added hydrazine hydrate (5.9 g, 118 mmol). The solution was stirred at room temperature for 16 hours then concentrated to yield compound **67b** as an off-white solid (3.1 g, 100%).

**Synthesis of (S)-benzyl (1-(2-(3-amino-3-oxopropyl)hydrazinyl)-4-methyl-1-oxopentan-2-yl)carbamate (67c).** Hydrazine derivative **67b** (2.0 g, 7.16 mmol) is dissolved in PrOH (3 mL), acrylamide (0.412 g, 7.16 mmol) added, and the solution refluxed for 36 h. The solution is cooled, filtered and the solvent removed in vacuum to give viscous oil. The oily compound was purified by chromatography column (Ethyl acetate: ethanol 9:1) to yield compound **67c** (400 mg, 16%) and 1 g of the starting material was recovered. $^{1}$H NMR (400 MHz, DMSO-$d_6$) δ 9.44 (d, $J$ = 6.2 Hz, 1H), 7.61 – 7.21 (m, 5H), 6.78 (s, 1H), 5.02 (s, 2H), 4.95 (q, $J$ = 6.0 Hz, 1H), 4.07 – 3.84 (m, 1H), 2.94 – 2.73 (m, 2H), 2.16 (t, $J$ = 7.0 Hz, 2H), 1.73 – 1.30 (m, 3H), 0.86 (dd, $J$ = 11.8, 6.6 Hz, 6H).$^{13}$C NMR (101 MHz, DMSO) δ 173.57, 171.35, 156.31, 137.55, 128.79, 128.23, 128.14, 65.81, 52.23, 47.90, 34.26, 24.67, 23.29, 22.12.

**Synthesis of (S)-benzyl (1-(2-(3-amino-3-oxopropyl)-2-propionylhydrazinyl)-4-methyl-1-oxopentan-2-yl)carbamate (MPI67, AzaPep).** To a solution **67c** (86 mg, 0.246 mmol) and propionic acid (22 mg, 0.295 mmol) were dissolved in dry DMF (3 mL) and the reaction was cooled to 0 °C. HATU (122 mg, 0.319 mmol) and DIPEA (0.174 mL, 0.983 mmol) were added, and the reaction mixture was allowed warm up to room temperature and stirred for 12 h. The mixture was then poured into water (10 mL) and extracted with ethyl acetate (4×20 mL). The organic layer was washed with aqueous hydrochloric acid 10% v/v (2×20 mL), saturated aqueous $NaHCO_3$ (2×20 mL), brine (2×20 mL) and dried over $Na_2SO_4$. The organic phase was evaporated to dryness and the crude material purified by silica gel column chromatography (1-10% MeOH in $CH_2Cl_2$ as the eluent) to afford **MPI67 (AzaPep)** white solid (52 mg, 52%). $^{1}$H NMR (400 MHz,

Chloroform-*d*) δ 9.86 (s, 1H), 7.47 – 7.21 (m, 5H), 6.94 (s, 1H), 6.09 (s, 1H), 5.89 (d, *J* = 7.9 Hz, 1H), 5.07 (s, 2H), 4.28 (q, *J* = 7.7 Hz, 1H), 3.76 (s, 2H), 2.60 – 2.34 (m, 2H), 2.30 – 2.12 (m, 2H), 1.75 – 1.54 (m, 3H), 1.04 – 0.85 (m, 9H). $^{13}$C NMR (101 MHz, $CDCl_3$): δ 176.42, 174.71, 172.59, 156.41, 136.06, 128.59, 128.33, 128.04, 67.21, 60.43, 52.45, 40.68, 33.73, 25.37, 24.73, 22.87, 21.76, 8.74. HRMS (APCI+) m/z calculated for C20H30N4O5 $^{+}$ $(M + H)^{+}$ 407.2289, found 407.2283.

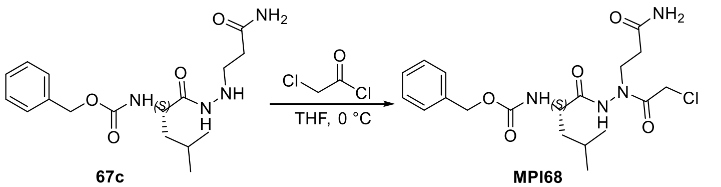


**Scheme 2**: The synthesis of **MPI68**

**Synthesis of (S)-benzyl (1-(2-(3-amino-3-oxopropyl)-2-(2-chloroacetyl)hydrazinyl)-4-methyl-1-oxopentan-2-yl)carbamate (MPI68).** To a stirred solution of **67c** (0.1 g, 0.285 mmol) in THF (5 mL) at 0 ◦C, chloroacetylchloride (22 μL, 0.314mmol) was added drop wise, and the mixture was stirred for further 30 min at same temperature. The reaction was quenched with water (5 mL), and the mixture was concentrated in vacuum. The residue was partitioned between EtOAc (30 mL) and $H_2O$ (20 mL). The aqueous layer was extracted with EtOAc (2 × 20 mL). The combined organic layer was washed with brine, dried over $MgSO_4$ and concentrated in vacuum. The residue was purified by flash column chromatography (0 to 6% MeOH in $CH_2Cl_2$ as the eluent)) to give the title compound **MPI68** as white solid (30 mg, 25%). $^{1}$H NMR (400 MHz, Chloroform-*d*) δ 9.43 (s, 1H), 7.49 – 7.29 (m, 5H), 6.12 (s, 1H), 5.76 (s, 1H), 5.33 (s, 1H), 5.12 (s, 2H), 4.29 – 4.15 (m, 1H), 4.11 – 3.75 (m, 4H), 2.65 – 2.43 (m, 2H), 1.80 – 1.51 (m, 3H), 1.09 – 0.82 (m, 6H). HRMS (ESI+) m/z calculated for C19H27ClN4O5 $^{+}$ $(M + H)^{+}$ 427.1743, found 427.1736.

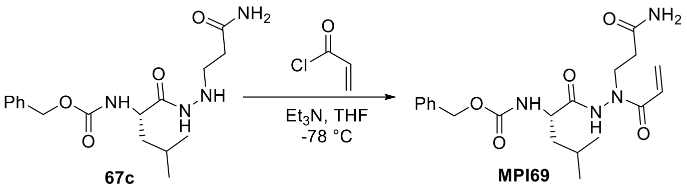


**Scheme 3**: The synthesis of **MPI69**

**Synthesis of (S)-benzyl (1-(2-acryloyl-2-(3-amino-3-oxopropyl)hydrazinyl)-4-methyl-1-oxopentan-2-yl)carbamate (MPI69).** To a stirred solution of **67c** (0.12 g, 0.342 mmol) in THF (5 mL) at −78 ◦C was added $Et_3N$ (960 µL, 0.685 mmol). After 15 min, acryloyl chloride (33µL, 0.41mmol) was added, and the mixture was stirred for further 30 min. The reaction was quenched with water (5 mL), and the mixture was concentrated in vacuum. The residue was partitioned between EtOAc (30 mL) and $H_2O$ (20 mL). The aqueous layer was extracted with EtOAc (2 × 20 mL). The combined organic layer was washed with brine, dried over $MgSO_4$ and concentrated in vacuum. The residue was purified by flash column chromatography (0 to 6% MeOH in $CH_2Cl_2$ as the eluent)) to give the title compound **MPI69** as white solid (55 mg, 40%).$^1$H NMR (400 MHz, Methanol-$d_4$) δ 7.37 – 7.06 (m, 5H), 6.51 (s, 1H), 6.17 (dd, $J$ = 17.0, 2.0 Hz, 1H), 5.60 (d, $J$ = 18.0 Hz, 1H), 5.02 (d, $J$ = 4.0 Hz, 2H), 4.07 (dd, $J$ = 9.8, 5.3 Hz, 1H), 3.91 – 3.49 (m, 2H), 2.51 – 2.26 (m, 2H), 1.70 – 1.41 (m, 3H), 0.87 (dd, $J$ = 13.1, 6.5 Hz, 6H). $^{13}$C NMR (101 MHz, MeOD) δ 174.72, 173.40, 168.17, 157.23, 136.85, 128.60, 128.16, 127.71, 127.53, 126.36, 66.39, 53.52, 39.83, 32.77, 24.50, 21.97, 20.23. HRMS (ESI+) m/z calculated for C20H28N4O5 $^+$ (M + H)$^+$ 405.2132, found 405.2122.

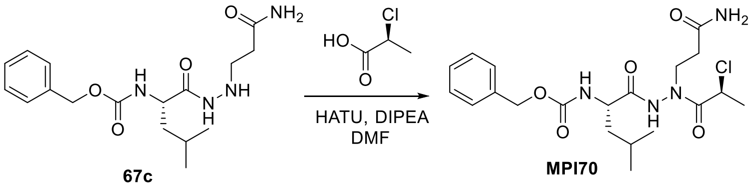

**Scheme 4**: The synthesis of **MPI70**

**Synthesis of benzyl ((S)-1-(2-(3-amino-3-oxopropyl)-2-((S)-2-chloropropanoyl)hydrazineyl)-4-methyl-1-oxopentan-2-yl)carbamate (MPI70).** To a solution of **67c** (70 mg, 0.2 mmol) in DMF (1 mL) was added (S)-2-chloropropanoic acid (27 mg, 0.25 mmol) and DIPEA (26 mg, 0.2 mmol). The solution was cooled at 0 °C and HATU (95 mg, 0.25 mmol) was added to the solution. The reaction mixture was then allowed to warm up to room temperature and stirred at room temperature overnight. The reaction mixture was the diluted with ethyl acetate (10 mL), washed with saturated $NaHCO_3$ solution (2×10 mL), 1 M HCl (2×10 mL) and saturated brine (10 mL) sequentially. The organic layers were then dried over anhydrous $Na_2SO_4$, concentrated *in vacuo* and the residue was purified with flash chromatography (1-10% methanol in dichloromethane as the eluent) to afford **MPI70** as white solid. $^1$H NMR (400 MHz, Chloroform-*d*) δ 9.70 (s, 1H), 7.42 – 7.28 (m, 5H), 6.45 – 6.24 (m, 1H), 5.83 (s, 1H), 5.65 – 5.42 (m, 1H), 5.11 (s, 2H), 4.56 (s, 1H), 4.34 – 4.21 (m, 1H), 4.01 (s, 1H), 3.55 (s, 1H), 2.56 (s, 1H), 2.46 (s, 1H), 1.95 (s, 1H), 1.81 – 1.60 (m, 3H), 1.54 (d, *J* = 6.6 Hz, 3H), 1.11 – 0.84 (m, 6H). HRMS (ESI+) m/z calculated for C20H29ClN4O5 $^+$ $(M + H)^+$ 441.1899, found 441.1890.

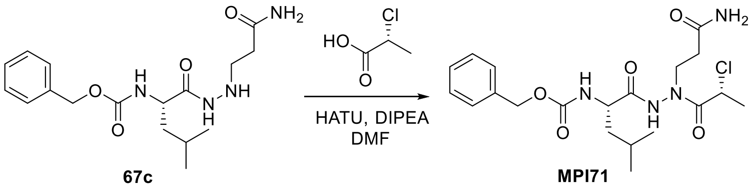

**Scheme 5**: The synthesis of **MPI71**

**Synthesis of benzyl ((S)-1-(2-(3-amino-3-oxopropyl)-2-((R)-2-chloropropanoyl)hydrazineyl)-4-methyl-1-oxopentan-2-yl)carbamate (MPI71).** To a solution of **67c** (70 mg, 0.2 mmol) in DMF (1 mL) was added (R)-2-chloropropanoic acid (27 mg, 0.25 mmol) and DIPEA (26 mg, 0.2 mmol). The solution was cooled at 0 °C and HATU (95 mg, 0.25 mmol) was added to the solution.

The reaction mixture was then allowed to warm up to room temperature and stirred at room temperature overnight. The reaction mixture was the diluted with ethyl acetate (10 mL), washed with saturated $NaHCO_3$ solution (2×10 mL), 1 M HCl (2×10 mL) and saturated brine (10 mL) sequentially. The organic layers were then dried over anhydrous $Na_2SO_4$, concentrated *in vacuo* and the residue was purified with flash chromatography (1-10% methanol in dichloromethane as the eluent) to afford **MPI71** as white solid. $^1$H NMR (400 MHz, Chloroform-*d*) δ 9.51 (d, *J* = 70.4 Hz, 1H), 7.41 – 7.30 (m, 5H), 6.15 (d, *J* = 44.0 Hz, 1H), 5.72 (s, 1H), 5.60 – 5.42 (m, 1H), 5.12 (s, 2H), 4.52 (s, 1H), 4.26 (d, *J* = 10.2 Hz, 1H), 4.17 – 3.90 (m, 1H), 3.52 (d, *J* = 33.7 Hz, 1H), 2.56 (s, 1H), 2.47 (s, 1H), 1.84 – 1.52 (m, 7H), 1.05 – 0.81 (m, 6H).

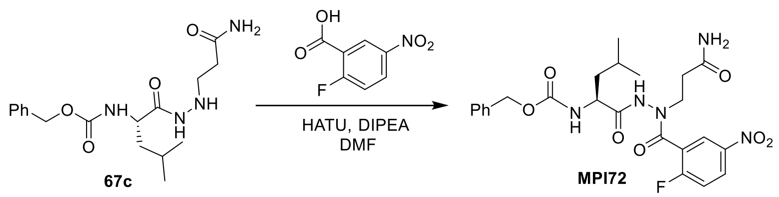


**Scheme 6**: The synthesis of **MPI72**

**Synthesis of benzyl (S)-(1-(2-(3-amino-3-oxopropyl)-2-(2-fluoro-5-nitrobenzoyl)hydrazineyl)-4-methyl-1-oxopentan-2-yl)carbamate (MPI72).** To a solution of **67c** (58 mg, 0.31 mmol, 1.0 equiv) in anhydrous DMF (25 mL) at 0 °C, and then 2-Fluoro-5-nitrobenzoic acid (120 mg, 0.35 mmol, 1.1 equiv), HATU (160 mg, 0.4 mmol, 1.3 equiv), DIPEA (0.25 mL, 1.3 mmol, 4.0 equiv) was added sequentially. The mixture was stirred at RT overnight. The mixture was diluted with EtOAc and washed with water, 1M HCl, sat. NaCl, dried over $Na_2SO_4$, and concentrated. The residue was purified by column chromatography (Hexane: EA = 3:1 v/v) to afford the pure product **MPI72** as a colorless oil. $^1$H NMR (400 MHz, Chloroform-*d*) δ 9.39 (s, 1H), 8.34 – 8.24 (m, 1H), 8.12 (s, 1H), 7.35 – 7.21 (m, 5H), 7.04 (s, 1H), 5.88 (s, 1H), 5.67 (s, 1H), 4.96 (s, 3H), 4.05 (q, *J* = 7.1 Hz, 1H), 3.90 (d, *J* = 7.3 Hz, 1H), 2.59 (s, 2H), 1.19 (t, *J* = 7.1 Hz, 4H), 0.67 (s, 6H). HRMS (ESI+) m/z calculated for C24H28FN5O7 $^+$ $(M + H)^+$ 518.2046, found 518.2038.

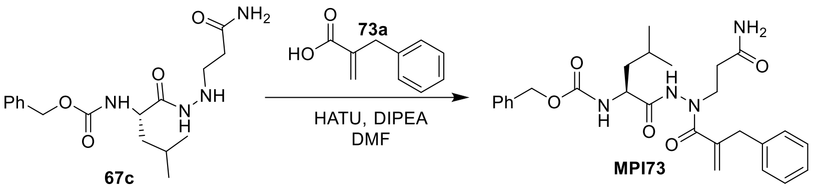


**Scheme 7**: The synthesis of **MPI73**

**Synthesis of benzyl (S)-(1-(2-(3-amino-3-oxopropyl)-2-(2-benzylacryloyl)hydrazineyl)-4-methyl-1-oxopentan-2-yl)carbamate (MPI73).** To a solution of **67c** (50 mg, 0.14 mmol, 1.0 equiv) in anhydrous DMF (25 mL) at 0 °C, and then **73a** (26 mg, 0.16 mmol, 1.1 equiv), HATU (71 mg, 0.18 mmol, 1.3 equiv), DIPEA (0.1 mL, 0.57 mmol, 4.0 equiv) was added sequentially. The mixture was stirred at RT overnight. The mixture was diluted with EtOAc and washed with water, 1M HCl, saturated NaCl, dried over $Na_2SO_4$, and concentrated. The residue was purified by column chromatography (DCM: methanol = 10:1 v/v) to afford the pure product **MPI73** as a colorless oil. $^1$H NMR (400 MHz, Chloroform-*d*) δ 8.63 (d, *J* = 20.8 Hz, 1H), 7.27 (d, *J* = 5.0 Hz, 6H), 7.17 – 7.02 (m, 3H), 6.12 (s, 1H), 5.58 (s, 1H), 5.30 – 4.72 (m, 4H), 4.16 – 3.96 (m, 1H), 3.65 (d, *J* = 52.9 Hz, 1H), 3.48 (s, 1H), 2.38 (d, *J* = 42.3 Hz, 2H), 1.74 – 1.26 (m, 4H), 1.00 – 0.70 (m, 6H). HRMS (ESI+) m/z calculated for C27H34N4O5 $^+$ (M + H)$^+$ 495.2602, found 495.2603.

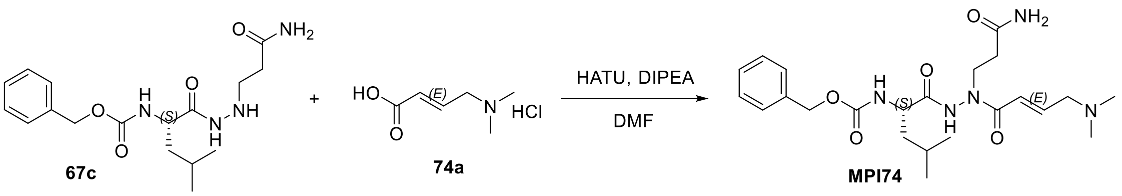


**Scheme 8**: The synthesis of **MPI74**

**Synthesis of benzyl (S,E)-(1-(2-(3-amino-3-oxopropyl)-2-(4-(dimethylamino)but-2-enoyl)hydrazineyl)-4-methyl-1-oxopentan-2-yl)carbamate (MPI74)**. To a stirred solution of **67c** (50 mg, 0.143 mmol) and the acid **74a** (28 mg, 0.171 mmol) were dissolved in dry DMF (3 mL) and the reaction was cooled to 0 °C. HATU (71 mg, 0.186 mmol) and DIPEA (0.1 mL, 0.571 mmol) were added, and the reaction mixture was allowed warm up to room temperature and stirred for 12 h. The mixture was then poured into water (10 mL) and extracted with ethyl acetate (4×20 mL). The organic layer was washed with aqueous hydrochloric acid 10% v/v (2×5 mL), saturated aqueous $NaHCO_3$ (2×5 mL), brine (2×5 mL) and dried over $Na_2SO_4$. The organic phase was evaporated to dryness and the crude material purified by silica gel column chromatography (1-8% MeOH in $CH_2Cl_2$ as the eluent) to afford **MPI74**. The LC-MS analysis of the purified compound showed two close peaks with same mass corresponding to **MPI74**, which could not be separated by column chromatography. HRMS (ESI+) m/z calculated for C23H35N5O5 $^+$ (M + H) $^+$ 462.2711, found 462.2703.

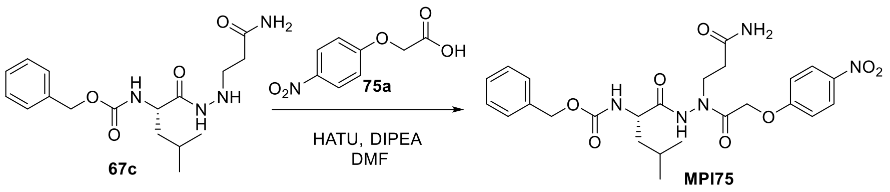


**Scheme 9**: The synthesis of **MPI75**

**(S)-benzyl (1-(2-(3-amino-3-oxopropyl)-2-(2-(4-nitrophenoxy)acetyl)hydrazinyl)-4-methyl-1-oxopentan-2-yl)carbamate (MPI75).** To a stirred solution of **67c** (50 mg, 0.143 mmol) and the acid **75a** (34 mg, 0.171 mmol) were dissolved in dry DMF (3 mL) and the reaction was cooled to 0 °C. HATU (71 mg, 0.186 mmol) and DIPEA (0.1 mL, 0.571 mmol) were added, and the reaction mixture was allowed warm up to room temperature and stirred for 12 h. The mixture was then poured into water (10 mL) and extracted with ethyl acetate (4×20 mL). The organic layer was washed with aqueous hydrochloric acid 10% v/v (2×5 mL), saturated aqueous $NaHCO_3$ (2×5 mL), brine (2×5 mL) and dried over $Na_2SO_4$. The organic phase was evaporated to dryness and the crude material purified by silica gel column chromatography (1-8% MeOH in $CH_2Cl_2$ as the eluent) to afford **MPI75** white solid (15 mg, 20%).$^1$H NMR (400 MHz, Chloroform-*d*) δ 9.31 (s, 1H), 8.16 (d, *J* = 9.2 Hz, 2H), 7.42 – 7.27 (m, 5H), 6.98 (d, *J* = 8.7 Hz, 2H), 5.72 (s, 1H), 5.46 (s, 1H), 5.23 – 5.08 (m, 3H), 4.71 (s, 2H), 4.30 – 4.18 (m, 1H), 3.67 – 3.25 (m, 2H), 2.59 (s, 2H), 1.78 – 1.61

(m, 3H), 1.08 – 0.92 (m, 6H). HRMS (ESI+) m/z calculated for C25H31N5O8 $^+$ (M + H) $^+$ 530.2245, found 530.2233.

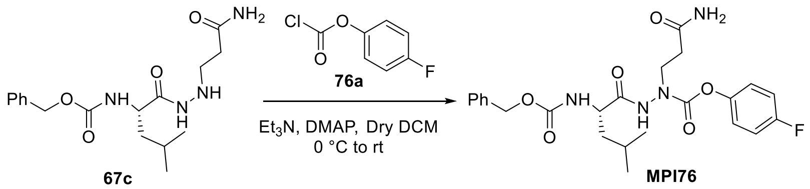

**Scheme 10**: The synthesis of **MPI76**

**(S)-4-fluorophenyl1-(3-amino-3-oxopropyl)-2-(2-(((benzyloxy)carbonyl)amino)-4-methylpentanoyl)hydrazinecarboxylate (MPI76).** To a stirred solution of **67c** (80 mg, 0.228 mmol) in dry $CH_2Cl_2$ (5 mL) at 0 ◦C were added $Et_3N$ (960 µL, 0.342 mmol) and DMAP (cat.). After 15 min, compound **76a** (33µL, 0.251mmol) was added, and the mixture was stirred at room temperature up to completion of starting material (by TLC conformation). The reaction was quenched with water (5 mL), and the mixture was concentrated in vacuum. The residue was partitioned between $CH_2Cl_2$ (30 mL) and $H_2O$ (10 mL). The aqueous layer was extracted with $CH_2Cl_2$ (2 × 20 mL). The combined organic layer was washed with brine, dried over $MgSO_4$ and concentrated in vacuum. The residue was purified by flash column chromatography (0 to 6% MeOH in $CH_2Cl_2$ as the eluent)) to give the title compound **MPI76** as white solid (40 mg, 36%)$^1$H NMR (400 MHz, DMSO-$d_6$) δ 9.39 (s, 1H), 7.95 (d, *J* = 8.3 Hz, 1H), 7.53 – 7.29 (m, 6H), 7.10 – 6.88 (m, 3H), 6.86 – 6.68 (m, 2H), 5.06 (d, *J* = 1.9 Hz, 2H), 4.57 (dt, *J* = 13.7, 7.0 Hz, 1H), 3.81 (t, *J* = 7.1 Hz, 2H), 2.47 (t, *J* = 7.2 Hz, 2H), 1.71 – 1.54 (m, 3H), 0.88 (dd, *J* = 11.2, 5.7 Hz, 6H). $^{13}$C NMR (101 MHz, DMSO) δ 171.58, 157.11, 156.25, 155.96, 154.80, 154.06, 154.04, 153.33, 137.20, 128.86, 128.40, 128.25, 116.57, 116.49, 116.19, 115.96, 66.24, 46.43, 42.08, 33.55, 24.37, 23.01, 21.85. HRMS (APCI+) m/z calculated for C24H29FN4O6 $^+$ (M + H) $^+$ 489.2144, found 489.2139.

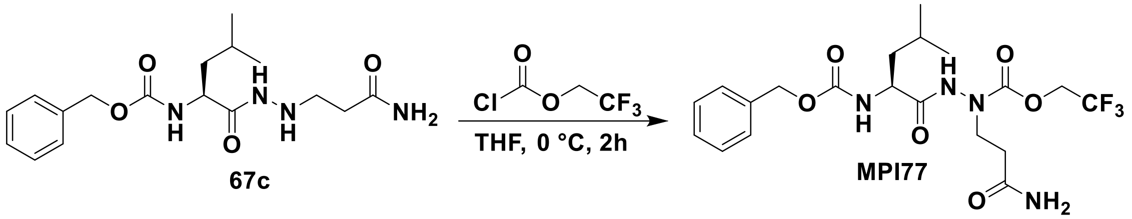

**Scheme 11**: The synthesis of **MPI77**

**2,2,2-trifluoroethyl 1-(3-amino-3-oxopropyl)-2-(((benzyloxy)carbonyl)-*L*-leucyl)hydrazine-1-carboxylate (MPI77).** To a stirred solution of **4** (50 mg, 0.14 mmol) in THF (5 mL) at 0 °C, 2,2,2-trifluoroethyl chloroformate (0.027 g, 0.17) was added dropwise, and the mixture was stirred for further 2 h at the room temperature. The reaction mixture is concentrated in a vacuum. The residue was purified by flash column chromatography (0 to 8% MeOH in $CH_2Cl_2$ as the eluent)) to give the title compound **6** as white solid (30 mg). $^1H$ NMR (400 MHz, DMSO-$d_6$) δ 10.49 (s, 1H), 7.61 (d, *J* = 7.8 Hz, 1H), 7.49-7.31 (m, 6H), 6.95 (d, *J* = 17.1 Hz, 1H), 5.14 – 5.03 (m, 2H), 4.90-4.58 (m, 1H), 4.11 (d, *J* = 10.1 Hz, 1H), 2.39 (t, *J* = 8.8, 4.3 Hz, 2H), 1.80-1.43 (m, 3H), 1.38-1.22 (m, 2H), 0.99 – 0.85 (m, 6H).

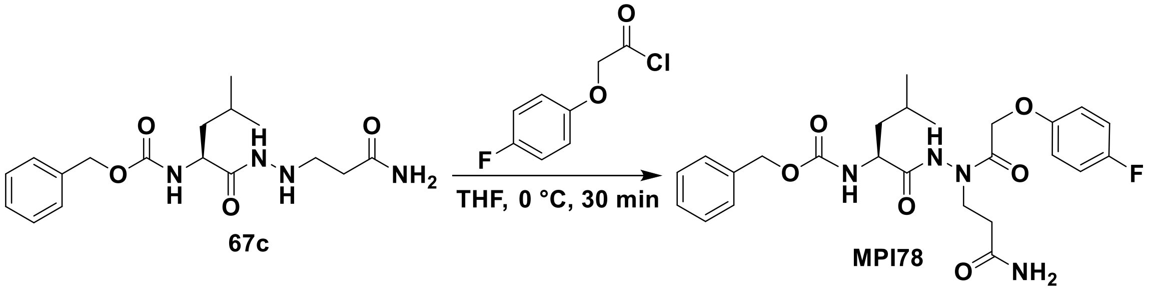

**Scheme 12**: The synthesis of **MPI78**

**Benzyl (*S*)-(1-(2-(3-amino-3-oxopropyl)-2-(2-(4-fluorophenoxy)acetyl)hydrazineyl)-4-methyl-1-oxopentan-2-yl)carbamate (MPI78).** To a stirred solution of 4 (50 mg, 0.085 mmol) in THF (5 mL) at 0 °C, 2-(4-fluorophenoxy)acetyl chloride (0.032 g, 0.17) was added dropwise, and the mixture was stirred for further 30 min at the same temperature. The reaction was quenched with water (5 mL), and the mixture was concentrated in a vacuum. The residue was partitioned between EtOAc (30 mL) and $H_2O$ (20 mL). The aqueous layer was extracted with EtOAc (2 × 20 mL). The combined organic layer was washed with brine, dried over $MgSO_4$, and concentrated in a vacuum. The residue was purified by flash column chromatography (0 to 6% MeOH in $CH_2Cl_2$ as the eluent)) to give the title compound 6 as white solid (30 mg). $^1$H NMR (400 MHz, DMSO-*d*$_6$) δ 10.73 (s, 1H), 7.52-7.05 (m, 8H), 7.45 (s, 1H), 6.97-6.8 (m, 2H), 5.12 – 5.00 (m, 2H), 4.76-4.40 (m, 2H), 4.20-4.08 (m, 1H), 3.94-3.75 (m, 1H), 2.47-2.37 (m, 2H), 1.80-1.54 (m, *J* = 13.0, 2H), 1.40-1.27(m, 2H), 1.01 – 0.75 (m, 6H).

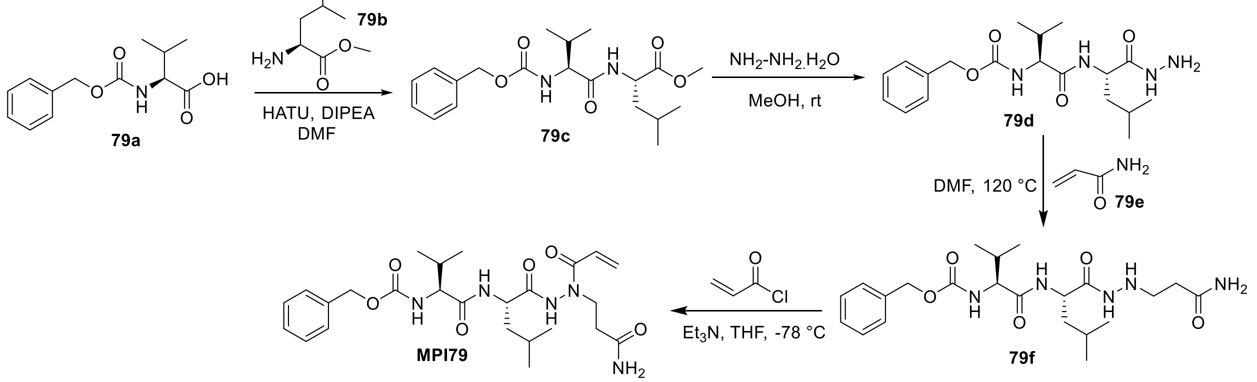


**Scheme 13**: The synthesis of **MPI79**

**Synthesis of (S)-methyl 2-((S)-2-(((benzyloxy)carbonyl)amino)-3-methylbutanamido)-4-methylpentanoate (79c).** To a solution of compound **79a** (1.0 g, 5.52 mmol) and compound **79b** (1.88 g, 6.08 mmol) in dry DMF (20 mL), was added HATU (2.52 g, 6.62 mmol) and DIPEA (3.92 mL, 22.08 mmol) and the reaction was cooled to 0 °C. The reaction mixture was allowed warm up to room temperature and stirred for 12 h. The mixture was then poured into water (50 mL) and extracted with ethyl acetate (4×20 mL). The organic layer was washed with aqueous hydrochloric acid 10% v/v (2×20 mL), saturated aqueous $NaHCO_3$ (2×20 mL), brine (2×20 mL) and dried over $Na_2SO_4$. The organic phase was evaporated to dryness and the crude material purified by silica gel column chromatography (15-50% EtOAc in n-hexane as the eluent) to afford **79c** white solid (1.82, 69%). $^1$H NMR ($CDCl_3$, 400MHz) δ 7.28-7.22 (m, 5H), 6,28 (d, *J*=7.72 Hz, 1H), 5.35 (d, =864 Hz, 1H), 5.03 (s, 2H), 4.56-451 (m, 1H), 3.97 (t, *J*=8.12 Hz, 1H), 3.65 (s, 3H), 2.19-1.98 (m, 1H),

1.62-1.43 (m, 3H), 0.92-0.83 (m, 12H); $^{13}$C NMR ($CDCl_3$, 100MHz) δ 173.2, 171.1, 156., 136.2, 128.5 (2C), 128.2, 128 (2C), 67.0, 60.2, 52.3, 50.7, 1., 31.3, 2.8, 22.7, 21.9, 19.1, 17.8.

**Synthesis of benzyl ((S)-1-(((S)-1-hydrazinyl-4-methyl-1-oxopentan-2-yl)amino)-3-methyl-1-oxobutan-2-yl)carbamate (79d).** To a stirred solution of compound **79c** (800 mg, 2.11 mmol) in ethanol (20 mL) at room temperature was added $N_2H_4.H_2O$ (80%, 8.46 mmol). The reaction mixture was stirred for 24 h at room temperature and concentrated under reduced pressure. The residue was purified by chromatography column (dichloromethane⁄methanol 9.5:0.5) to afford pure **79d**. $^{1}$H NMR (400 MHz, Methanol-$d_4$) δ 7.52 – 7.26 (m, 5H), 5.13 (s, 2H), 4.42 (dd, $J$ = 9.3, 5.7 Hz, 1H), 3.95 (d, $J$ = 7.0 Hz, 1H), 2.15 – 2.01 (m, 1H), 1.71 – 1.52 (m, 3H), 1.01 – 0.89 (m, 12H).

**Synthesis of benzyl ((S)-1-(((S)-1-(2-(3-amino-3-oxopropyl)hydrazinyl)-4-methyl-1-oxopentan-2-yl)amino)-3-methyl-1-oxobutan-2-yl)carbamate (79f).** To a solution of compound **79d** (800 mg, 2.11 mmol) in dissolved in PrOH (1.5 mL), was added acrylamide (0.15 g, 2.11 mmol) and the solution refluxed for 36 h. The solution was cooled, filtered and the solvent removed in vacuum to give viscous oil. The oily compound was purified by chromatography column (Ethyl acetate⁄methanol 9:1). (Yield 80 mg, 9%) and 500 mg of the starting material was recovered .$^{1}$H NMR (400 MHz, Methanol-$d_4$) δ 7.45 – 7.13 (m, 5H), 5.00 (s, 2H), 4.25 (dd, $J$ = 9.3, 5.5 Hz, 1H), 3.82 (dd, $J$ = 6.9, 3.6 Hz, 1H), 3.03 – 2.83 (m, 2H), 2.34 – 2.17 (m, 2H), 2.03 – 1.88 (m, 1H), 1.71 – 1.40 (m, 3H), 0.92 – 0.76 (m, 12H).

**Synthesis of Benzyl ((S)-1-(((S)-1-(2-acryloyl-2-(3-amino-3-oxopropyl)hydrazinyl)-4-methyl-1-oxopentan-2-yl)amino)-3-methyl-1-oxobutan-2-yl)carbamate (MPI79).** To a stirred solution of **79f** (40 g, 0.089 mmol) in THF (5 mL) at −78 ◦C, was added $Et_3N$ (25 µL, 0.178 mmol). After 15 min, acryloyl chloride (9 µL, 0.106 mmol) was added, and the mixture was stirred for further 30 min. The reaction was quenched with water (5 mL), and the mixture was concentrated in vacuum. The residue was partitioned between EtOAc (20 mL) and $H_2O$ (10 mL). The aqueous layer was extracted with EtOAc (2 × 20 mL). The combined organic layer was washed with brine, dried over $MgSO_4$ and concentrated in vacuum. The residue was purified by flash column chromatography (MeOH: $CH_2Cl_2$ 0.5:9.5) to give the title compound as a white solid (25 mg, 56%). $^{1}$H NMR (400 MHz, Methanol-$d_4$) δ 7.55 – 7.19 (m, 5H), 6.67 (s, 1H), 6.31 (dd, $J$ = 16.9, 2.1 Hz, 1H), 5.74 (d, $J$ = 10.3 Hz, 1H), 5.12 (s, 2H), 4.42 (dd, $J$ = 9.8, 5.3 Hz, 1H), 3.98 (d, $J$ = 7.0 Hz, 2H), 3.74 (s, 1H), 2.52 (s, 2H), 2.19 – 2.04 (m, 1H), 1.86 – 1.57 (m, 3H), 1.13 – 0.90 (m, 12H).

$^{13}C$ NMR (101 MHz, MeOD): δ 174.66, 173.30, 172.66, 168.19, 157.91, 128.10, 127.66, 127.50, 66.41, 60.89, 50.53, 39.48, 32.65, 30.51, 24.48, 21.89, 20.50, 18.35, 17.23. HRMS (ESI+) m/z calculated for C25H37N5O6 $^{+}$ (M + H) $^{+}$ 504.2817, found 504.2805.

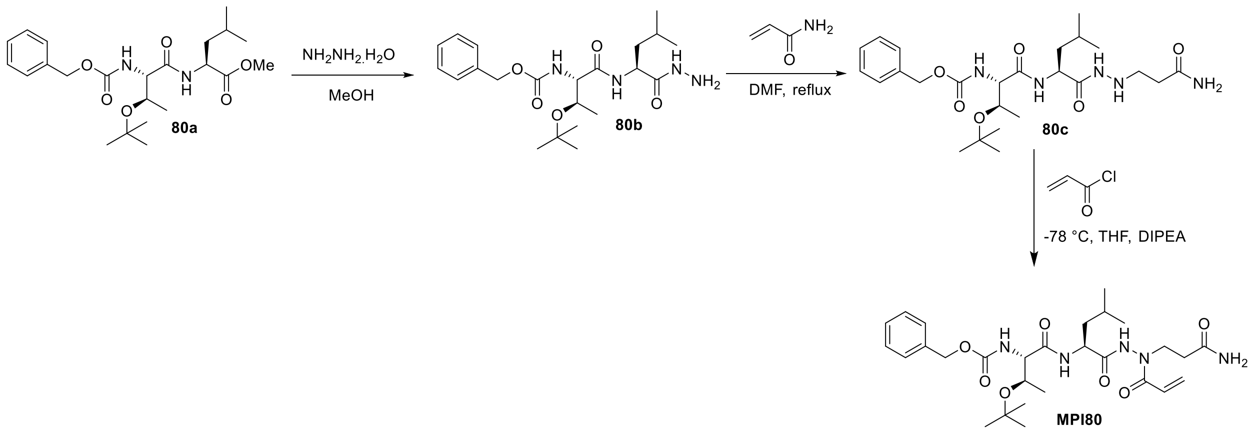


**Scheme 14**: The synthesis of **MPI80**

**Synthesis of benzyl ((2S,3R)-3-(tert-butoxy)-1-(((S)-1-hydrazineyl-4-methyl-1-oxopentan-2-yl)amino)-1-oxobutan-2-yl)carbamate (80b)**

To a solution of **80a** (4.36 g, 10 mmol) in methanol (50 mL) was dropwise added hydrazine hydrate (600 mg, 12 mmol). The resulting solution was stirred at room temperature overnight. Then the reaction mixture was evaporated to dryness *in vacuo* to afford **2** as white solid (3.86 g, 88%). $^{1}H$ NMR (400 MHz, Chloroform-*d*) δ 7.85 (s, 1H), 7.41 – 7.29 (m, 5H), 6.87 (d, *J* = 8.8 Hz, 1H), 5.88 (d, *J* = 5.2 Hz, 1H), 5.18 – 5.02 (m, 2H), 4.45 (td, *J* = 9.2, 5.0 Hz, 1H), 4.27 – 4.13 (m, 2H), 1.78 (ddd, *J* = 13.7, 8.7, 5.0 Hz, 1H), 1.70 – 1.58 (m, 1H), 1.51 (ddd, *J* = 13.7, 9.7, 5.2 Hz, 1H), 1.29 (s, 9H), 1.06 (d, *J* = 6.4 Hz, 3H), 0.93 (dd, *J* = 8.8, 6.5 Hz, 6H).

**Synthesis of benzyl ((2S,3R)-1-(((S)-1-(2-(3-amino-3-oxopropyl)hydrazineyl)-4-methyl-1-oxopentan-2-yl)amino)-3-(tert-butoxy)-1-oxobutan-2-yl)carbamate (80c)**

To a solution of **80b** (872 mg, 2 mmol) in DMF (2 mL) was added acrylamide (144 mg, 2 mmol). The reaction mixture was then heated to reflux for 36 h. The reaction mixture was then cooled and diluted with ethyl acetate (20 mL). The organic layer was washed with water and saturated brine, dried over anhydrous $Na_2SO_4$ and concentrated *in vacuo*. The residue was purified with flash chromatography (1-10% methanol in dichloromethane as the eluent) to yield **80c** as yellowish solid (185 mg, 18%). $^1$H NMR (400 MHz, Chloroform-*d*) δ 8.26 (s, 1H), 7.44 – 7.30 (m, 5H), 6.94 (d, *J* = 8.4 Hz, 2H), 5.88 (d, *J* = 5.8 Hz, 1H), 5.49 (s, 1H), 5.20 – 5.02 (m, 2H), 4.39 (td, *J* = 8.9, 5.3 Hz, 1H), 4.25 – 4.12 (m, 2H), 3.11 (t, *J* = 6.0 Hz, 2H), 2.35 (t, *J* = 6.1 Hz, 2H), 1.75 (ddd, *J* = 13.6, 8.4, 5.3 Hz, 1H), 1.64 (dp, *J* = 13.5, 6.4 Hz, 1H), 1.51 (ddd, *J* = 13.4, 9.4, 5.3 Hz, 1H), 1.26 (s, 9H), 1.08 (d, *J* = 6.3 Hz, 3H), 0.95 (d, *J* = 6.6 Hz, 3H), 0.92 (d, *J* = 6.4 Hz, 3H).

**Synthesis of benzyl ((2S,3R)-1-(((S)-1-(2-acryloyl-2-(3-amino-3-oxopropyl)hydrazineyl)-4-methyl-1-oxopentan-2-yl)amino)-3-(tert-butoxy)-1-oxobutan-2-yl)carbamate (MPI80)**

To a solution of **80c** (0.1 mmol, 50 mg) in anhydrous THF (5 mL) was added DIPEA (0.2 mmol, 26 mg). The solution was then cooled at -78 °C and acryloyl chloride (0.2 mmol, 23 mg) was added to the solution. The solution was stirred at -78 °C for 3 h for complete consumption of **3**. The reaction mixture was then allowed to warm up to room temperature, diluted with DCM (20 mL) and washed with 1 M HCl. The organic layers were combined, dried over anhydrous $Na_2SO_4$ and concentrated *in vacuo*. The residue was then purified with flash chromatography (1-10% methanol in dichloromethane as the eluent) to afford **MPI80** as yellowish solid (30 mg, 54 %). $^1$H NMR (400 MHz, Chloroform-*d*) δ 9.59 (s, 1H), 7.44 – 7.29 (m, 5H), 6.57 – 6.41 (m, 2H), 6.35 (d, *J* = 16.8 Hz, 1H), 5.96 (d, *J* = 5.4 Hz, 1H), 5.84 (s, 1H), 5.67 (d, *J* = 10.3 Hz, 1H), 5.11 (q, *J* = 12.2 Hz, 2H), 4.43 (s, 1H), 4.18 (q, *J* = 6.2, 5.0 Hz, 2H), 4.03 – 3.64 (m, 2H), 2.51 (s, 2H), 1.89 (s, 2H), 1.75 – 1.52 (m, 3H), 1.24 (s, 9H), 1.09 (d, *J* = 6.0 Hz, 3H), 0.97 (d, *J* = 5.7 Hz, 3H), 0.93 (d, *J* = 5.7 Hz, 3H). $^{13}$C NMR (101 MHz, $CDCl_3$) δ 174.1, 171.6, 170.5, 156.3, 136.0, 129.8, 128.6, 128.4, 128.2, 126.0, 77.2, 75.6, 67.2, 66.8, 59.1, 50.7, 40.3, 33.6, 28.2, 24.9, 22.9, 21.9, 17.6. HRMS (ESI+) m/z calculated for C28H43N5O7 $^+$ (M + H) $^+$ 562.3235, found 562.3221.

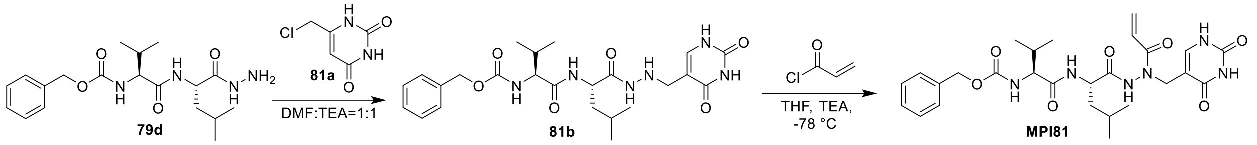


**Scheme 15**: The synthesis of **MPI81**

**Synthesis of benzyl ((S)-1-(((S)-1-(2-((2,4-dioxo-1,2,3,4-tetrahydropyrimidin-5-yl)methyl)hydrazineyl)-4-methyl-1-oxopentan-2-yl)amino)-3-methyl-1-oxobutan-2-yl)carbamate (81b).** To a solution of **79d** (100mg, 0.264mmol, 1.0equiv) in 0.5 ml anhydrous DMF and 0.5 ml triethylamine, was added compound **81a** (42mg, 0.264mmol,1.0equiv), and the mixture was refluxed to 50°C overnight. The solvent was removed under reduced pressure and the residue was purified by column chromatography (MeOH: ethyl acetate = 1:10 v/v) to afford the pure product **81b** as a white solid (50 mg, 38%). $^{1}$H NMR (400 MHz, Methanol-$d_4$) δ 7.33 – 7.11 (m, 5H), 5.54 (d, *J* = 7.9 Hz, 1H), 4.99 (d, *J* = 4.2 Hz, 2H), 4.55 – 4.12 (m, 2H), 3.82 (q, *J* = 6.7, 5.5 Hz, 1H), 3.61 (s, 1H), 3.50 (s, 1H), 2.05 – 1.78 (m, 3H), 1.67 – 1.34 (m, 4H), 0.96 (t, *J* = 7.1 Hz, 1H), 0.88 – 0.71 (m, 13H).

**Synthesis of benzyl ((S)-1-(((S)-1-(2-acryloyl-2-((2,4-dioxo-1,2,3,4-tetrahydropyrimidin-5-yl)methyl)hydrazineyl)-4-methyl-1-oxopentan-2-yl)amino)-3-methyl-1-oxobutan-2-yl)carbamate (MPI81).** To a stirred solution of **81b** (44 mg,0.088 mmol, 1.0 equiv) in THF (5 mL) at −78 ◦C was triethylamine (27 µl, 0.176mmol, 2.0equiv). After 15 min, acryloyl chloride (10 µL, 0.105 mmol, 1.2 equiv.) was added, and the mixture was stirred for further 3h. The reaction was quenched with water (5 mL), and the mixture was concentrated in vacuum. The residue was partitioned between EtOAc (20 mL) and $H_2O$ (10 mL). The aqueous layer was extracted with EtOAc (2 × 20 mL). The combined organic layer was washed with brine, dried over $MgSO_4$ and concentrated in vacuum. The residue was purified by flash column chromatography (MeOH: $CH_2Cl_2$ 0.5:9.5) to give the title compound as a white solid (20 mg, 42%). $^{1}$H NMR (400 MHz, Chloroform-*d*) δ 7.47 – 7.14 (m, 4H), 6.66 (d, *J* = 12.8 Hz, 1H), 6.36 (d, *J* = 17.0 Hz, 1H), 5.96 – 5.74 (m, 1H), 5.60 (s, 1H), 5.07 (s, 2H), 4.63 – 4.23 (m, 2H), 3.98 – 3.83 (m, 1H), 2.20 – 1.88 (m, 2H), 1.76 – 1.46 (m, 3H), 1.43 – 1.04 (m, 5H), 1.01 – 0.78 (m, 12H). HRMS (ESI+) m/z calculated for C27H36N6O7 $^{+}$ (M + H)$^{+}$ 557.2718, found 557.2709.

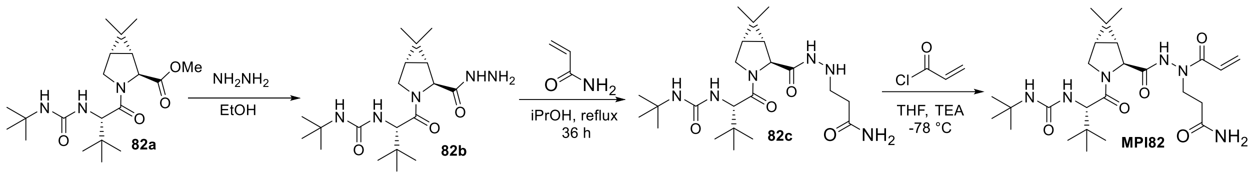

**Scheme 16**: The synthesis of **MPI82**

**Synthesis of 1-(tert-butyl)-3-((S)-1-((1R,2S,5S)-2-(hydrazinecarbonyl)-6,6-dimethyl-3-azabicyclo[3.1.0]hexan-3-yl)-3,3-dimethyl-1-oxobutan-2-yl)urea (82b).** To a solution of compound **82a** (500 mg, 1.0 mmol, 1.0 equiv) in ethanol, was added hydrazine (352 mg, 11.0 mmol, 11.0 equiv). The reaction mixture was stirred at RT overnight. After the reaction was completed, the solvent was removed on *vacuo*. The residue was used in the next step without further purification.

**Synthesis of 3-(2-((1R,2S,5S)-3-((S)-2-(3-(tert-butyl)ureido)-3,3-dimethylbutanoyl)-6,6-dimethyl-3-azabicyclo[3.1.0]hexane-2-carbonyl)hydrazinyl)propanamide (82c).** To a solution of compound **82b** (1.0 g, 2.62 mmol) in PrOH (1.5 mL), was added acrylamide (0.186 g, 2.62 mmol) and the solution refluxed for 36 h. The solution is cooled, filtered and the solvent removed in vacuum to give viscous oil. The oily compound was purified by chromatography column (Ethyl acetate ⁄ methanol 9:1). (Yield 90 mg, 8%) and 500 mg of the starting material was recovered .$^{1}$H NMR (400 MHz, Methanol-$d_4$) δ 4.16 (d, $J$ = 2.6 Hz, 1H), 4.14 (s, 1H), 3.95 (d, $J$ = 10.4 Hz, 1H), 3.85 (dd, $J$ = 10.3, 5.4 Hz, 1H), 3.29 (t, $J$ = 7.8 Hz, 1H), 3.25 (s, 3H), 2.95 (td, $J$ = 6.7, 3.3 Hz, 2H), 2.29 (td, $J$ = 6.7, 2.0 Hz, 2H), 1.48 (dd, $J$ = 7.6, 5.2 Hz, 1H), 1.27 (d, $J$ = 7.6 Hz, 1H), 1.16 (s, 9H), 0.96 (s, 3H), 0.89 (s, 9H), 0.83 (s, 3H). $^{13}$C NMR (101 MHz, MeOD): δ 175.95, 172.17, 171.45, 158.31, 59.24, 57.61, 49.32, 48.46, 45.66, 34.35, 33.33, 31.79, 30.84, 28.26, 27.80, 25.61, 25.11, 19.04, 18.97, 11.74.

**Synthesis of 3-(1-acryloyl-2-((1R,2S,5S)-3-((S)-2-(3-(tert-butyl)ureido)-3,3-dimethylbutanoyl)-6,6-dimethyl-3-azabicyclo[3.1.0]hexane-2-carbonyl)hydrazinyl)propanamide (MPI82).** To a stirred solution of **82c** (50 mg, 0.11 mmol) in THF (5 mL) at −78 ◦C was added $Et_3N$ (31 µL, 0.221 mmol). After 15 min, acryloyl chloride (11 µL, 0.132 mmol) was added, and the mixture was stirred for further 30 min. The reaction was quenched with water (5 mL), and the mixture was concentrated in vacuum. The residue was partitioned between EtOAc (20 mL) and $H_2O$ (10 mL). The aqueous layer was extracted with EtOAc (2 × 20 mL). The combined organic layer was washed with brine, dried over $MgSO_4$ and concentrated in vacuum. The residue was purified by flash column chromatography (MeOH: $CH_2Cl_2$ 0.5:9.5) to give the title compound as a white solid (25 mg, 49%).$^1$H NMR (400 MHz, Chloroform-*d*) δ 10.07 (s, 1H), 6.92 (s, 1H), 6.54 (dd, *J* = 16.8, 10.4 Hz, 1H), 6.37 – 6.22 (m, 1H), 6.18 – 6.05 (m, 1H), 5.54 (s, 2H), 4.31 (d, *J* = 9.8 Hz, 2H), 4.09 (d, *J* = 10.4 Hz, 1H), 3.93 – 3.59 (m, 3H), 2.50 (s, 2H), 1.53 (dd, *J* = 7.6, 5.3 Hz, 1H), 1.34 (d, *J* = 6.3 Hz, 1H), 1.19 (s, 9H), 0.98 (s, 5H), 0.86 (d, *J* = 2.3 Hz, 12H).

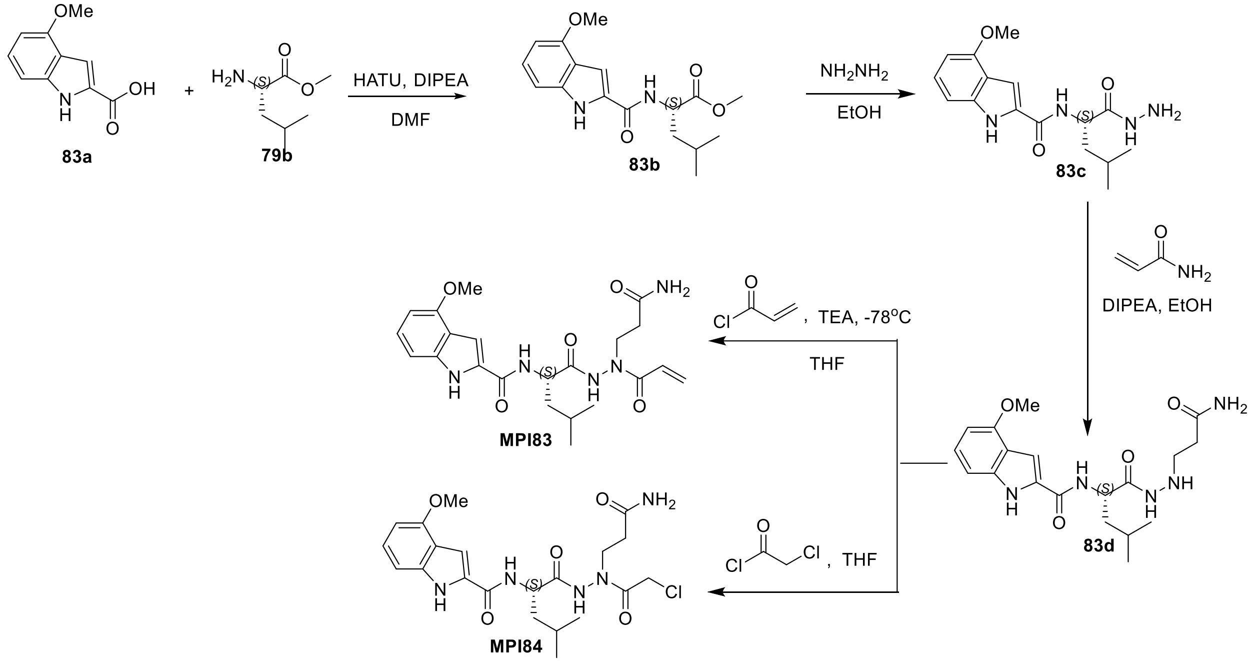


**Scheme 17**: The synthesis of **MPI83** and **MPI84**

**Synthesis of methyl (4-methoxy-1H-indole-2-carbonyl)-L-leucinate (83b).** To a solution of **83a** (800 mg, 4.18 mmol, 1.0 equiv) in anhydrous DMF (10 mL) at 0 °C, and then **79b** (838 mg, 4.6 mmol, 1.1 equiv), HATU (2 g, 5.43 mmol, 1.3 equiv), DIPEA (3 mL, 16.72 mmol, 4.0 equiv) was added sequentially. The mixture was stirred at RT overnight. The mixture was diluted with EtOAc and washed with water, 1M HCl, sat. NaCl, dried over $Na_2SO_4$, and concentrated. The residue was purified by column chromatography (Hexane: EA = 3:1 v/v) to afford the pure product **83b** as a colorless oil. $^{1}$H NMR (400 MHz, Chloroform-*d*) δ 9.68 (s, 1H), 7.22 (t, *J* = 8.0 Hz, 1H), 7.17 – 7.00 (m, 2H), 6.70 (d, *J* = 8.4 Hz, 1H), 6.53 (d, *J* = 7.7 Hz, 1H), 5.03 – 4.75 (m, 1H), 3.97 (s, 3H), 3.80 (s, 3H), 1.87 – 1.65 (m, 3H), 1.01 (t, *J* = 5.6 Hz, 6H). $^{13}$C NMR (100 MHz, Chloroform-*d*) δ 173.46, 161.45, 154.18, 137.93, 128.73, 125.62, 118.87, 105.17, 100.55, 99.65, 55.31, 52.47, 50.91, 41.80, 24.99, 22.86, 22.02.

**Synthesis of (S)-N-(1-hydrazineyl-4-methyl-1-oxopentan-2-yl)-4-methoxy-1H-indole-2-carboxamide (83c).** Compound **83b** (1.26 g, 3.96 mmol, 1.0 equiv) and hydrazine (1.27 g, 39.6 mmol, 10.0 equiv) were dissolved in ethanol (25 mL). The reaction mixture was stirred at RT overnight. After the reaction was completed, the solvent was removed on *vacuo*. The residue was used in the next step without further purification. $^{1}$H NMR (400 MHz, Chloroform-*d*) δ 10.80 (s, 1H), 9.78 (s, 1H), 8.08 (d, *J* = 9.1 Hz, 1H), 7.41 (s, 1H), 6.92 (t, *J* = 8.0 Hz, 1H), 6.66 (d, *J* = 8.3 Hz, 1H), 6.23 (d, *J* = 7.8 Hz, 1H), 4.98 – 4.69 (m, 1H), 3.63 (s, 3H), 1.86 – 1.40 (m, 3H), 0.84 (dd, *J* = 11.3, 6.4 Hz, 6H). $^{13}$C NMR (100 MHz, Chloroform-*d*) δ 173.09, 161.90, 154.05, 138.22, 129.20, 125.14, 118.64, 105.20, 101.75, 99.23, 55.20, 50.08, 40.86, 24.90, 22.80, 22.43.

**Synthesis of (S)-N-(1-(2-(3-amino-3-oxopropyl)hydrazineyl)-4-methyl-1-oxopentan-2-yl)-4-methoxy-1H-indole-2-carboxamide (83d).** To a solution of **83c** (500 mg, 1.57 mmol,1.0 equiv), acrylamide (1.34 mg, 1.89 mmol, 1.2 equiv) and DIPEA (1.4 mL, 7.85 mmol, 5.0 equiv) in ethanol. The reaction mixture was refluxed for 48 h. After the reaction was completed, the solution was removed on *vacuo*. The resulted residue was purified by column chromatography (DCM: MeOH = 10:1 v/v) to afford the pure product **83d** as a white solid (312 mg, yield 51%). $^{1}$H NMR (400 MHz, Chloroform-*d*) δ 10.58 (s, 1H), 9.44 (s, 1H), 7.68 (s, 1H), 7.17 (s, 1H), 7.08 – 6.86 (m, 3H), 6.54 (s, 1H), 6.46 – 6.16 (m, 1H), 4.60 (q, *J* = 8.3, 7.1 Hz, 1H), 3.76 (s, 3H), 3.33 (t, *J* = 5.9 Hz, 1H), 2.90 (q, *J* = 9.7, 6.4 Hz, 2H), 2.45 (t, *J* = 5.9 Hz, 1H), 2.24 – 2.05 (m, 2H), 2.01 – 1.91 (m, 1H), 1.85 (d, *J* = 6.3 Hz, 1H), 1.78 (s, 1H), 1.69 – 1.50 (m, 4H), 1.41 – 1.24 (m, 1H), 0.98 – 0.63 (m, 6H).

**Synthesis of (S)-N-(1-(2-acryloyl-2-(3-amino-3-oxopropyl)hydrazineyl)-4-methyl-1-oxopentan-2-yl)-4-methoxy-1H-indole-2-carboxamide (MPI83).** To a solution of **83d** (50 mg, 0.13 mmol, 1.0 equiv) in anhydrous THF at -78 $^{o}$C was added TEA (36 μL, 0.26 mmol, 2.0 equiv) and acryloyl chloride (14 mg, 0.15 mmol, 1.2 equiv) at this temperature. The reaction was quenched by slow addition of $H_2O$ and removed THF on *vacuo*. The mixture was diluted with $H_2O$ and extracted with EtOAc, washed with sat. NaCl, dried over $Na_2SO_4$ and concentrated. The residue was purified by column chromatography (DCM: MeOH = 10:1 v/v) to afford the pure product **MPI83** (36 mg, yield 63%) as a white solid. $^{1}$H NMR (400 MHz, Chloroform-*d*) δ 9.42 (d, *J* = 35.2 Hz, 2H), 7.13 (d, *J* = 8.0 Hz, 1H), 7.08 – 6.98 (m, 1H), 6.95 (d, *J* = 8.3 Hz, 1H), 6.50 – 6.19 (m, 3H), 6.05 (s, 2H), 5.52 (d, *J* = 37.4 Hz, 2H), 4.66 (s, 1H), 3.86 (s, 3H), 2.44 (s, 2H), 1.87 – 1.60 (m, 3H), 1.17 (d, *J* = 1.8 Hz, 2H), 0.91 (dd, *J* = 11.8, 5.9 Hz, 6H). HRMS (APCI+) m/z calculated for $C22H29N5O5^{+}$ $(M + H)^{+}$ 444.2241, found 444.2239.

**Synthesis of (S)-N-(1-(2-(3-amino-3-oxopropyl)-2-(2-chloroacetyl)hydrazineyl)-4-methyl-1-oxopentan-2-yl)-4-methoxy-1H-indole-2-carboxamide (MPI84).** To a solution of **83d** (50 mg, 0.13 mmol, 1.0 equiv) in anhydrous THF at 0 $^{o}$C and then added chloroacetyl chloride (12.5 mg, 0.15 mmol,1.2 equiv). The resulted mixture was stirred at RT for 2 h. After the reaction was completed, removed the solvent on vacuo and purified by column chromatography (DCM: MeOH = 10:1 v/v) to afford the pure product **MPI84** (40 mg, yield 67%) as a white solid. $^{1}$H NMR (400 MHz, Chloroform-*d*) δ 10.28 (s, 1H), 9.85 (d, *J* = 51.6 Hz, 1H), 7.17 – 7.01 (m, 3H), 6.94 (d, *J* = 8.3 Hz, 1H), 6.51 (s, 1H), 6.38 (d, *J* = 7.8 Hz, 1H), 6.04 (s, 1H), 4.63 (s, 1H), 4.19 – 3.89 (m, 2H), 3.82 (s, 3H), 3.74 – 3.61 (m, 1H), 2.30 (s, 2H), 2.04 – 1.86 (m, 1H), 1.85 – 1.74 (m, 1H), 1.69 (d, *J* = 6.0 Hz, 3H), 0.97 – 0.75 (m, 6H). HRMS (APCI+) m/z calculated for $C21H28ClN5O5^{+}$ $(M + H)^{+}$ 466.1852, found 466.1849.

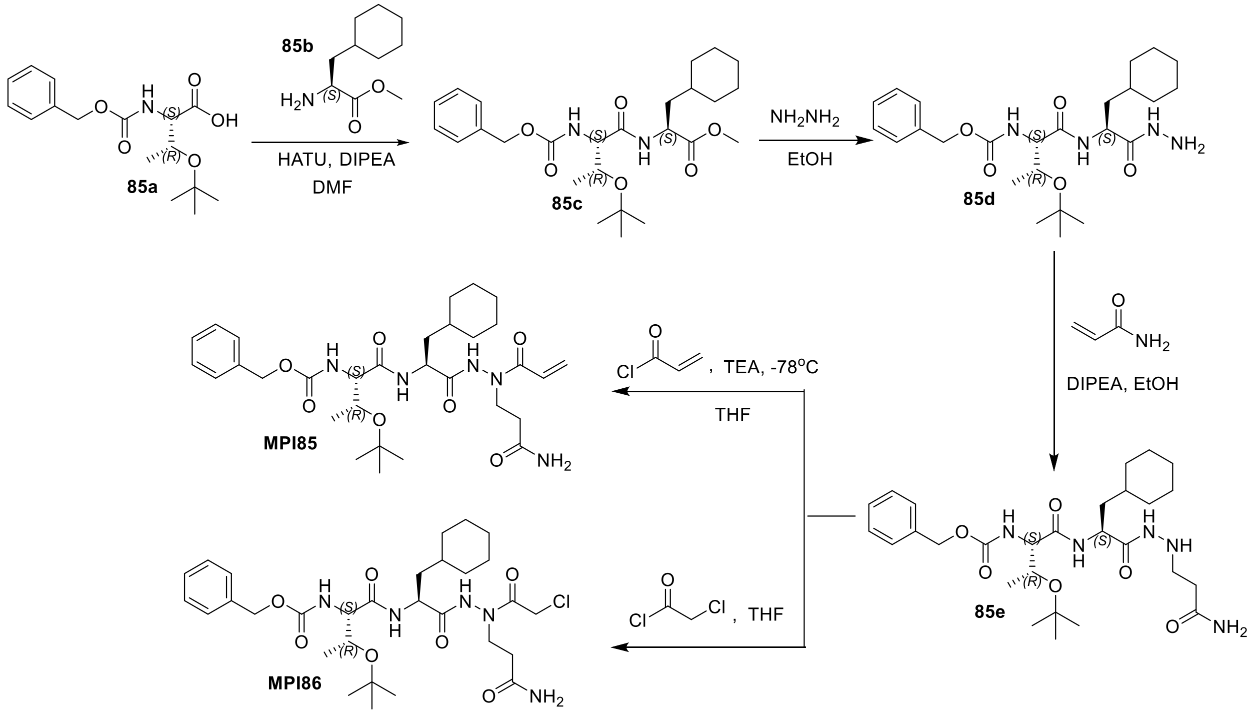


**Scheme 18.** The synthesis of **MPI85** and **MPI86**.

**Synthesis of Methyl (S)-2-((2S,3R)-2-(((benzyloxy)carbonyl)amino)-3-(tert-butoxy)butanamido)-3-cyclohexylpropanoate (85c).** To a solution of compound **85a** (500 mg, 1.6 mmol, 1.0 equiv) in anhydrous DMF (5 mL) at 0 °C, and then compound **85b** (360 mg, 1.6 mmol, 1.0 equiv), HATU (912 mg, 2.4 mmol, 1.5 equiv), DIPEA (1.1 mL, 6.4 mmol, 4.0 equiv) was added sequentially. The mixture was stirred at RT overnight. The mixture was diluted with EtOAc and washed with water, 1M HCl, sat. NaCl, dried over $Na_2SO_4$, and concentrated. The residue was purified by column chromatography (Hexane: EA = 3:1 v/v) to afford the pure product **85c** as a colorless oil (570 mg, yield 75%). $^1$H NMR (400 MHz, Chloroform-*d*) δ 7.7 (d, *J* = 7.8 Hz, 1H), 7.4 – 7.3 (m, 5H), 6.1 (d, *J* = 5.3 Hz, 1H), 5.3 – 5.1 (m, 2H), 4.6 (td, *J* = 8.4, 5.1 Hz, 1H), 4.3 – 4.2 (m, 2H), 3.7 (s, 3H), 1.8 – 1.7 (m, 6H), 1.6 (ddd, *J* = 14.2, 9.0, 5.7 Hz, 1H), 1.3 (s, 9H), 1.2 (dd, *J* = 24.2, 7.4 Hz, 7H), 1.0 – 0.9 (m, 2H). $^{13}$C NMR (100 MHz, Chloroform-*d*) δ 172.7, 169.2, 155.9, 136.2, 128.3, 128.3, 127.8, 127.7, 127.7, 75.2, 66.7, 66.5, 60.0, 51.8, 50.2, 39.5, 34.0, 33.3, 32.3, 28.0, 28.0, 28.0, 26.1, 26.0, 25.8, 20.7.

**Synthesis of Benzyl ((2S,3R)-3-(tert-butoxy)-1-(((S)-3-cyclohexyl-1-hydrazineyl-1-oxopropan-2-yl)amino)-1-oxobutan-2-yl)carbamate (85d).** To a solution of compound **85c** (500 mg, 1.0 mmol, 1.0 equiv) in ethanol, was added hydrazine (350 mg, 11.0 mmol, 11.0 equiv). The reaction mixture was stirred at RT overnight. After the reaction was completed, the solvent was removed on *vacuo*. The residue was used in the next step without further purification. $^{1}$H NMR (400 MHz, Methanol-$d_4$) δ 7.4 (ddt, *J* = 19.6, 13.8, 6.8 Hz, 5H), 5.1 (d, *J* = 3.4 Hz, 2H), 4.5 (dd, *J* = 9.4, 5.5 Hz, 1H), 4.2 – 4.1 (m, 2H), 1.9 – 1.6 (m, 7H), 1.4 (tq, *J* = 5.1, 2.8 Hz, 1H), 1.2 (d, *J* = 36.1 Hz, 15H), 1.0 – 0.9 (m, 2H). $^{13}$C NMR (100 MHz, Methanol-$d_4$) δ 173.6, 172.2, 158.5, 138.0, 129.5, 129.1, 129.0, 76.0, 68.8, 67.9, 61.3, 51.2, 40.7, 35.2, 34.8, 33.5, 28.6, 27.5, 27.3, 27.1, 19.6.

**Synthesis of Benzyl ((2S,3R)-1-(((S)-1-(2-(3-amino-3-oxopropyl)hydrazineyl)-3-cyclohexyl-1-oxopropan-2-yl)amino)-3-(tert-butoxy)-1-oxobutan-2-yl)carbamate (85e).** To a solution of **85d** (300 mg, 0.63 mmol,1.0 equiv), acrylamide (54 mg, 0.76 mmol, 1.2 equiv) and DIPEA (550 μL, 3.15 mmol, 5.0 equiv) in ethanol. The reaction mixture was refluxed for 48 h. After the reaction was completed, the solution was removed on *vacuo*. The resulted residue was purified by column chromatography (EA: MeOH = 8:1 v/v) to afford the pure product **85e** as a white solid (150 mg, yield 44%).$^{1}$H NMR (400 MHz, Methanol-$d_4$) δ 7.4 – 7.3 (m, 5H), 5.2 – 5.1 (m, 2H), 4.4 (dd, *J* = 9.2, 5.8 Hz, 1H), 4.2 – 4.1 (m, 2H), 3.0 (t, *J* = 6.8 Hz, 2H), 2.4 (t, *J* = 6.8 Hz, 2H), 1.8 – 1.6 (m, 7H), 1.4 – 1.4 (m, 1H), 1.2 (d, *J* = 25.5 Hz, 15H), 1.0 – 0.9 (m, 2H).$^{13}$C NMR (100 MHz, Methanol-$d_4$) δ 175.8, 171.8, 171.2, 157.2, 136.7, 128.1, 127.7, 127.6, 74.5, 67.4, 66.6, 60.2, 50.0, 47.2, 39.2, 33.9, 33.4, 33.2, 32.2, 27.3, 26.2, 26.0, 25.7, 18.4.

**Benzyl ((2S,3R)-1-(((S)-1-(2-acryloyl-2-(3-amino-3-oxopropyl)hydrazineyl)-3-cyclohexyl-1-oxopropan-2-yl)amino)-3-(tert-butoxy)-1-oxobutan-2-yl)carbamate (MPI85).** To a solution of **85e** (100 mg, 0.18 mmol, 1.0 equiv) in anhydrous THF at -78 $^{o}$C. Added TEA (50 μL, 0.36 mmol, 2.0 equiv) and acryloyl chloride (20 mg, 0.22 mmol, 1.2 equiv) at this temperature. The reaction was quenched by slow addition of $H_2O$ and removed THF on *vacuo*. The mixture was diluted with $H_2O$ and extracted with EtOAc, washed with sat. NaCl, dried over $Na_2SO_4$ and concentrated. The residue was purified by column chromatography (EA: MeOH = 8:1 v/v) to afford the pure product **MPI85** (40 mg, yield 37%) as a white solid. $^{1}$H NMR (400 MHz, Methanol-$d_4$) δ 7.4 – 7.3 (m, 5H), 6.3 (dd, *J* = 16.9, 2.0 Hz, 1H), 5.8 (d, *J* = 10.6 Hz, 1H), 5.2 – 5.1 (m, 2H), 4.4 (t, *J* = 7.5 Hz, 1H), 4.2 – 4.1 (m, 2H), 3.7 (dq, *J* = 13.2, 6.9 Hz, 1H), 2.5 (s, 2H), 1.9 – 1.6 (m, 7H), 1.5 (s, 1H), 1.2 (d, *J* = 21.1 Hz, 15H), 1.1 – 0.9 (m, 4H). $^{13}$C NMR (100 MHz, Methanol-$d_4$) δ 174.6, 172.5, 171.7, 168.1, 157.3, 136.6, 128.8, 128.1, 127.7, 127.6, 126.3, 74.5, 67.4, 66.6, 60.2, 50.0, 44.8, 38.6, 33.9, 33.3, 32.7, 32.1, 27.4, 26.1, 26.0, 25.8, 18.4. HRMS (APCI+) m/z calculated for C31H47N5O7 $^{+}$ (M + H) $^{+}$ 602.3548, found 602.3541.

**Synthesis of benzyl (2S,3R)-1-(((S)-1-(2-(3-amino-3-oxopropyl)-2-(2-chloroacetyl)hydrazineyl)-3-cyclohexyl-1-oxopropan-2-yl)amino)-3-(tert-butoxy)-1-oxobutan-2-yl)carbamate (MPI86).** To a solution of **85e** (560 mg, 0.9 mmol, 1.0 equiv) in anhydrous THF at 0 $^{o}$C and then added chloroacetyl chloride (100 mL, 1.08 mmol,1.2 equiv). The resulting mixture was stirred at RT for 2 h. After the reaction was completed, the solvent was removed on vacuo and purified by column chromatography (DCM: MeOH = 10:1 v/v) to afford the pure product **MPI86** (420 mg, yield 75%) as a white solid. $^{1}$H NMR (400 MHz, Chloroform-*d*) δ 9.21 (s, 1H), 7.34 – 7.23 (m, 5H), 6.23 (dd, *J* = 17.1, 1.4 Hz, 1H), 6.08 (dd, *J* = 17.1, 10.2 Hz, 1H), 6.02 – 5.78 (m, 2H), 5.66 (dd, *J* = 10.3, 1.4 Hz, 1H), 5.15 – 4.93 (m, 2H), 4.65 – 4.49 (m, 1H), 4.28 – 3.93 (m, 5H), 3.93 – 3.80 (m, 1H), 3.73 (t, *J* = 6.4 Hz, 1H), 2.62 (t, *J* = 6.5 Hz, 1H), 2.55 – 2.36 (m, 1H), 2.27 – 1.89 (m, 3H), 1.74 – 1.47 (m, 8H), 1.24 – 1.13 (m, 5H), 1.13 – 0.99 (m, 6H), 0.96 – 0.69 (m, 2H). HRMS (ESI+) m/z calculated for C30H46ClN5O7 $^{+}$ (M + H) $^{+}$ 624.3159, found 624.3161.

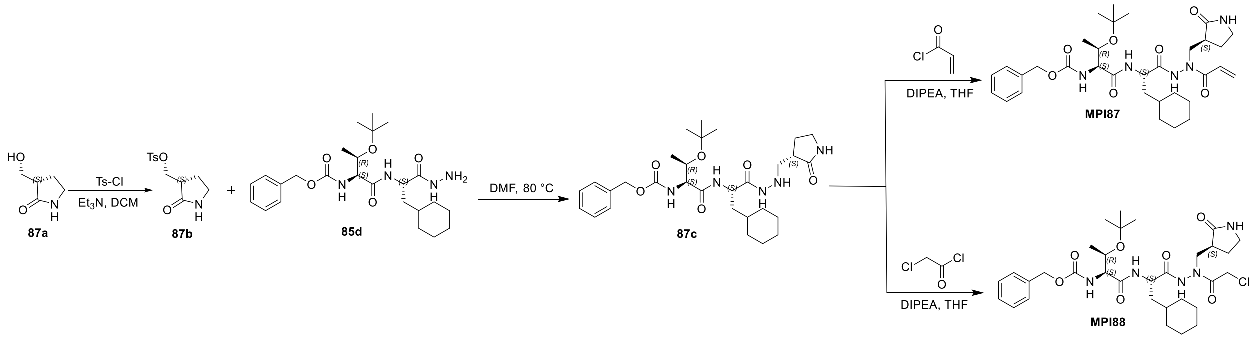


+

**Scheme 19.** The synthesis of **MPI87** and **MPI88**.

**Synthesis of (2-oxopyrrolidin-3-yl)methyl 4-methylbenzenesulfonate (87b).** To a solution of compound **87a** (0.3 g, 2.60 mmol) in dry $CH_2Cl_2$ (10 mL), was added TsCl (0.74 g, 3.9 mmol), $Et_3N$ (0.54 mL, 12.9 mmol), and a catalytic amount of DMAP, under nitrogen. The reaction mixture was stirred at room temperature for 2 h. Quench the reaction mixture with water. Extract the aqueous layer with $CH_2Cl_2$ (3 x 30 mL). Wash the combined organic layers with brine. Concentrate the combined organic layers and evaporated under reduced pressure which was carried forward to the next step without further purification.

**Synthesis of benzyl ((2*S*,3*R*)-3-(*tert*-butoxy)-1-(((2*S*)-3-cyclohexyl-1-oxo-1-(2-((2-oxopyrrolidin-3-yl)methyl)hydrazineyl)propan-2-yl)amino)-1-oxobutan-2-yl)carbamate (87c).** To a stirred solution of **87b** (0.2 g, 0.74 mmol) in 6.0 mL of DMF, was added compound **85d** (0.48 g, 1.48 mmol), and heated to 90 ° C for 6 hours. The mixture was diluted with EtOAc and washed with water, 1M HCl, sat. NaCl, dried over $Na_2SO_4$, and concentrated. The residue was purified by column chromatography (methanol: dichloromethane = 1:10 v/v) to afford the pure product **87c** as a colorless solid. $^1$H NMR (400 MHz, Chloroform-*d*) δ 8.54 (d, *J* = 2.4 Hz, 1H), 7.31 – 7.20 (m, 4H), 7.23 – 7.12 (m, 1H), 6.50 (d, *J* = 11.4 Hz, 1H), 5.91 (q, *J* = 5.7, 4.7 Hz, 1H), 5.05 (d, *J* = 12.3 Hz, 2H), 4.34 (td, *J* = 8.8, 5.5 Hz, 1H), 4.10 (m, 2H), 3.24 (q, *J* = 8.1, 5.9 Hz, 2H), 3.05 (m, 1H), 2.84 (m, 1H), 2.49 (m, 1H), 2.22 (m, 1H), 1.87 (m, 1H), 1.70 – 1.52 (m, 5H), 1.48-1.39 (m, 1H), 1.28-1.05 (m, 11H), 1.00 (d, *J* = 6.0 Hz, 3H), 0.95 – 0.74 (m, 2H). $^{13}$C NMR (101 MHz, $CDCl_3$) δ 179.13, 179.12, 170.75, 170.71, 169.64, 169.61, 156.23, 145.90, 136.15, 128.62, 128.57, 128.52, 128.24, 128.09, 75.61, 67.02, 66.90, 58.96, 52.73, 51.87, 50.07, 40.53, 40.13, 34.10, 34.07, 33.59, 32.55, 28.23, 26.34, 26.18, 25.97, 25.83, 17.43.

**Synthesis of benzyl ((2*S*,3*R*)-3-(*tert*-butoxy)-1-(((2*S*)-1-(2-(2-chloroacetyl)-2-((2-oxopyrrolidin-3-yl)methyl)hydrazineyl)-3-cyclohexyl-1-oxopropan-2-yl)amino)-1-oxobutan-2-yl)carbamate (MPI87).** To a stirred solution of **87c** (50 mg, 0.085 mmol) in THF (5 mL) at −78 ◦C was added $Et_3N$ (25 µL, 0.12 mmol). After 15 min, acryloyl chloride (9 µL, 0.10 mmol) was added, and the mixture was stirred for a further 30 min. The reaction was quenched with water (5 mL), and the mixture was concentrated in a vacuum. The residue was partitioned between EtOAc (20 mL) and $H_2O$ (10 mL). The aqueous layer was extracted with EtOAc (2 × 20 mL). The combined organic layer was washed with brine, dried over $MgSO_4$, and concentrated in a vacuum. The residue was purified by flash column chromatography (MeOH: $CH_2Cl_2$ 0.5:9.5) to give the title compound a white solid (25 mg). $^1$H NMR (400 MHz, Chloroform-*d*) δ 7.44 (m, *J* = 19.7, 7.6 Hz, 2H), 7.28 (m, *J* = 10.7 Hz, 2H), 7.28 – 7.18 (m, 2H), 6.46 (m, *J* = 16.8, 10.1 Hz, 3H), 6.33 (m, *J* = 16.5 Hz, 1H), 6.27 (m, *J* = 5.5 Hz, 1H), 6.28 – 6.18 (m, 1H), 5.91 (s, 1H), 5.63 – 5.54 (m, 2H), 5.22 (s, 1H), 5.04 (qd, *J* = 12.2, 3.9 Hz, 4H), 4.41 (t, *J* = 7.5 Hz, 1H), 4.30 (s, 2H), 4.22 – 4.00 (m, 4H), 3.67 (h, *J* = 4.3 Hz, 1H), 3.22 (dt, *J* = 13.3, 8.6 Hz, 3H), 2.70 (s, 3H), 2.28 – 2.15 (m, 2H),

1.97 (s, 0H), 1.83 – 1.72 (m, 1H), 1.73 – 1.60 (m, 7H), 1.60 (s, 7H), 1.59 – 1.47 (m, 1H), 1.32 – 1.22 (m, 2H), 1.19 (s, 15H), 1.19 – 1.05 (m, 4H), 1.01 (d, $J$ = 6.4 Hz, 7H), 0.95 – 0.76 (m, 2H). $^{13}$C NMR (101 MHz, $CDCl_3$) δ 14.20, 21.05, 25.61, 26.01, 26.23, 26.30, 28.24, 32.48, 32.54, 33.63, 34.15, 34.19, 38.92, 39.07, 40.54, 50.04, 53.46, 58.97, 60.40, 66.77, 66.95, 67.07, 67.12, 67.96, 75.49, 76.77, 77.09, 77.40, 126.27, 126.32, 128.18, 128.24, 128.32, 128.35, 128.62, 128.82, 129.33, 136.06, 136.11, 156.30, 169.08, 170.10, 170.24, 171.18. HRMS (ESI+) m/z calculated for C33H49N5O7 $^+$ (M + H) $^+$ 628.3705, found 628.3687.

**Synthesis of benzyl ((2*S*,3*R*)-1-(((2*S*)-1-(2-acryloyl-2-((2-oxopyrrolidin-3-yl)methyl)hydrazineyl)-3-cyclohexyl-1-oxopropan-2-yl)amino)-3-(*tert*-butoxy)-1-oxobutan-2-yl)carbamate (MPI88).** To a stirred solution of **87c** (50 mg, 0.085 mmol) in THF (5 mL) at 0 °C, chloro acetylchloride (11 µL, 0.10) was added dropwise, and the mixture was stirred for further 30 min at the same temperature. The reaction was quenched with water (5 mL), and the mixture was concentrated in a vacuum. The residue was partitioned between EtOAc (30 mL) and $H_2O$ (20 mL). The aqueous layer was extracted with EtOAc (2 × 20 mL). The combined organic layer was washed with brine, dried over MgSO4, and concentrated in a vacuum. The residue was purified by flash column chromatography (0 to 6% MeOH in $CH_2Cl_2$ as the eluent)) to give the title compound **6** as white solid (30 mg). $^1$H NMR (400 MHz, Chloroform-*d*) δ 7.53 (t, $J$ = 7.6 Hz, 2H), 7.37 (q, $J$ = 4.1, 3.6 Hz, 11H), 6.87 (d, $J$ = 15.6 Hz, 1H), 6.76 (s, 1H), 6.05 (s, 1H), 5.96 (d, $J$ = 5.4 Hz, 1H), 5.76 (s, 6H), 5.19 – 5.05 (m, 4H), 4.37 (d, $J$ = 24.2 Hz, 2H), 4.28 – 4.18 (m, 7H), 4.10 (d, $J$ = 12.3 Hz, 1H), 4.07 (s, 3H), 4.04 (s, 1H), 4.01 (s, 1H), 3.80 (s, 1H), 3.34 (dt, $J$ = 16.9, 7.4 Hz, 4H), 3.11 (s, 1H), 2.83 (s, 3H), 2.32 (dq, $J$ = 19.9, 8.2, 7.2 Hz, 2H), 1.88 (s, 2H), 1.80 – 1.59 (m, 16H), 1.60 (d, $J$ = 5.7 Hz, 1H), 1.35 (d, $J$ = 14.8 Hz, 3H), 1.27 (d, $J$ = 4.3 Hz, 18H), 1.15 (dt, $J$ = 23.9, 5.2 Hz, 11H), 1.03 – 0.85 (m, 3H).$^{13}$C NMR (101 MHz, $CDCl_3$) δ 14.12, 25.39, 25.96, 26.22, 26.27, 28.24, 32.34, 33.67, 34.14, 34.18, 38.60, 40.90, 40.93, 41.79, 50.37, 53.46, 59.14, 66.72, 67.20, 75.55, 77.28, 128.20, 128.39, 128.64, 136.00, 156.39, 170.53. HRMS (ESI+) m/z calculated for C32H48ClN5O7 $^+$ (M + H) $^+$ 650.3315, found 650.3301.

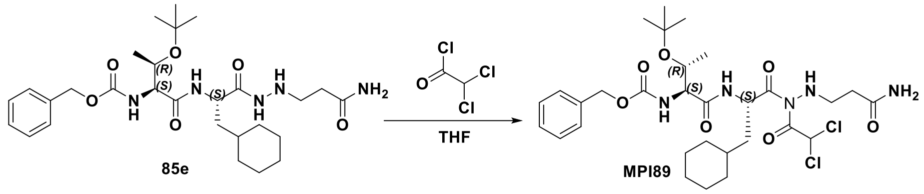


**Scheme 20.** The synthesis of **MPI89**.

**Synthesis of Benzyl ((2S,3R)-1-(((S)-1-(2-(3-amino-3-oxopropyl)-1-(2,2-dichloroacetyl)hydrazineyl)-3-cyclohexyl-1-oxopropan-2-yl)amino)-3-(tert-butoxy)-1-oxobutan-2-yl)carbamate (MPI89).** To a stirred solution of **85e** (0.05 g, 0.0914 mmol) in THF (5 mL) at 0 ◦C, dichloroacetylchloride (10.51 µL, 0.109 mmol) was added drop wise, and the mixture was stirred for further 30 min at same temperature. The reaction was quenched with water (5 mL), and the mixture was concentrated in vacuum. The residue was partitioned between EtOAc (10 mL) and $H_2O$ (10 mL). The aqueous layer was extracted with EtOAc (2 × 10 mL). The combined organic layer was washed with brine, dried over $MgSO_4$ and concentrated in vacuum. The residue was purified by flash column chromatography (0 to 6% MeOH in $CH_2Cl_2$ as the eluent)) to give the title compound **MPI89** as white solid (15 mg, 25%). $^1$H NMR (400 MHz, DMSO-$d_6$) δ 10.97 (s, 1H), 8.11 (s, 1H), 7.43 – 7.25 (m, 5H), 6.98 (d, *J* = 9.5 Hz, 1H), 6.89 (s, 1H), 6.72 (s, 1H), 6.52 (d, *J* = 16.2 Hz, 1H), 5.05 (q, J = 10.7 Hz, 2H), 4.40 – 4.00 (m, 3H), 3.94 – 3.73 (m, 2H), 2.41 – 2.25 (m, 2H), 1.80 – 1.45 (m, 8H), 1.43 – 1.30 (m, 1H), 1.29 – 1.14 (m, 2H), 1.11 (s, 9H), 1.05 (d, *J* = 6.2 Hz, 3H), 0.97 – 0.83 (m, 2H). HRMS (APCI+) m/z calculated for C30H45Cl2N5O7 $^+$ (M + H)$^+$ 658.2769, found 658.2757.

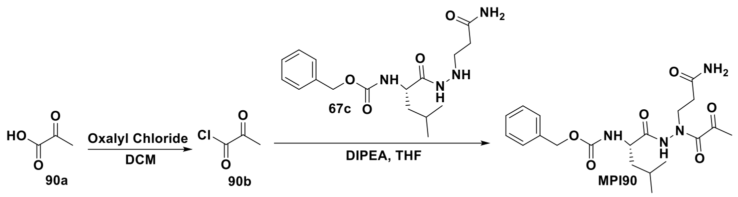


**Scheme 21.** The synthesis of **MPI90**

**Synthesis of 2-oxopropanoyl chloride (90b)**. To a stirred solution of **90a** (200 mg, 2.27 mmol) in DCM (5 mL) at 0 °C was added oxalyl chloride (233 µL, 2.72 mmol) and catalytic amount of DMF. The reaction mixture was stirred at room temperature for 3 h. After completion of reaction, solvent was concentrated in vacuum. The residue was used in the next step without further purification.

**Synthesis of benzyl (S)-(1-(2-(3-amino-3-oxopropyl)-2-(2-oxopropanoyl)hydrazineyl)-4-methyl-1-oxopentan-2-yl)carbamate (MPI90)**. To a stirred solution of **90b** (50 mg, 0.14 mmol) in THF (5 mL) at 0 °C was added DIPEA (37 µL, 0.21 mmol). After 15 min, compound **67c** (18 mg, 0.17 mmol) was added, and the mixture was stirred at room temperature for 3 h. The reaction was quenched with water (5 mL) and extracted with EtOAc (2 × 10 mL). The combined organic layer was washed with brine, dried over $MgSO_4$, and concentrated in vacuum. The residue was then purified with flash chromatography (0-10% MeOH in dichloromethane as the eluent) to afford **MPI90** as a yellow solid (30 mg, 50%). $^1$H NMR (400 MHz, $CDCl_3$) δ 9.36 (s, 1H), 7.27 (m, 5H), 5.84 (d, *J* = 79.9 Hz, 2H), 5.05 (s, 2H), 4.10 (s, 1H), 3.77 (m, 2H), 2.43 (m, 2H), 2.24 (s, 3H), 1.45 (dd, *J* = 18.9, 9.8 Hz, 2H), 1.24 (d, *J* = 19.4 Hz, 1H), 0.86 (d, *J* = 6.6 Hz, 6H). HRMS (ESI+) m/z calculated for C20H28N4O6 $^+$ (M + H) $^+$ 421.2082, found 421.2072.

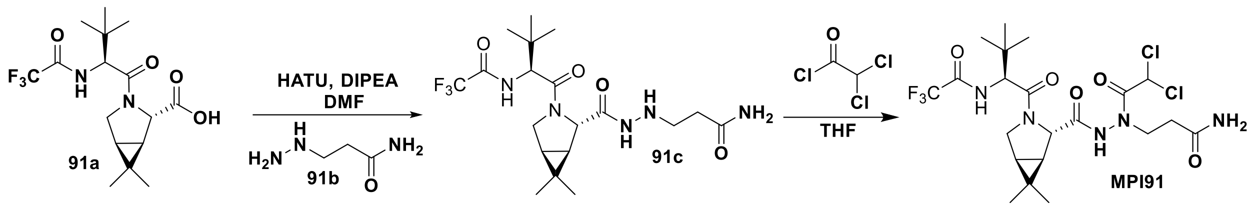


**Scheme 22.** The synthesis of **MPI91**

**Synthesis of 3-(2-((1R,2S,5S)-3-((S)-3,3-dimethyl-2-(2,2,2-trifluoroacetamido)butanoyl)-6,6-dimethyl-3-azabicyclo[3.1.0]hexane-2-carbonyl)hydrazineyl)propenamide (91c)**. To a solution of **91a** (70 mg, 0.2 mmol) in DMF (1 mL) was added compound **91b** (27 mg, 0.25 mmol) and DIPEA (26 mg, 0.2 mmol). The solution was cooled at 0 °C and HATU (95 mg, 0.25 mmol) was added to the solution. The reaction mixture was then allowed to warm up to room temperature and stirred at room temperature overnight. The reaction mixture was the diluted with ethyl acetate (10 mL), washed with saturated $NaHCO_3$ solution (2×10 mL), 1 M HCl (2×10 mL) and saturated brine (10 mL) sequentially. The organic layers were then dried over anhydrous $Na_2SO_4$, concentrated *in vacuo* and the residue was purified with flash chromatography (1-10% methanol in dichloromethane as the eluent) to afford **91c** as white solid. $^1$H NMR (400 MHz, DMSO) δ 9.41 (dd, *J* = 27.6, 7.2 Hz, 2H), 7.31 (s, 1H), 6.76 (s, 1H), 4.97 (q, *J* = 6.1 Hz, 1H), 4.43 (t, *J* = 8.9 Hz, 1H), 4.16 (s, 1H), 3.90 (dd, *J* = 10.3, 5.4 Hz, 1H), 3.68 (d, *J* = 10.3 Hz, 1H), 2.83 (dp, *J* = 12.1, 6.5 Hz, 2H), 2.18 (t, *J* = 7.0 Hz, 2H), 1.52 (dd, *J* = 7.6, 5.2 Hz, 1H), 1.27 (s, 1H), 1.01 (d, *J* = 7.5 Hz, 12H), 0.83 (s, 3H).

**Synthesis of 3-(1-(2,2-dichloroacetyl)-2-((1R,2S,5S)-3-((S)-3,3-dimethyl-2-(2,2,2-trifluoroacetamido)butanoyl)-6,6-dimethyl-3-azabicyclo[3.1.0]hexane-2-carbonyl)hydrazineyl)propenamide (MPI91)**. To a stirred solution of **91c** (30 mg, 0.066 mmol) in THF (5 mL) at 0 °C, dichloroacetyl chloride (7 µL, 0.08) was added dropwise, and the mixture was stirred for a further 30 min at the same temperature. After completion of the reaction, the reaction mixture concentrated in a vacuum. The residue was purified by flash column chromatography (0 to 4% MeOH in ethyl acetate as the eluent)) to give the title compound **MPI91** as white solid (15 mg). $^1$H NMR (400 MHz, $CDCl_3$) δ 9.36 (s, 1H), 7.27 (m, 5H), 5.84 (d, *J* = 79.9 Hz, 2H), 5.05 (s, 2H), 4.10 (s, 1H), 3.77 (m, 2H), 2.43 (m, 2H), 2.24 (s, 3H), 1.45 (dd, *J* = 18.9, 9.8 Hz, 2H), 1.24 (d, *J* = 19.4 Hz, 1H), 0.86 (d, *J* = 6.6 Hz, 6H). HRMS (ESI+) m/z calculated for C21H27Cl2N5O5 $^+$ (M + H) $^+$ 500.1462, found 500.1451.

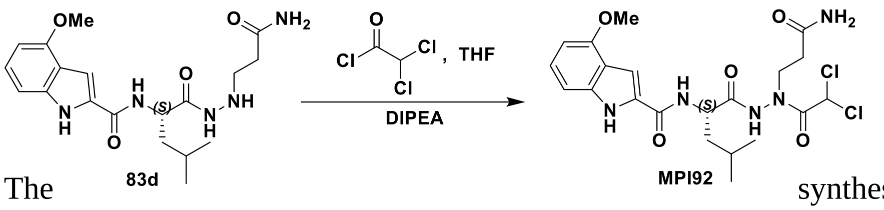


**Scheme 23.** The synthesis of **MPI92**

**Synthesis of (S)-N-(1-(2-(3-amino-3-oxopropyl)-2-(2,2-dichloroacetyl)hydrazineyl)-4-methyl-1-oxopentan-2-yl)-4-methoxy-1H-indole-2-carboxamide (MPI92).** To a solution of **83d** (45 mg, 0.12 mmol, 1.0 equiv) in anhydrous THF at 0 °C was added DIPEA (32 μL, 0.18 mmol, 1.5 equiv) and dichloroacetyl chloride (20 mg, 0.13 mmol, 1.1 equiv) at this temperature. The reaction was quenched by slow addition of $H_2O$ and removed THF on *vacuo*. The mixture was diluted with $H_2O$ and extracted with EtOAc, washed with sat. NaCl, dried over $Na_2SO_4$ and concentrated. The residue was purified by column chromatography (DCM: MeOH = 10:1 v/v) to afford the pure product **MPI92** (27 mg, yield 47%) as a white solid. 1H NMR (400 MHz, Chloroform-*d*) δ 10.80 (s, 1H), 9.98 – 9.55 (m, 1H), 7.14 – 7.00 (m, 2H), 6.98 – 6.87 (m, 1H), 6.84 (d, *J* = 8.1 Hz, 1H), 6.53 – 6.16 (m, 3H), 6.05 (d, *J* = 11.0 Hz, 1H), 5.05 – 4.86 (m, 1H), 4.31 – 3.96 (m, 1H), 3.84 (d, *J* = 9.0 Hz, 3H), 3.58 (s, 1H), 2.49 (t, *J* = 24.1 Hz, 2H), 2.00 (d, *J* = 21.2 Hz, 2H), 1.67 – 1.43 (m, 2H), 1.03 – 0.63 (m, 6H).

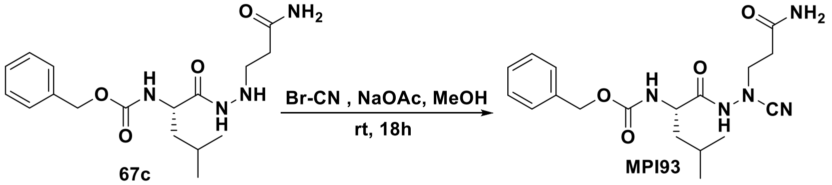


**Scheme 24.** The synthesis of **MPI93**

**Synthesis of benzyl (S)-(1-(2-(3-amino-3-oxopropyl)-2-cyanohydrazineyl)-4-methyl-1-oxopentan-2-yl)carbamate (MPI93).** To a solution **67c** (43 mg, 0.12 mmol) and cyanogen bromide (16 mg, 0.15 mmol) were added sodium acetate (16 mg, 0.18 mmol), dissolved in methanol (3 mL). The reaction was allowed to stir at room temperature for 18 h. The mixture was then poured into water (10 mL) and extracted with ethyl acetate (4×20 mL). The organic layer was washed with aqueous hydrochloric acid 10% v/v (2×20 mL), saturated aqueous $NaHCO_3$ (2×20 mL), brine (2×20 mL) and dried over $Na_2SO_4$. The organic phase was evaporated to dryness and the crude material purified by silica gel column chromatography (1-10% MeOH in $CH_2Cl_2$ as the eluent) to afford **MPI93** as white solid (25 mg, 54%). 1H NMR (400 MHz, Chloroform-*d*) δ 7.29 (d, *J* = 3.5 Hz, 5H), 7.01 (s, 1H), 5.49 (s, 1H), 5.30 (s, 1H), 5.06 (d, *J* = 10.0 Hz, 3H), 4.78 (d, *J* = 7.5 Hz, 1H), 4.04 (d, *J* = 6.9 Hz, 2H), 2.62 (s, 2H), 1.64 (d, *J* = 7.0 Hz, 3H), 0.95 – 0.84 (m, 6H). $^{13}C$ NMR (100 MHz, Chloroform-*d*) δ 171.29, 158.45, 156.96, 155.64, 135.83, 128.64, 128.42, 128.15, 113.42, 67.49, 43.45, 40.91, 32.94, 24.54, 22.52, 21.67.

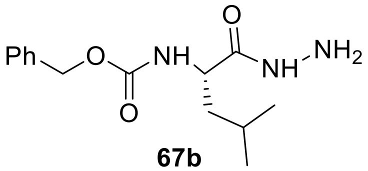

**67b**

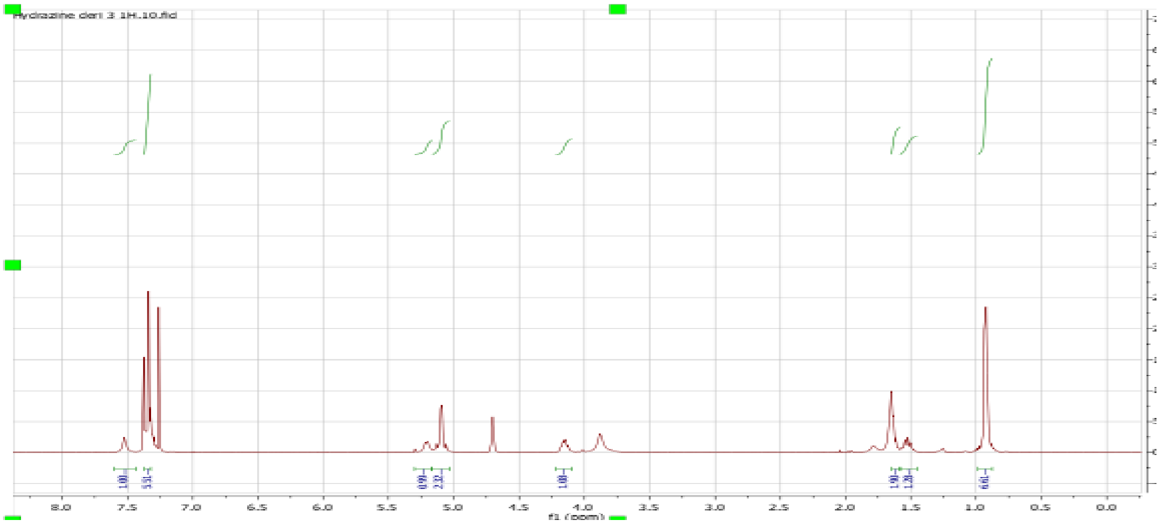

Hydrazine deri 3 1H.10.fid
8.0 7.5 7.0 6.5 6.0 5.5 5.0 4.5 4.0 3.5 3.0 2.5 2.0 1.5 1.0 0.5 0.0
f1 (ppm)

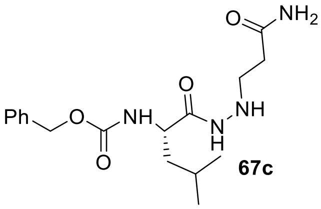
NH2
O
NH
Ph
O
NH
N
H
O
O

**67c**

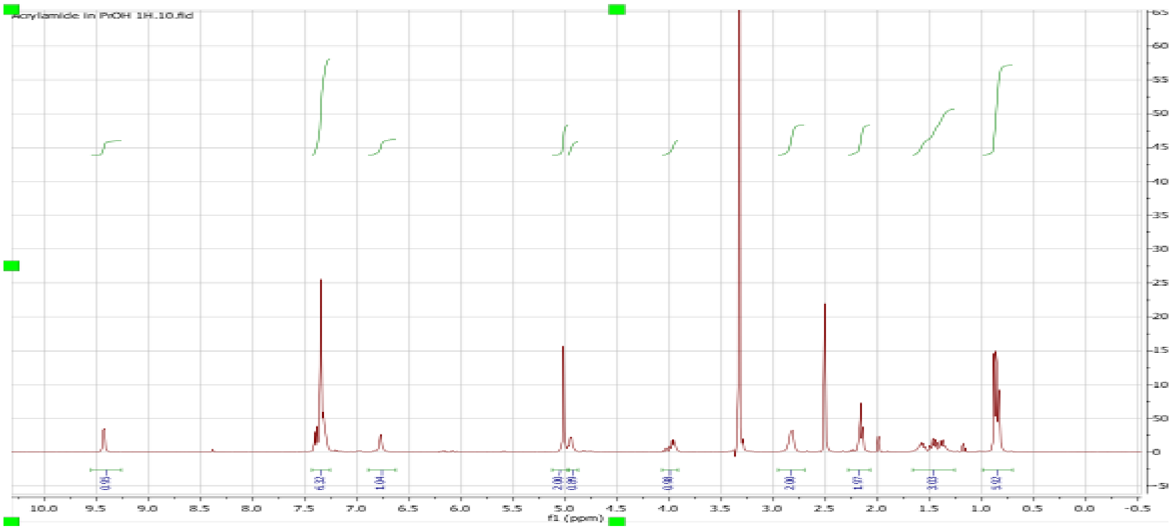
Acrylamide in PrOH 1H.10.fid
f1 (ppm)

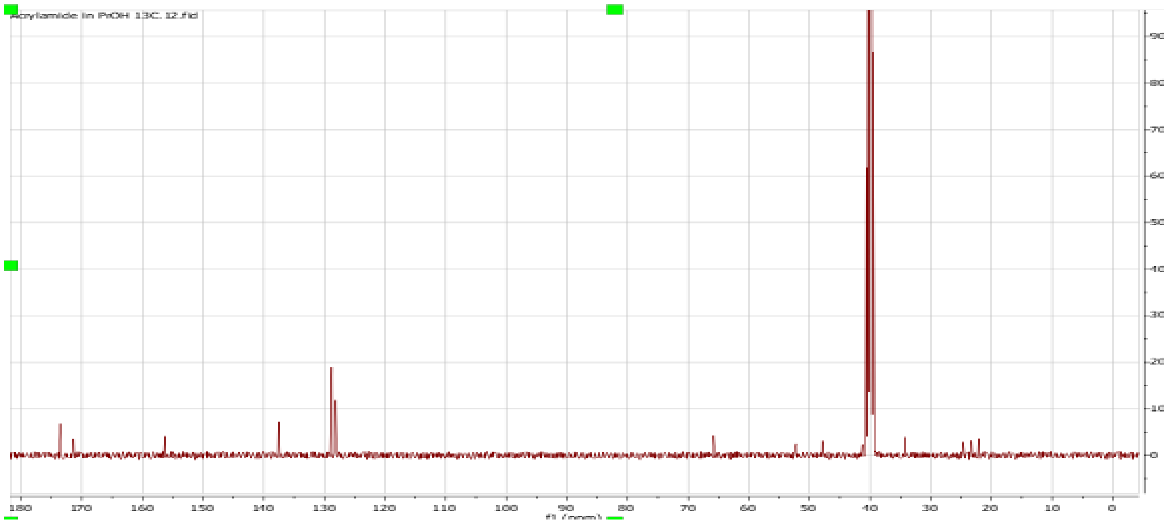
Acrylamide in PrOH 13C.12.fid

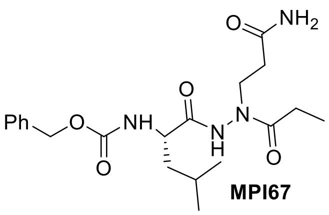

**MPI67**

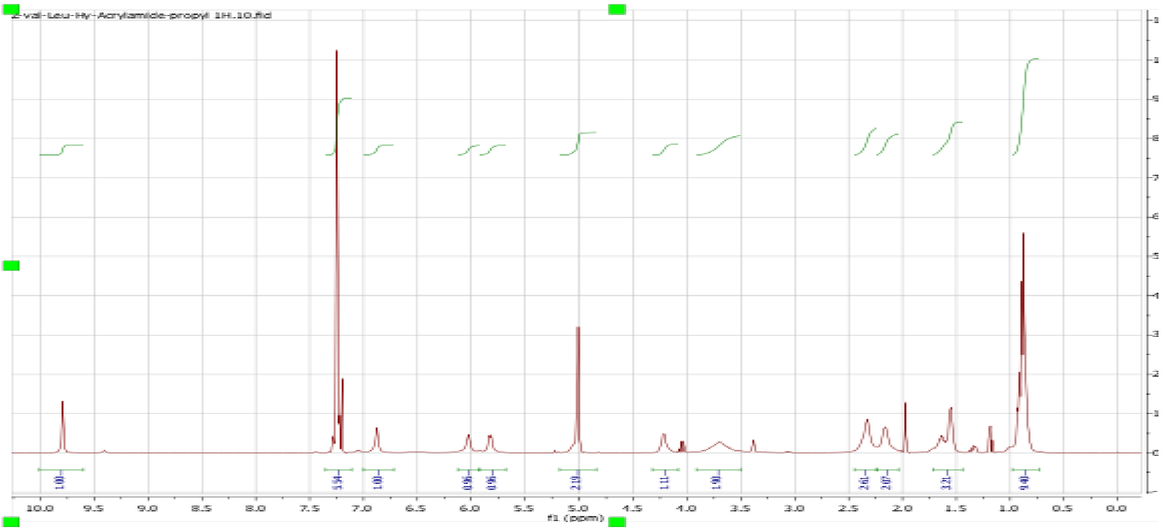
Z-val-Leu-Hy-Acrylamide-propyl 1H.10.fid
f1 (ppm)

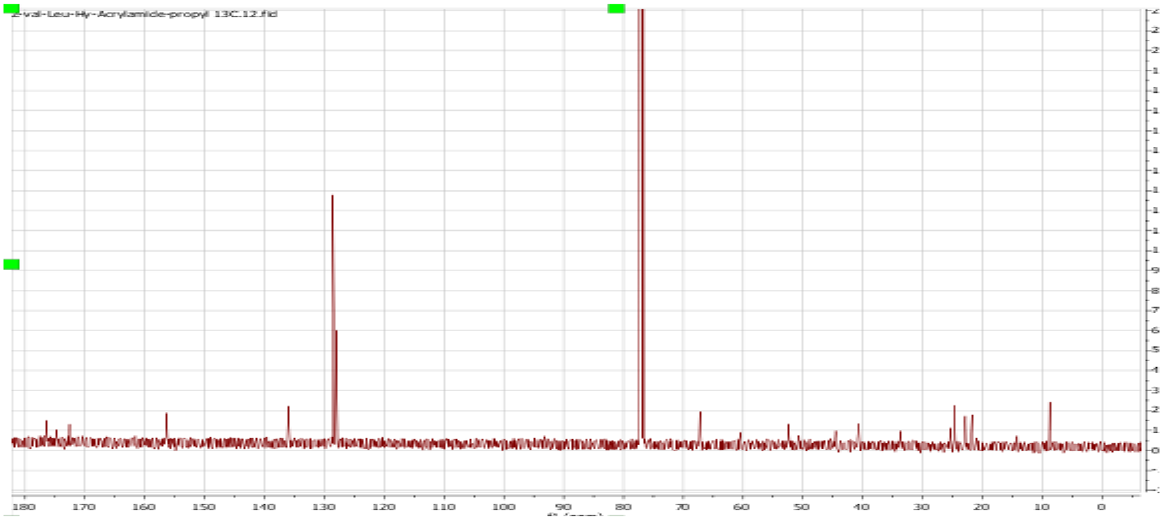
Z-val-Leu-Hy-Acrylamide-propyl 13C.12.fid

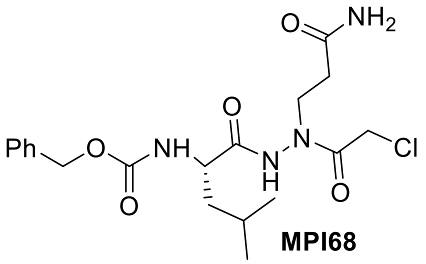
O
NH2
O
Ph
O
NH
N
N
H
Cl
O
O

**MPI68**

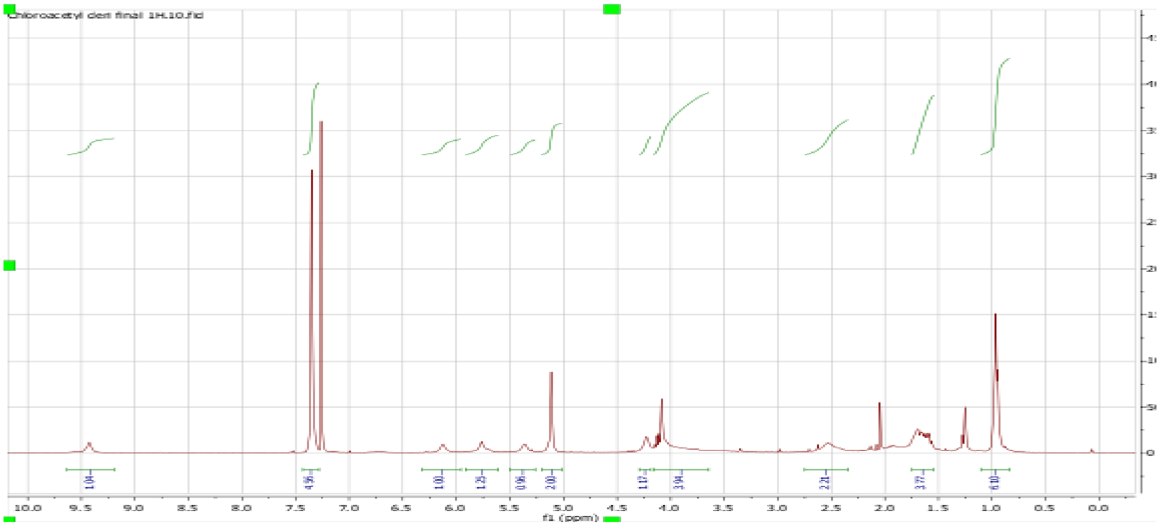
Chloroacetyl deri final 1H.10.fid
1.04
4.96
1.00
1.25
0.96
2.00
1.17
3.94
2.21
3.77
6.10
10.0
9.5
9.0
8.5
8.0
7.5
7.0
6.5
6.0
5.5
5.0
4.5
4.0
3.5
3.0
2.5
2.0
1.5
1.0
0.5
0.0
f1 (ppm)

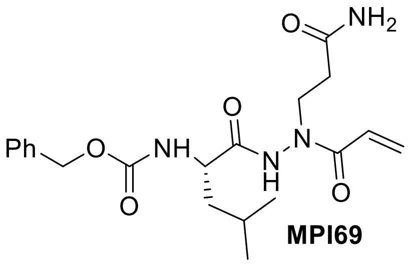
O
NH2
O
Ph
O
NH
N
N
H
O
O

**MPI69**

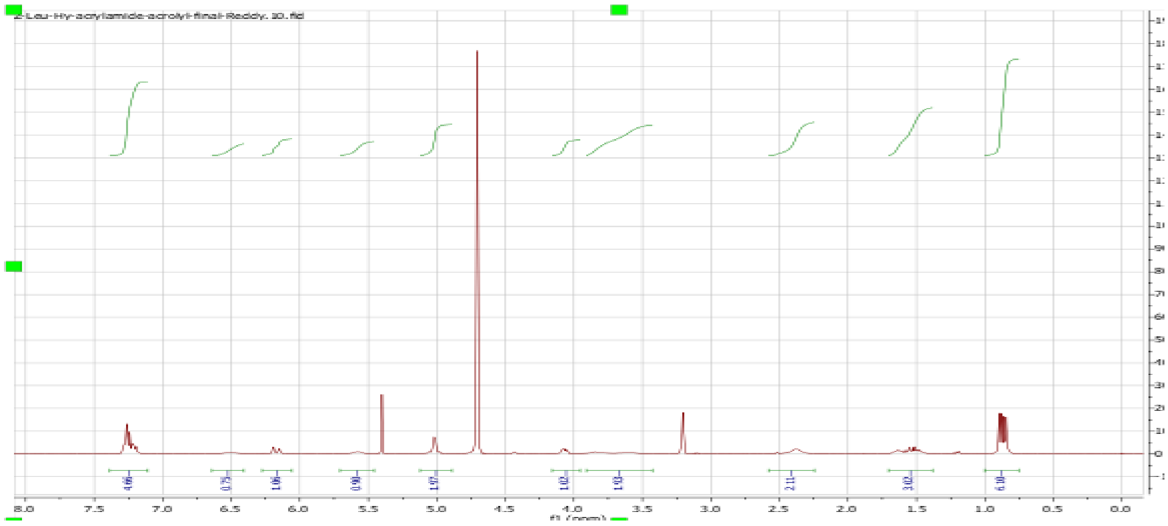
Z-Leu-Hy-acrylamide-acrolyl-final-Reddy.10.fid
f1 (ppm)

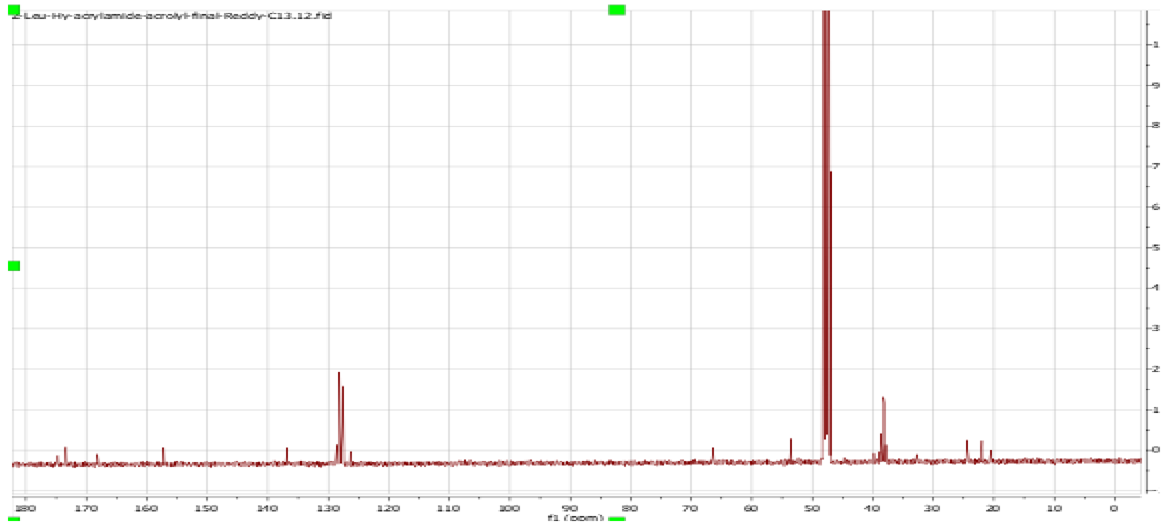
Z-Leu-Hy-acrylamide-acrolyl-final-Reddy-C13.12.fid
f1 (ppm)

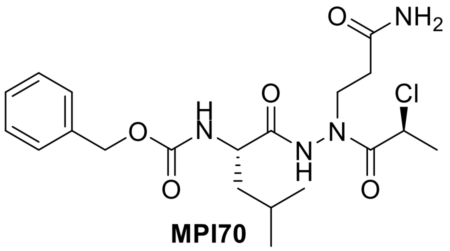

**MPI70**

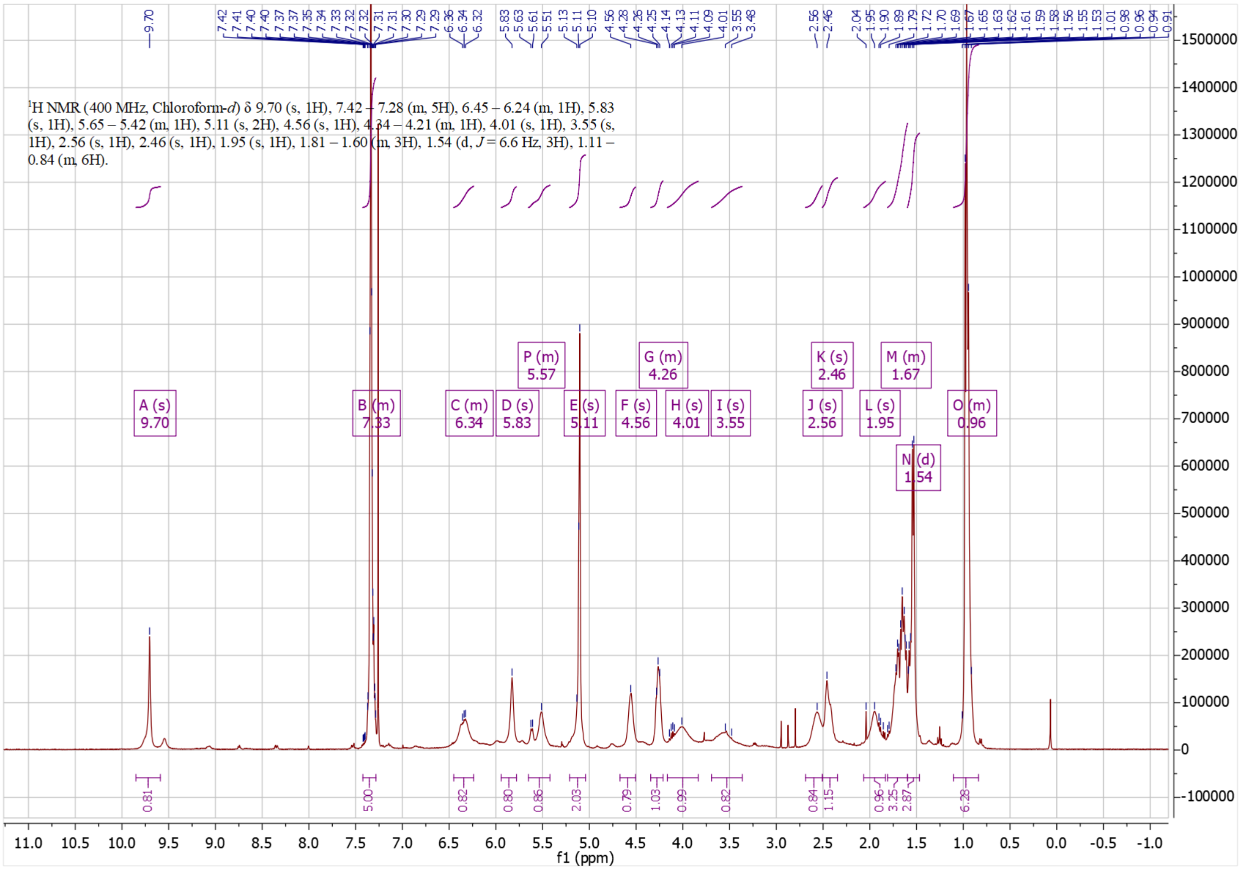

$^{1}$H NMR (400 MHz, Chloroform-*d*) δ 9.70 (s, 1H), 7.42 – 7.28 (m, 5H), 6.45 – 6.24 (m, 1H), 5.83 (s, 1H), 5.65 – 5.42 (m, 1H), 5.11 (s, 2H), 4.56 (s, 1H), 4.34 – 4.21 (m, 1H), 4.01 (s, 1H), 3.55 (s, 1H), 2.56 (s, 1H), 2.46 (s, 1H), 1.95 (s, 1H), 1.81 – 1.60 (m, 3H), 1.54 (d, *J* = 6.6 Hz, 3H), 1.11 – 0.84 (m, 6H).
f1 (ppm)

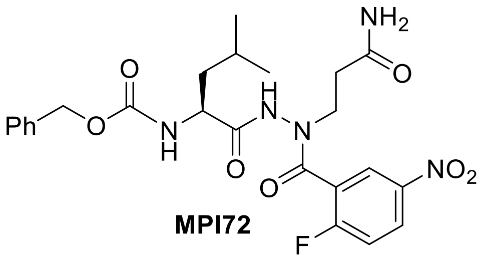

**MPI72**

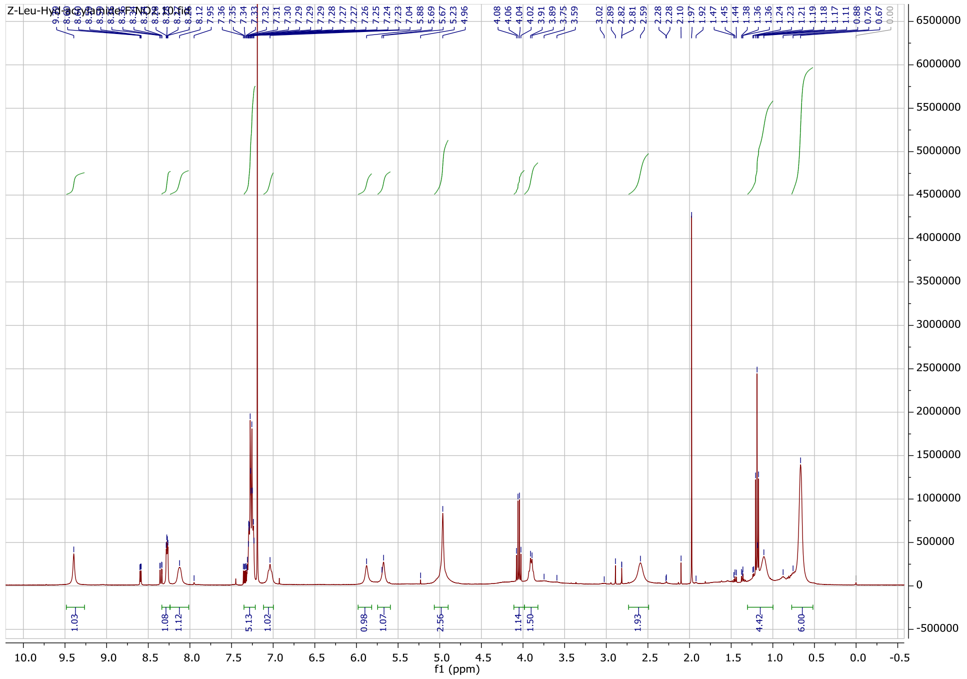

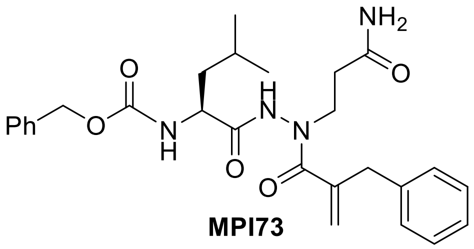

**MPI73**

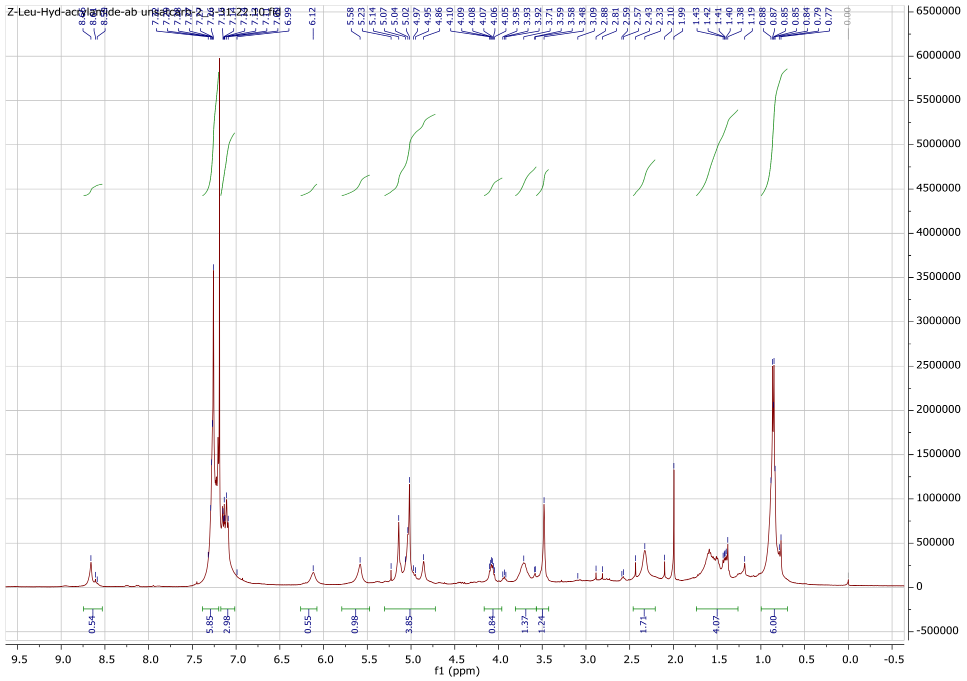

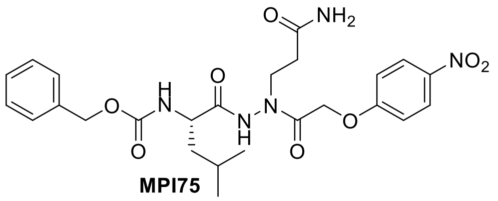

**MPI75**

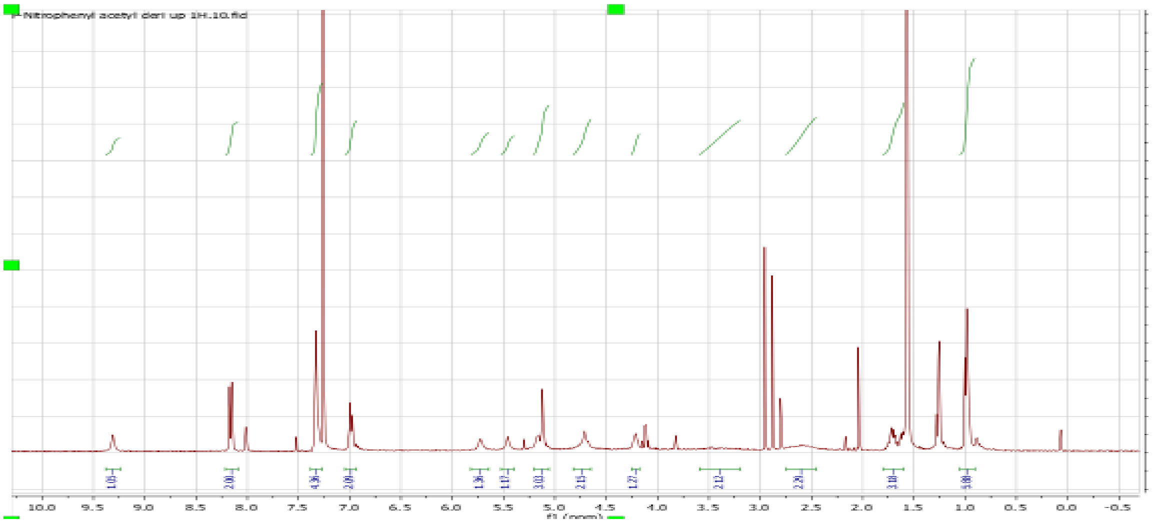
10.0 9.5 9.0 8.5 8.0 7.5 7.0 6.5 6.0 5.5 5.0 4.5 4.0 3.5 3.0 2.5 2.0 1.5 1.0 0.5 0.0 -0.5
f1 (ppm)

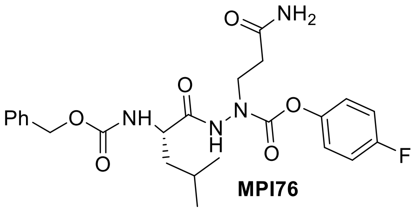
NH2
O
Ph
O
NH
O
O
N
H
N
O
O
F

**MPI76**

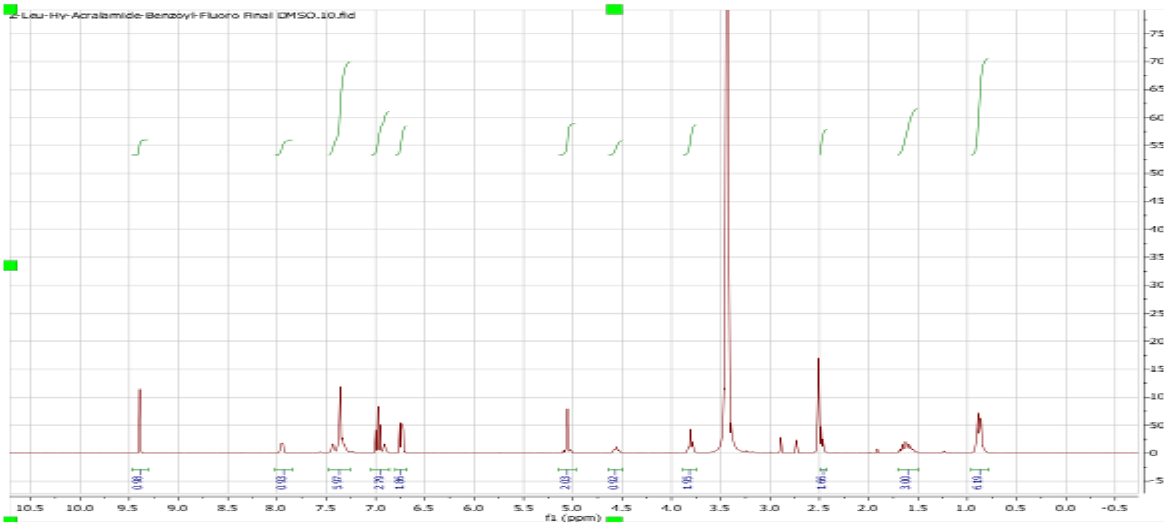
Z-Leu-Hy-Acralamide-Benzoyl-Fluoro Final DMSO.10.fid
f1 (ppm)

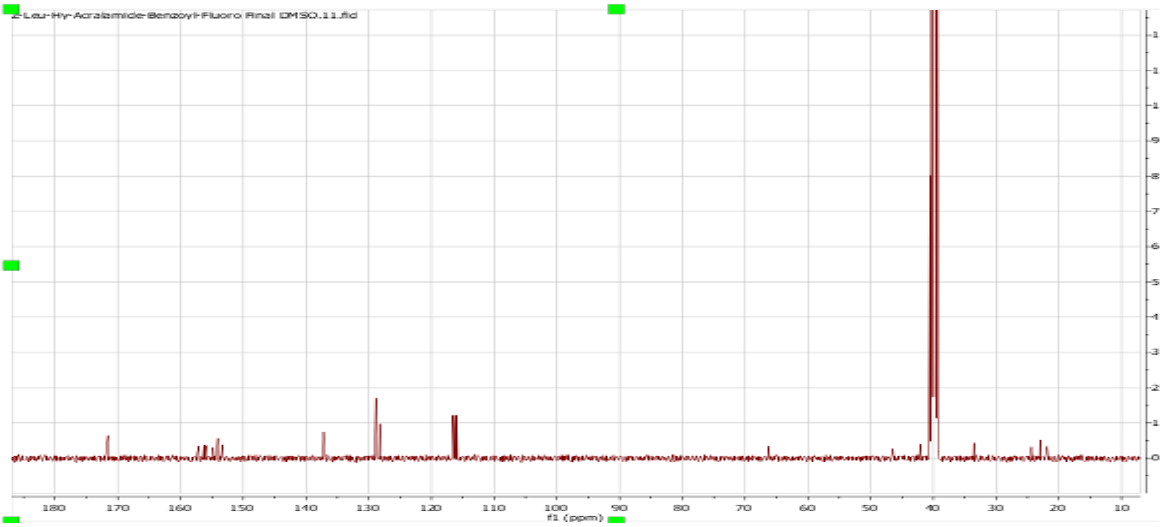
Z-Leu-Hy-Acralamide-Benzoyl-Fluoro Final DMSO.11.fid
f1 (ppm)

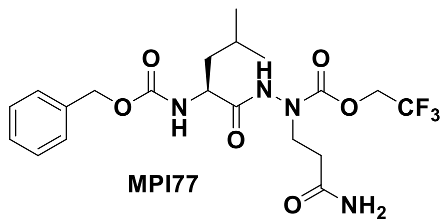
O
N
H
O
H
N
N
O
O
CF3
O
NH2

**MPI77**

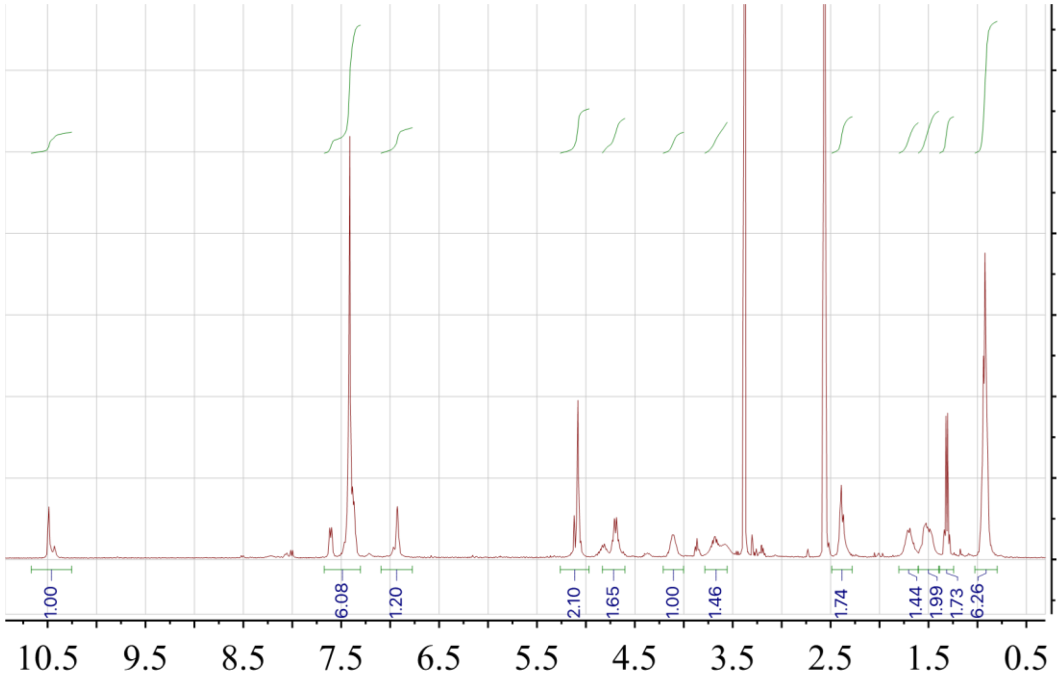
1.00
6.08
1.20
2.10
1.65
1.00
1.46
1.74
1.44
1.99
1.73
6.26
10.5
9.5
8.5
7.5
6.5
5.5
4.5
3.5
2.5
1.5
0.5

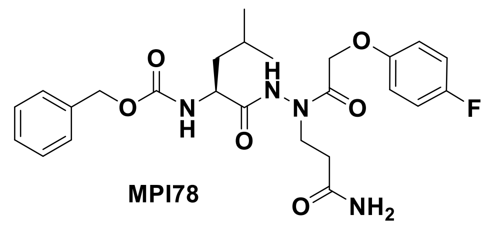

**MPI78**

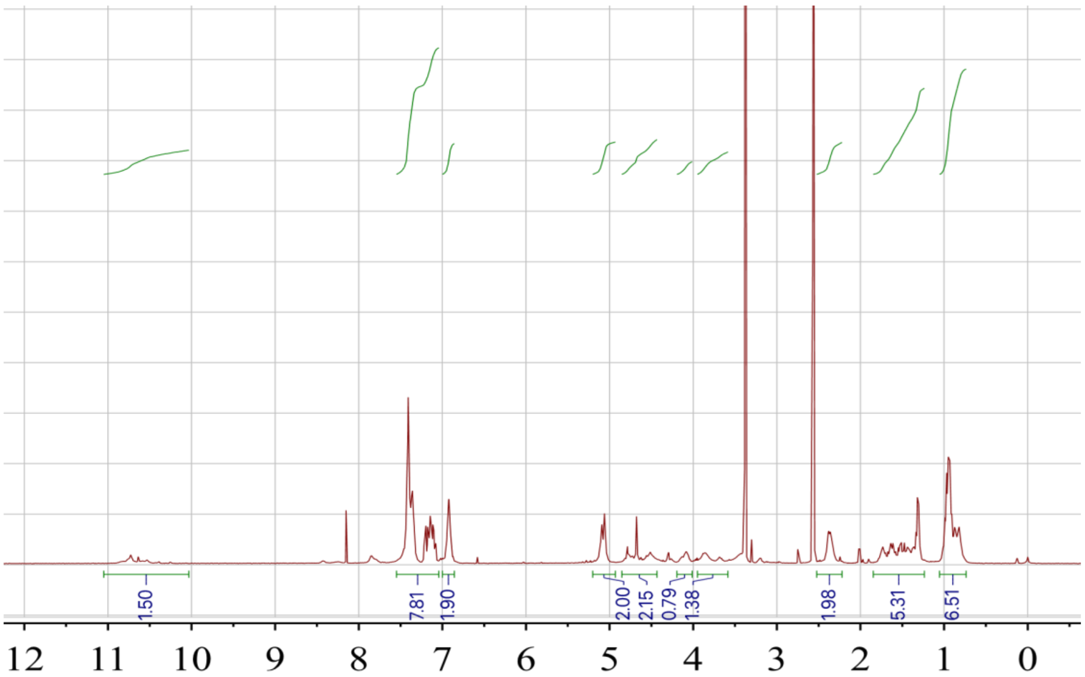

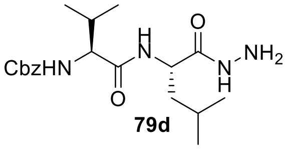
CbzHN
H
N
O
O
N
H
NH2

**79d**

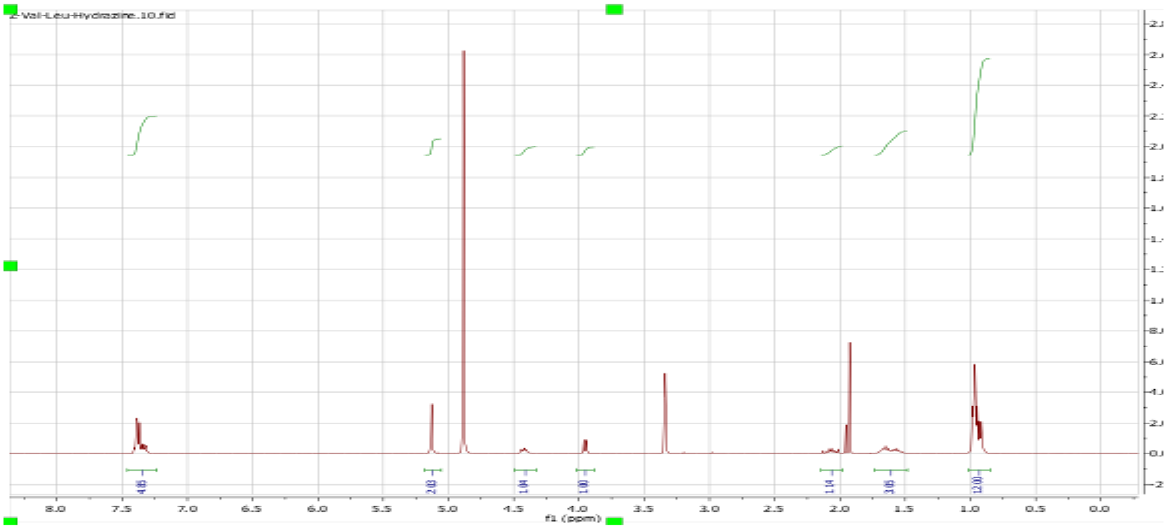
Z-Val-Leu-Hydrazine.10.fid
8.0
7.5
7.0
6.5
6.0
5.5
5.0
4.5
4.0
3.5
3.0
2.5
2.0
1.5
1.0
0.5
0.0
f1 (ppm)

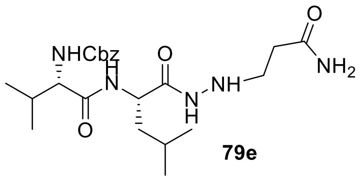

NHCbz
H
N
O
O
N
H
NH
O
NH2


**79e**

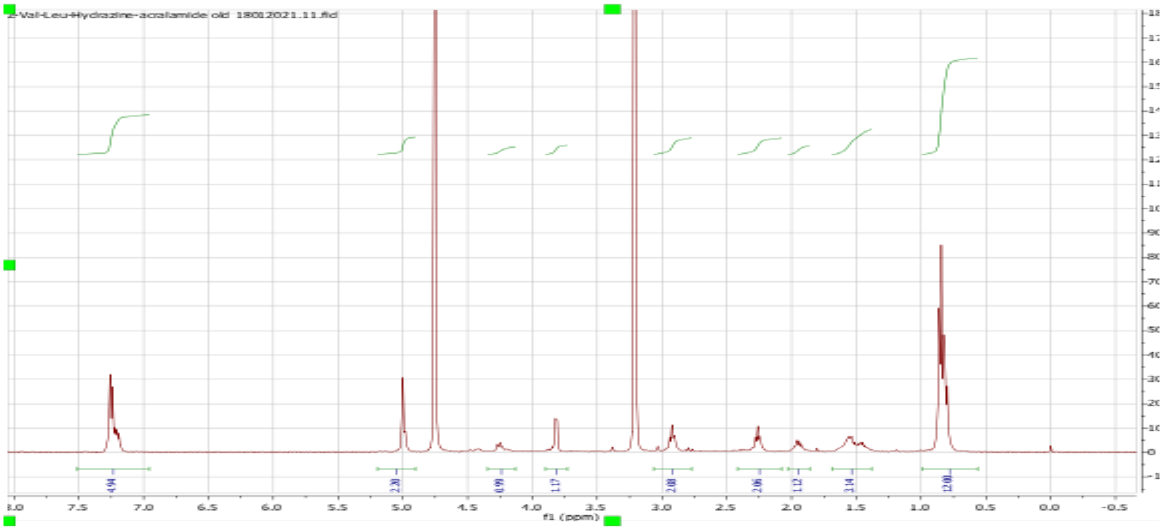

Z-Val-Leu-Hydrazine-acrylamide old 18012021.11.fid
4.94
2.30
0.99
1.17
2.00
2.06
1.12
3.14
12.00
8.0 7.5 7.0 6.5 6.0 5.5 5.0 4.5 4.0 3.5 3.0 2.5 2.0 1.5 1.0 0.5 0.0 -0.5
f1 (ppm)

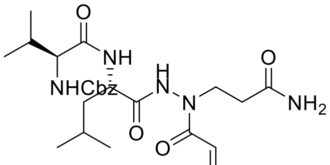
O
NH
H
N
N
O
NHCbz
NH2
O
O

**MPI79**

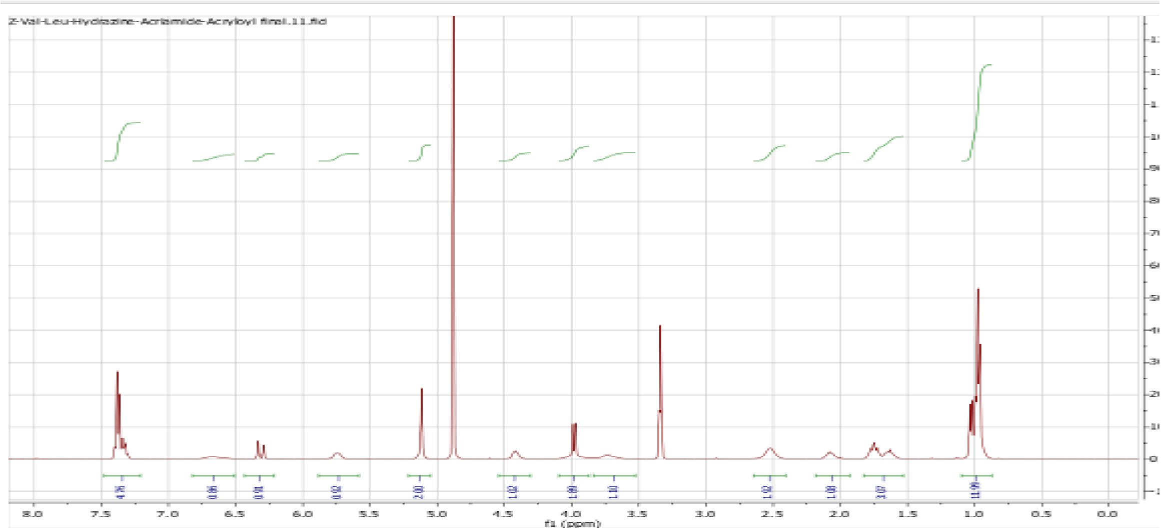
Z-Val-Leu-Hydrazine-Acrlamide-Acryloyl final.11.fid
4.76
0.86
0.91
0.92
2.00
1.02
1.09
1.10
1.02
1.08
3.07
11.99
8.0
7.5
7.0
6.5
6.0
5.5
5.0
4.5
4.0
3.5
3.0
2.5
2.0
1.5
1.0
0.5
0.0
f1 (ppm)

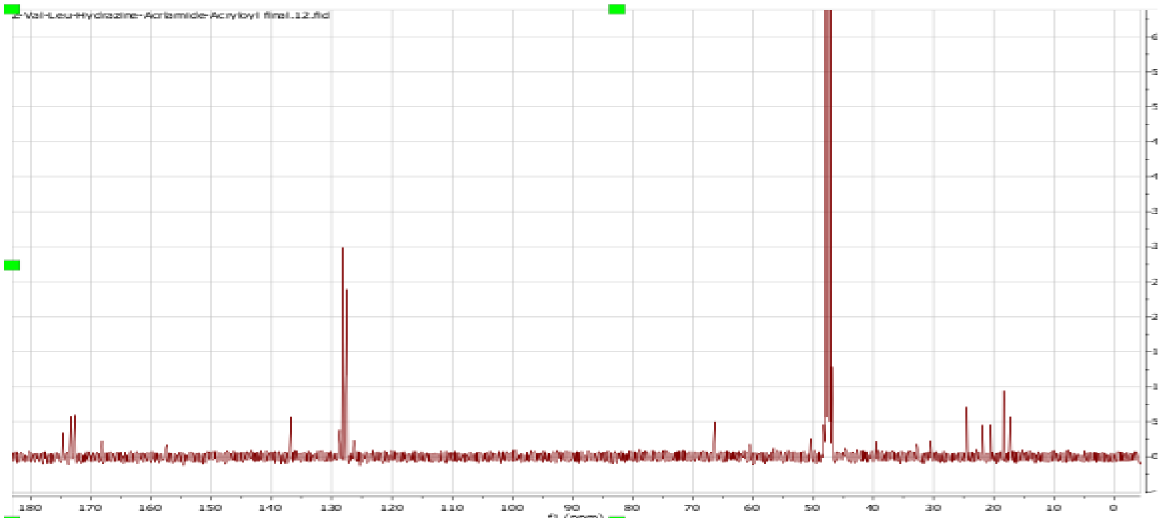
Z-Val-Leu-Hydrazine-Acrlamide-Acryloyl final.12.fid
180
170
160
150
140
130
120
110
100
90
80
70
60
50
40
30
20
10
0

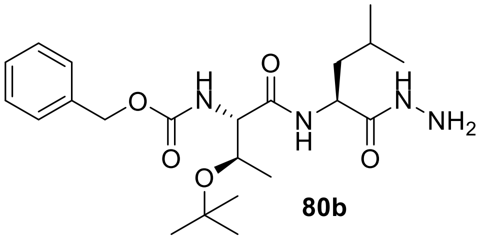

**80b**

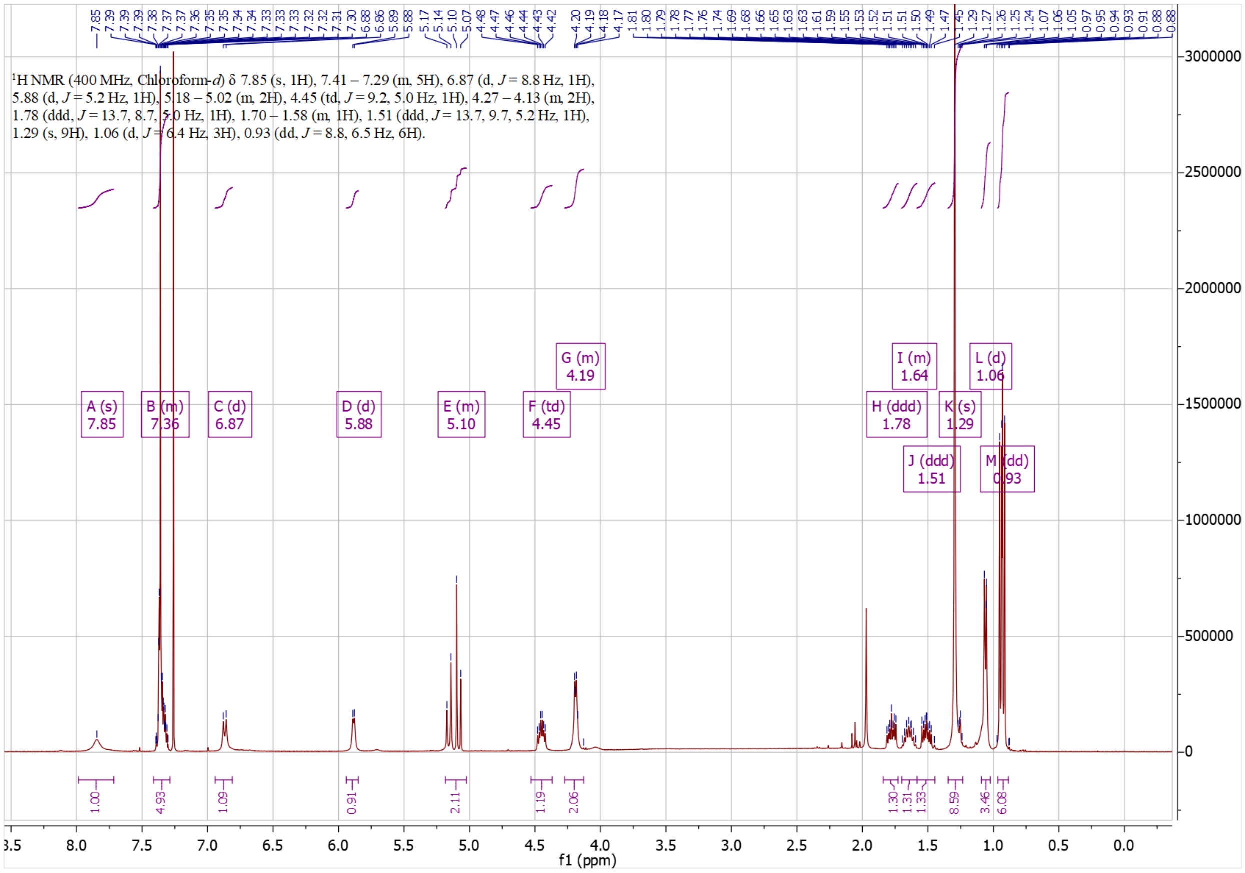

1H NMR (400 MHz, Chloroform-d) δ 7.85 (s, 1H), 7.41 – 7.29 (m, 5H), 6.87 (d, J = 8.8 Hz, 1H), 5.88 (d, J = 5.2 Hz, 1H), 5.18 – 5.02 (m, 2H), 4.45 (td, J = 9.2, 5.0 Hz, 1H), 4.27 – 4.13 (m, 2H), 1.78 (ddd, J = 13.7, 8.7, 5.0 Hz, 1H), 1.70 – 1.58 (m, 1H), 1.51 (ddd, J = 13.7, 9.7, 5.2 Hz, 1H), 1.29 (s, 9H), 1.06 (d, J = 6.4 Hz, 3H), 0.93 (dd, J = 8.8, 6.5 Hz, 6H).
f1 (ppm)

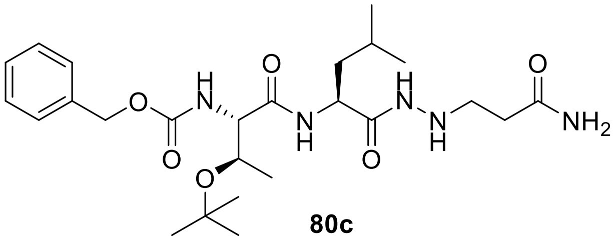

**80c**

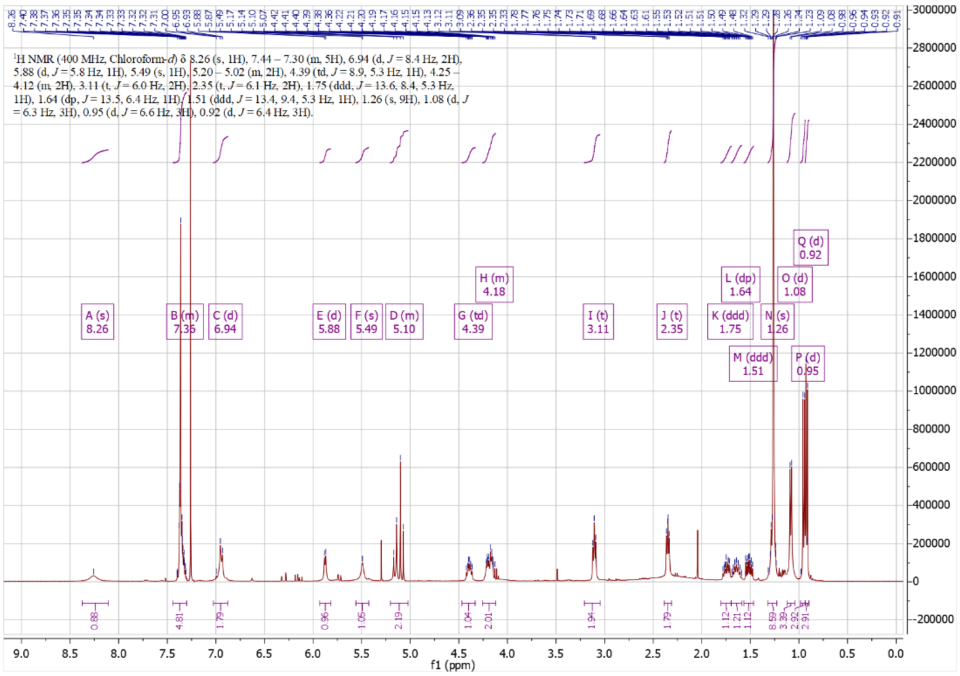

¹H NMR (400 MHz, Chloroform-*d*) δ 8.26 (s, 1H), 7.44 – 7.30 (m, 5H), 6.94 (d, *J* = 8.4 Hz, 2H), 5.88 (d, *J* = 5.8 Hz, 1H), 5.49 (s, 1H), 5.20 – 5.02 (m, 2H), 4.39 (td, *J* = 8.9, 5.3 Hz, 1H), 4.25 – 4.12 (m, 2H), 3.11 (t, *J* = 6.0 Hz, 2H), 2.35 (t, *J* = 6.1 Hz, 2H), 1.75 (ddd, *J* = 13.6, 8.4, 5.3 Hz, 1H), 1.64 (dp, *J* = 13.5, 6.4 Hz, 1H), 1.51 (ddd, *J* = 13.4, 9.4, 5.3 Hz, 1H), 1.26 (s, 9H), 1.08 (d, *J* = 6.3 Hz, 3H), 0.95 (d, *J* = 6.6 Hz, 3H), 0.92 (d, *J* = 6.4 Hz, 3H).
f1 (ppm)

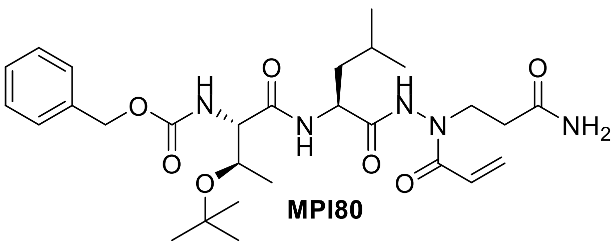
O
N
H
O
N
H
O
N
H
N
O
NH2
O
O
O

**MPI80**

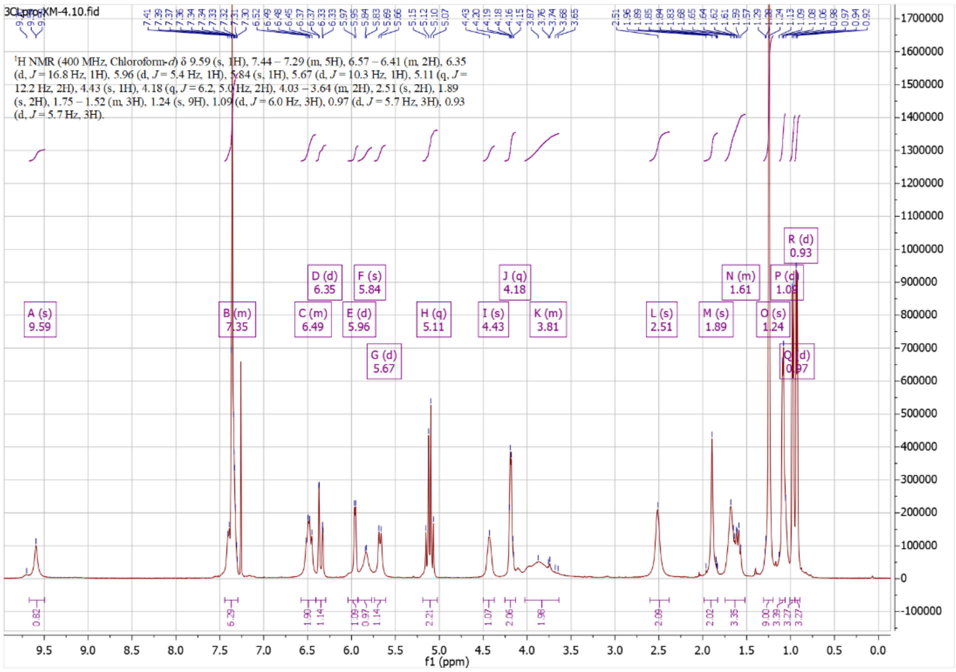
3CLpro-XM-4.10.fid
1H NMR (400 MHz, Chloroform-d) δ 9.59 (s, 1H), 7.44 – 7.29 (m, 5H), 6.57 – 6.41 (m, 2H), 6.35 (d, J = 16.8 Hz, 1H), 5.96 (d, J = 5.4 Hz, 1H), 5.84 (s, 1H), 5.67 (d, J = 10.3 Hz, 1H), 5.11 (q, J = 12.2 Hz, 2H), 4.43 (s, 1H), 4.18 (q, J = 6.2, 5.0 Hz, 2H), 4.03 – 3.64 (m, 2H), 2.51 (s, 2H), 1.89 (s, 2H), 1.75 – 1.52 (m, 3H), 1.24 (s, 9H), 1.09 (d, J = 6.0 Hz, 3H), 0.97 (d, J = 5.7 Hz, 3H), 0.93 (d, J = 5.7 Hz, 3H).
f1 (ppm)

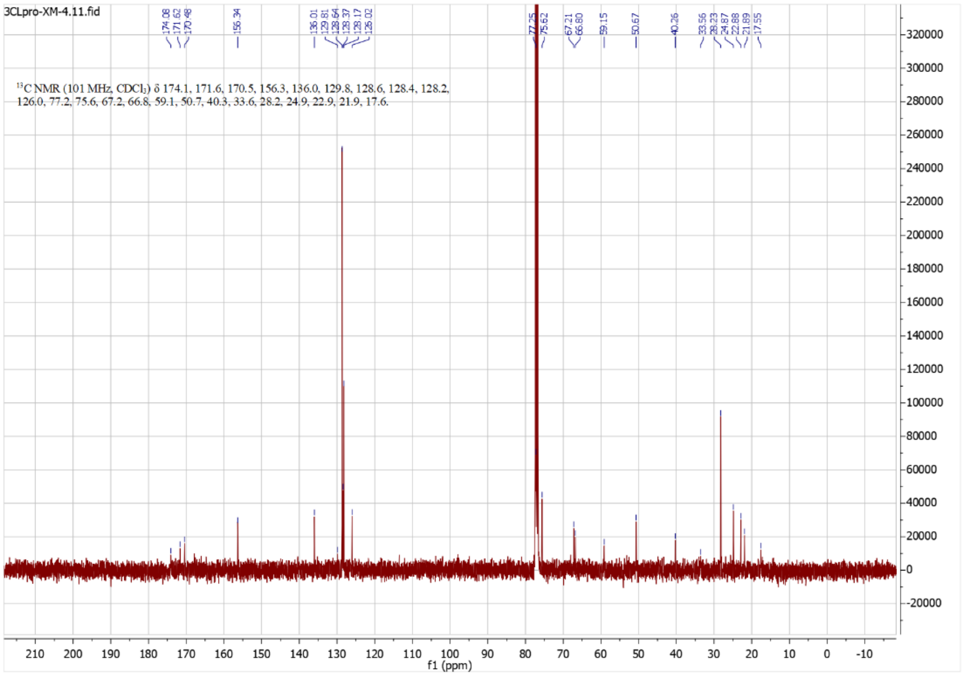
3CLpro-XM-4.11.fid
13C NMR (101 MHz, CDCl3) δ 174.1, 171.6, 170.5, 156.3, 136.0, 129.8, 128.6, 128.4, 128.2, 126.0, 77.2, 75.6, 67.2, 66.8, 59.1, 50.7, 40.3, 33.6, 28.2, 24.9, 22.9, 21.9, 17.6.
f1 (ppm)

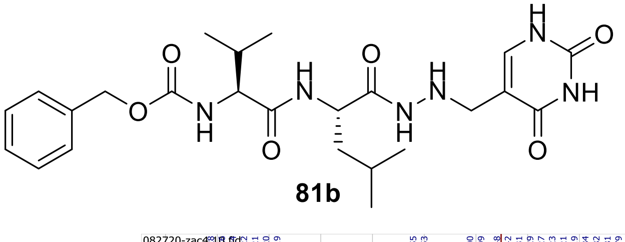

**81b**

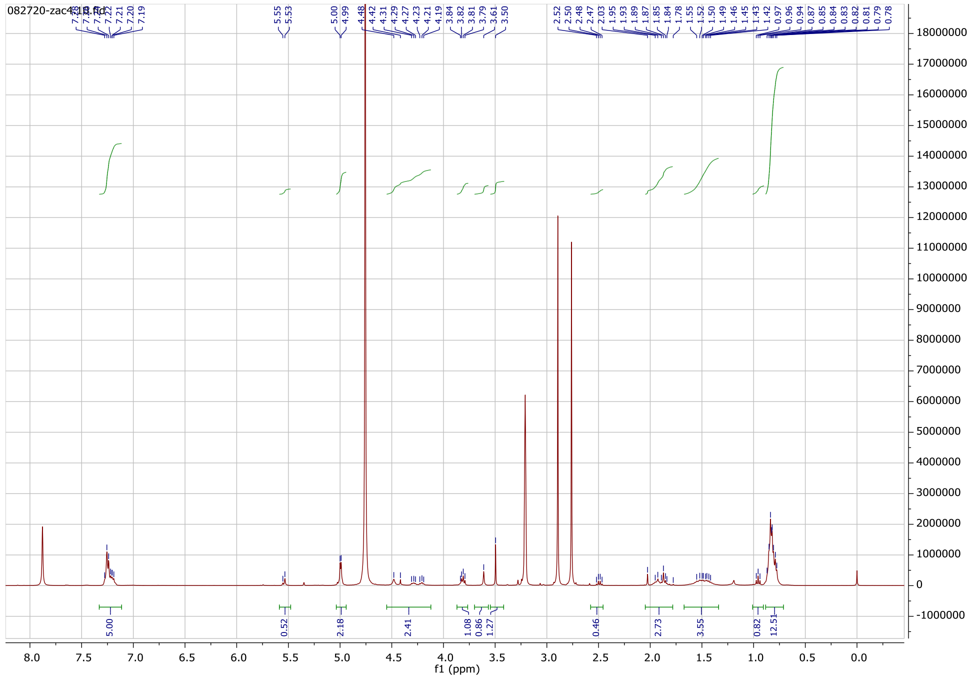

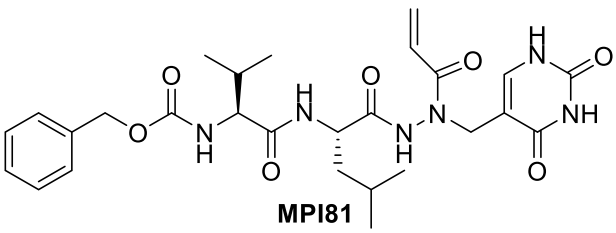

**MPI81**

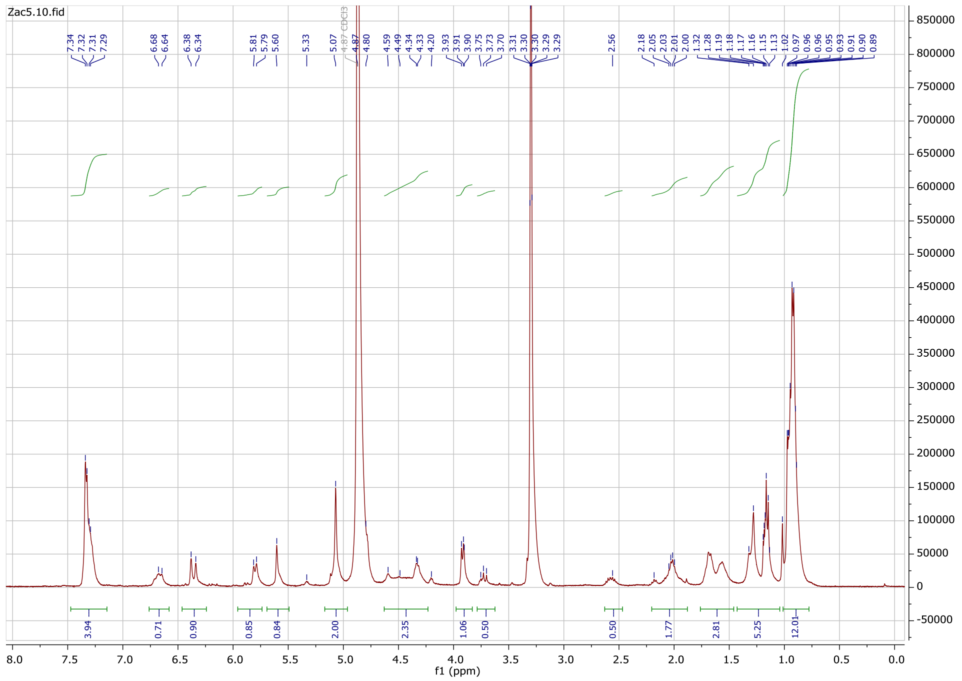
Zac5.10.fid
7.34 7.32 7.31 7.29
6.68 6.64
6.38 6.34
5.81 5.79 5.60
5.33
5.07
4.87 CDCl3
4.80
4.59 4.49 4.34 4.33 4.20
3.93 3.91 3.90 3.75 3.73 3.70
3.31 3.30 3.30 3.29 3.29
2.56
2.18 2.05 2.03 2.01 2.00
1.32 1.28 1.19 1.18 1.17 1.16 1.15 1.13
1.02 0.97 0.96 0.96 0.95 0.93 0.91 0.90 0.89
3.94
0.71
0.90
0.85
0.84
2.00
2.35
1.06
0.50
0.50
1.77
2.81
5.25
12.01
f1 (ppm)

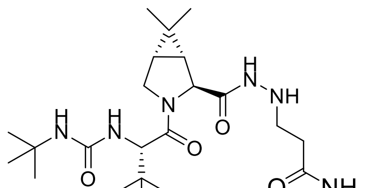

H
N
H
N
H
N
O
O
N
H
NH
O
O
NH2


**82c**

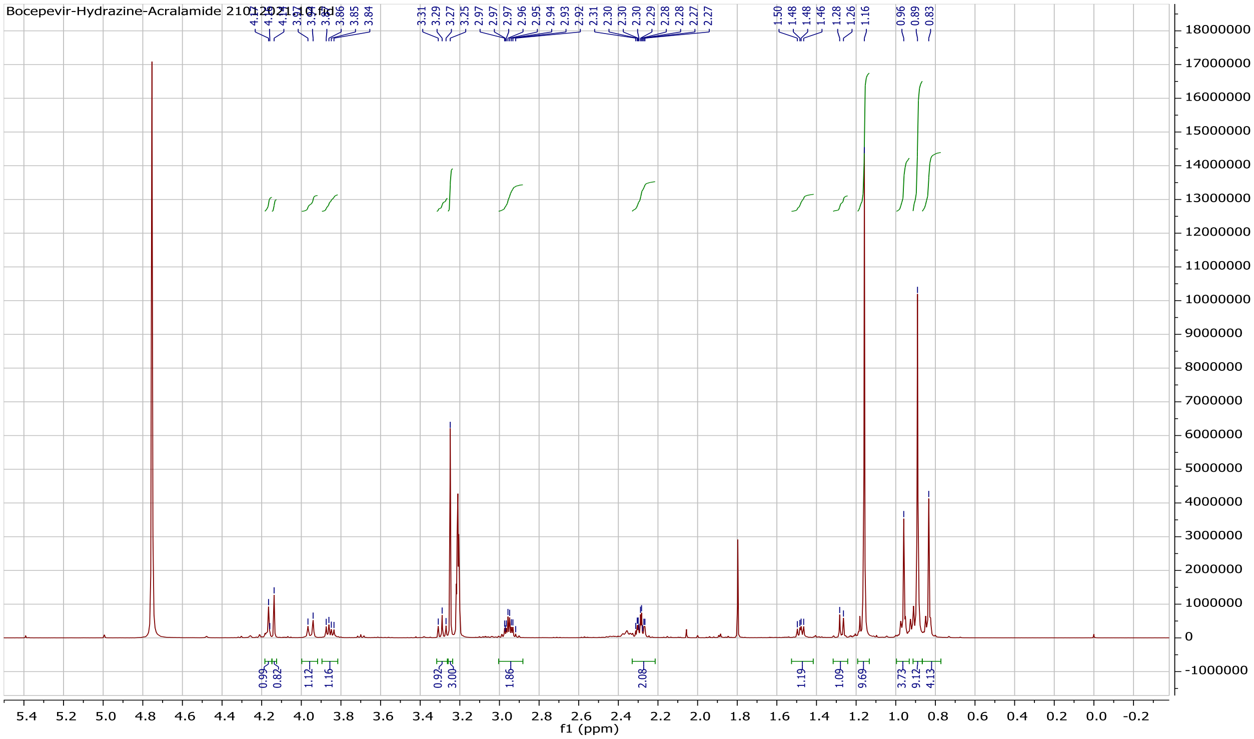

Bocepevir-Hydrazine-Acralamide
3.86
3.85
3.84
3.31
3.29
3.27
3.25
2.97
2.97
2.96
2.95
2.94
2.93
2.92
2.31
2.30
2.30
2.30
2.29
2.28
2.28
2.27
2.27
1.50
1.48
1.48
1.46
1.28
1.26
1.16
0.96
0.89
0.83
0.99
0.82
1.12
1.16
0.92
3.00
1.86
2.08
1.19
1.09
9.69
3.73
9.12
4.13
f1 (ppm)


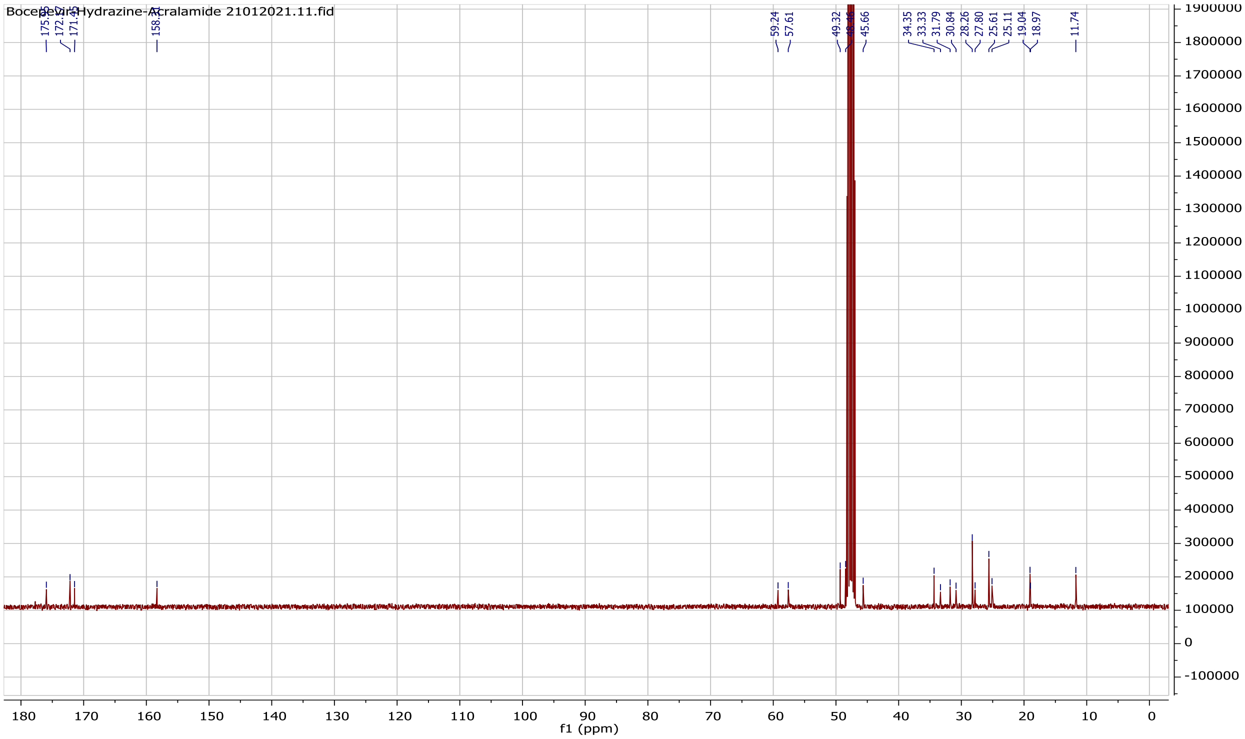

Bocepevir-Hydrazine-Acralamide 21012021.11.fid
59.24
57.61
49.32
45.66
34.35
33.33
31.79
30.84
28.26
27.80
25.61
25.11
19.04
18.97
11.74
f1 (ppm)

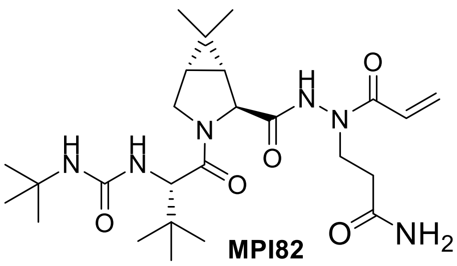

**MPI82**

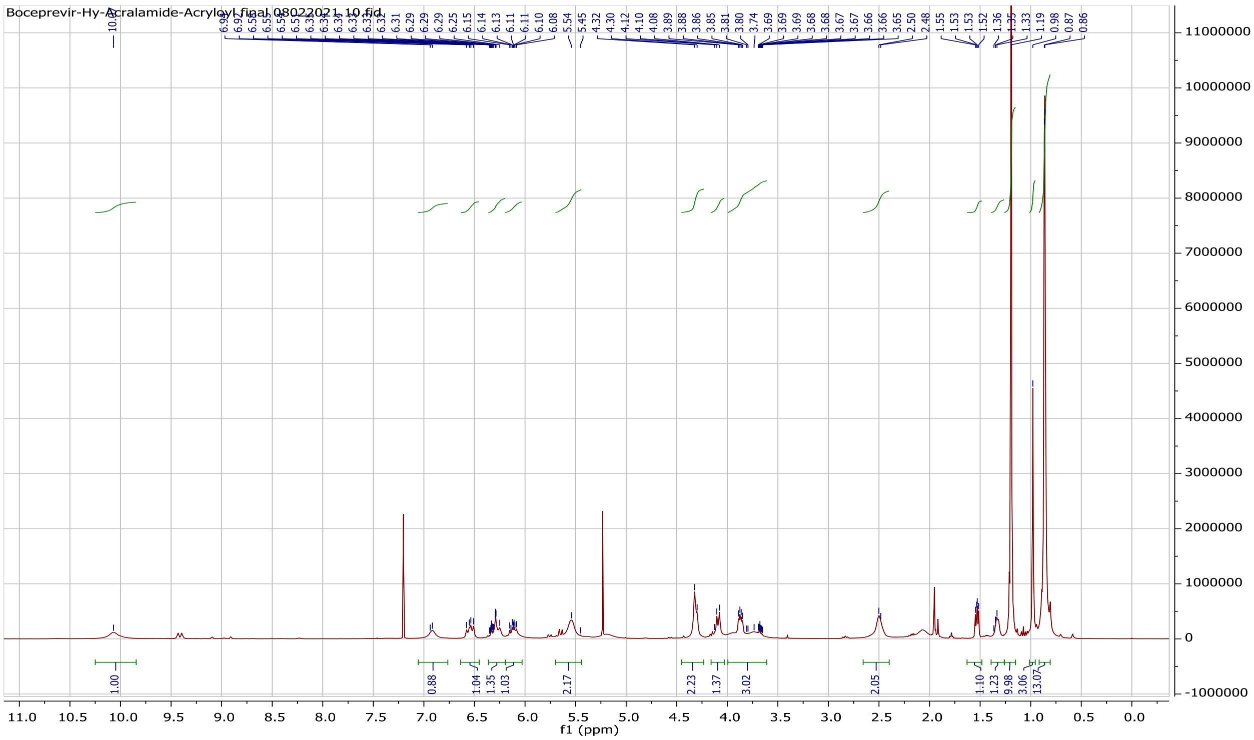
Boceprevir-Hy-Acralamide-Acryloyl final 08022021.10.fid
f1 (ppm)

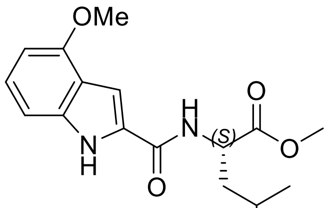
OMe
H
N
(S)
O
O
N
H
O

**83b**

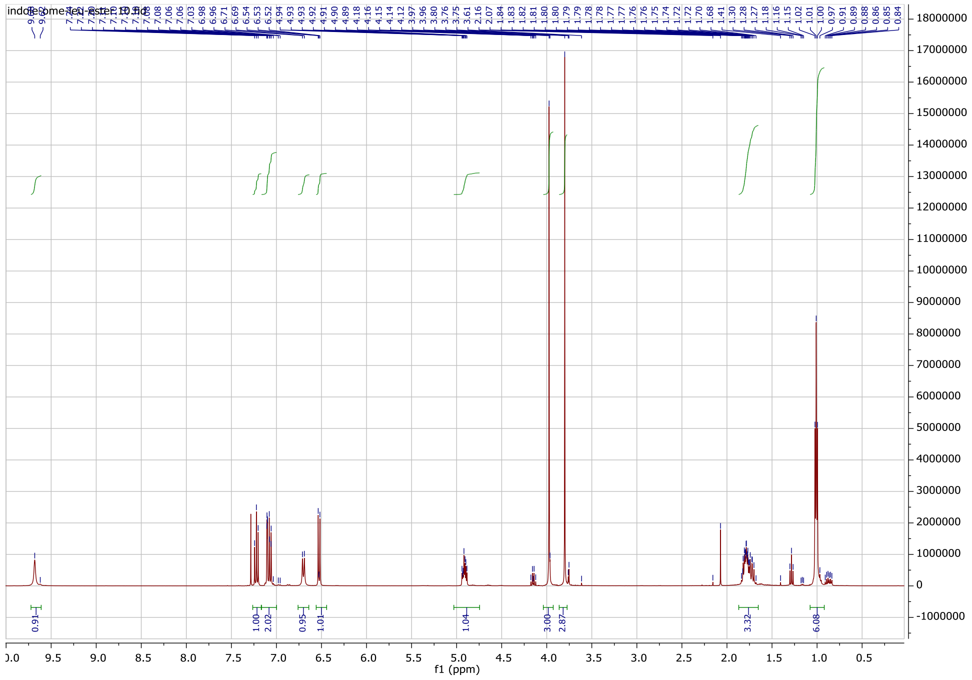
indole-ome-leu-ester.10.fid
0.91
1.00
2.02
0.95
1.01
1.04
3.00
2.87
3.32
6.08
f1 (ppm)

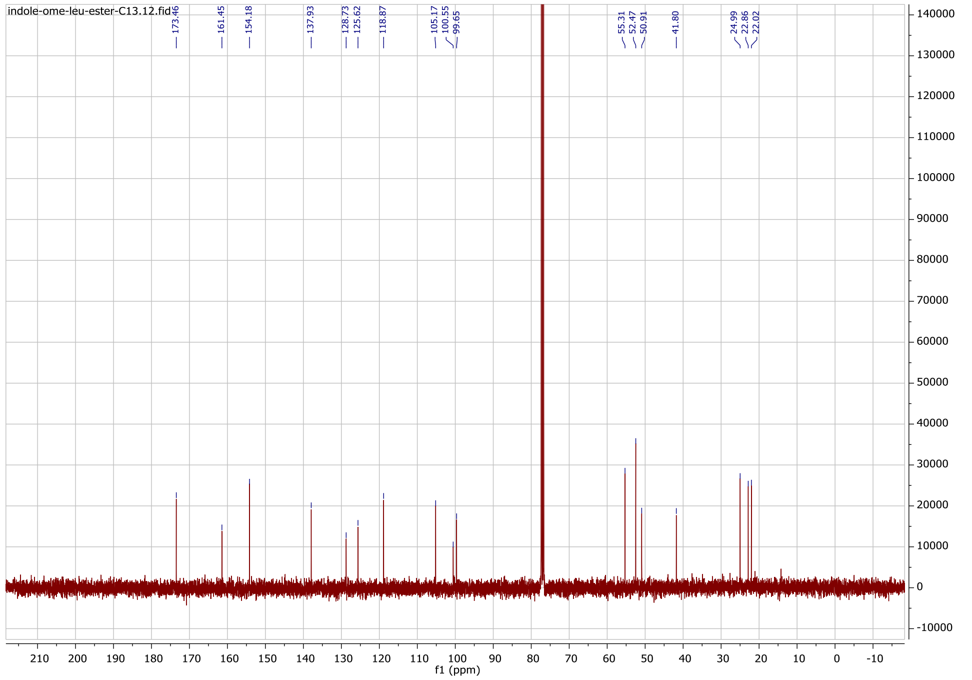
indole-ome-leu-ester-C13.12.fid
173.46
161.45
154.18
137.93
128.73
125.62
118.87
105.17
100.55
99.65
55.31
52.47
50.91
41.80
24.99
22.86
22.02
f1 (ppm)

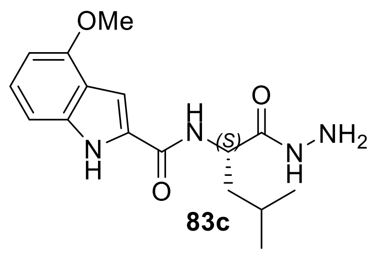

OMe
NH2
83c


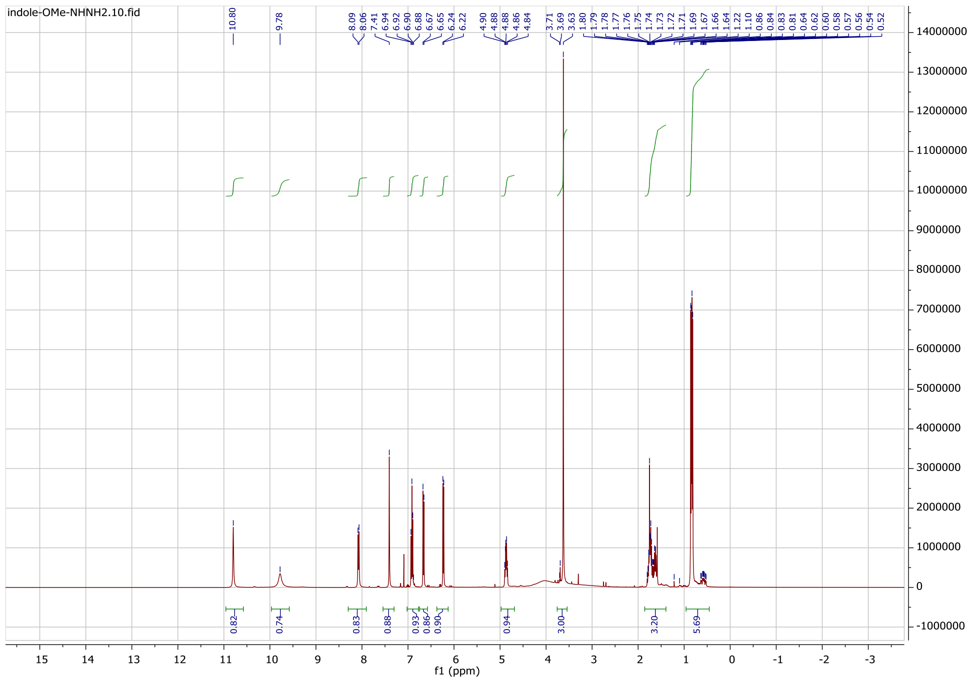

indole-OMe-NHNH2.10.fid
f1 (ppm)


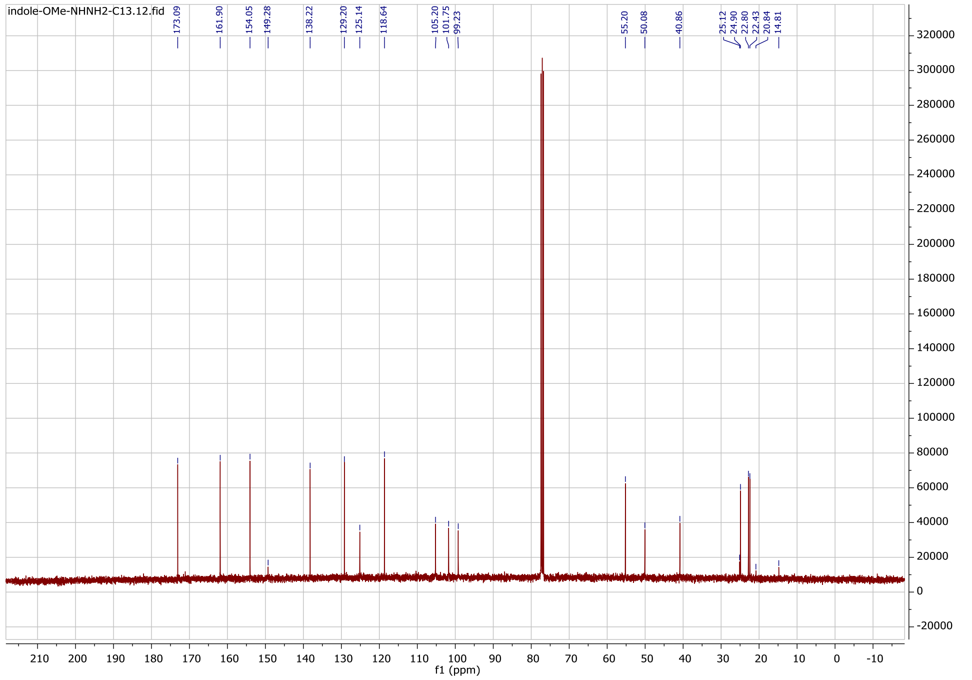

indole-OMe-NHNH2-C13.12.fid
f1 (ppm)

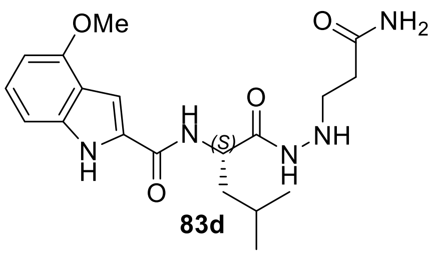

**83d**

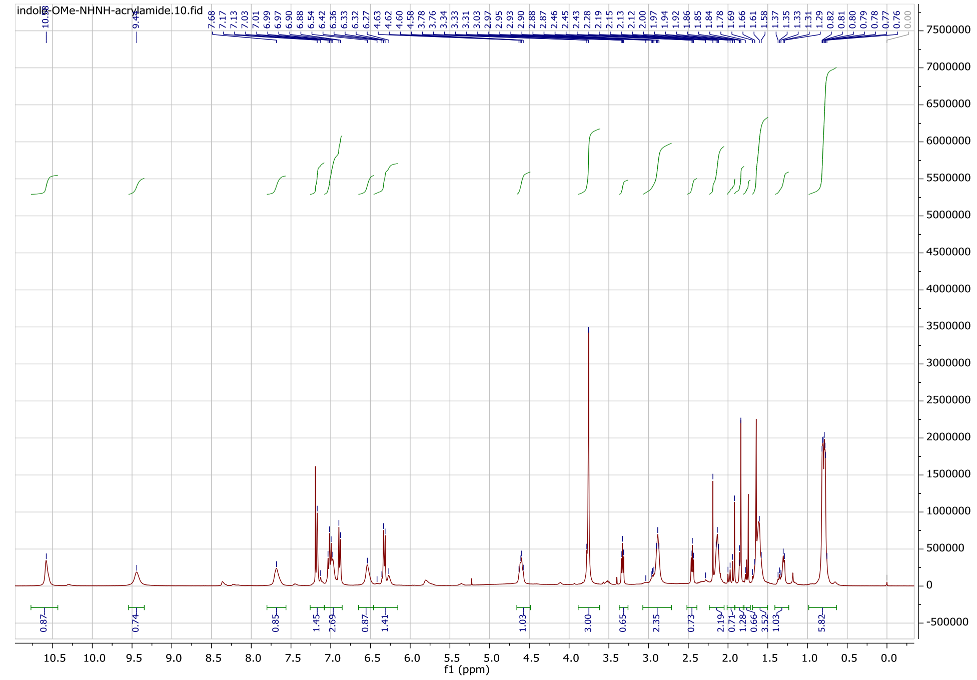

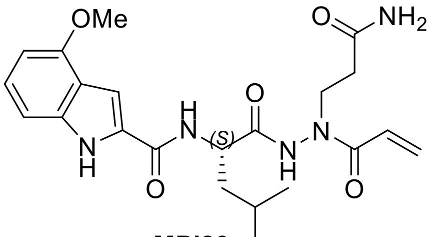

**MPI83**

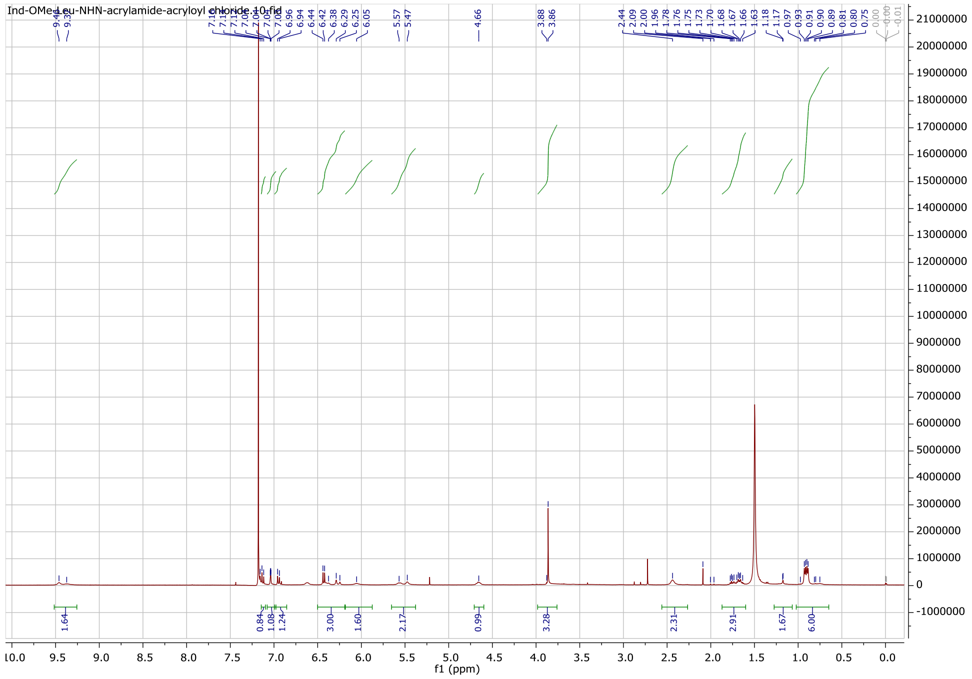

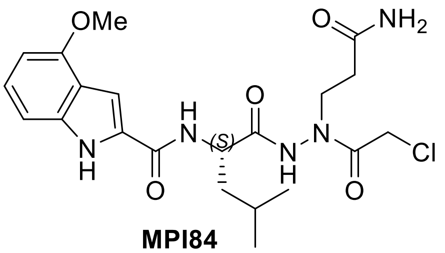
OMe
O
NH2
H
N
(S)
O
N
N
H
Cl
N
H
O
O

**MPI84**

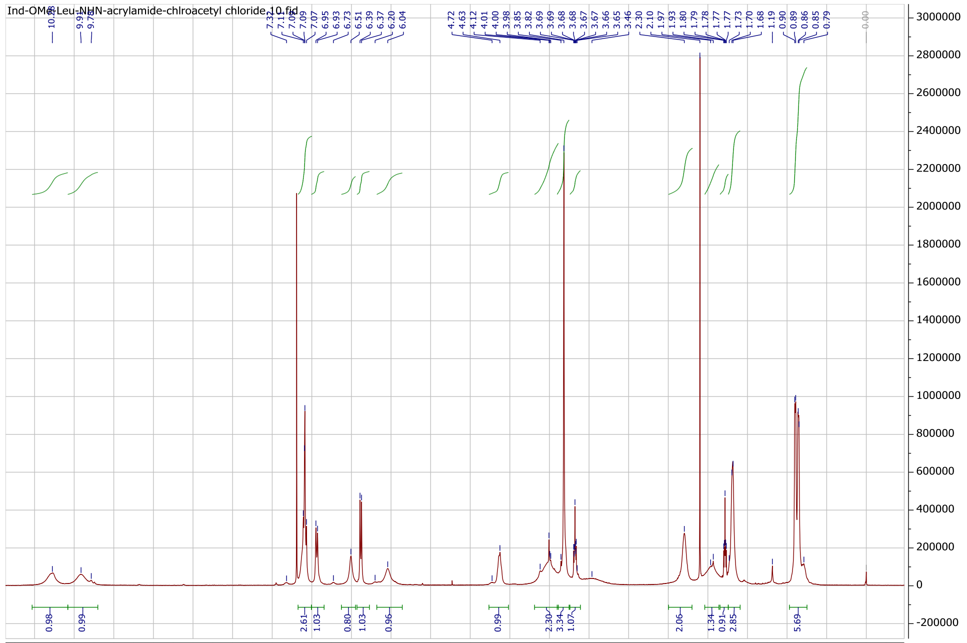
Ind-OMe-Leu-NHN-acrylamide-chlroacetyl chloride.10.fid
f1 (ppm)

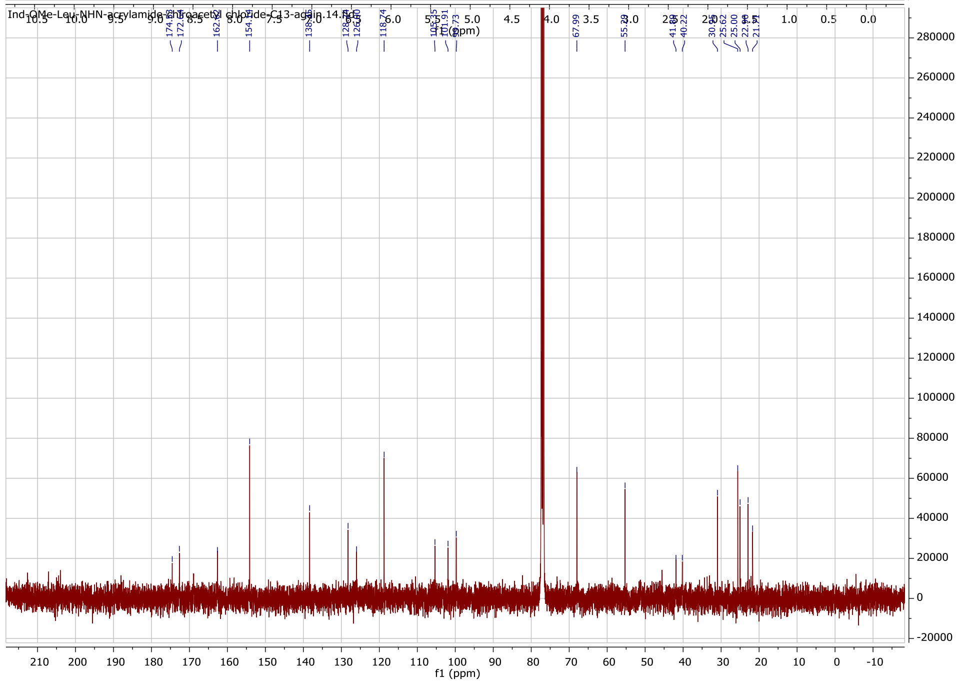
f1 (ppm)

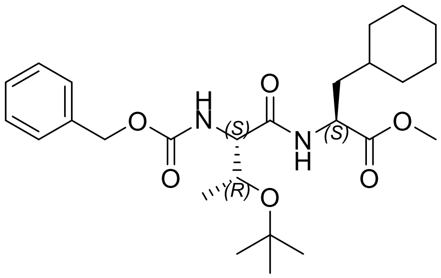

**85c**

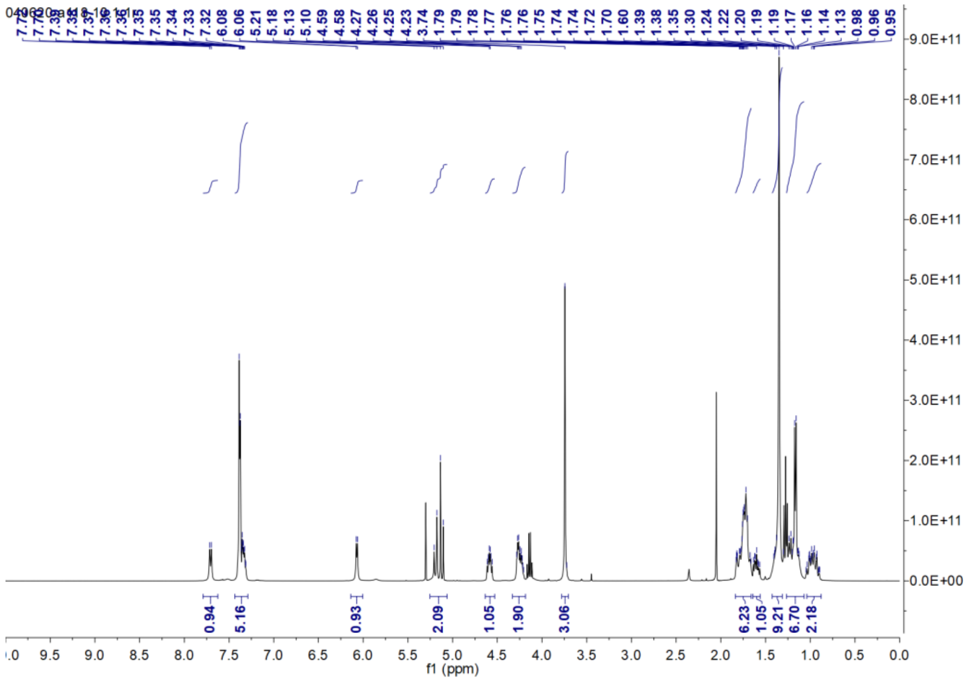

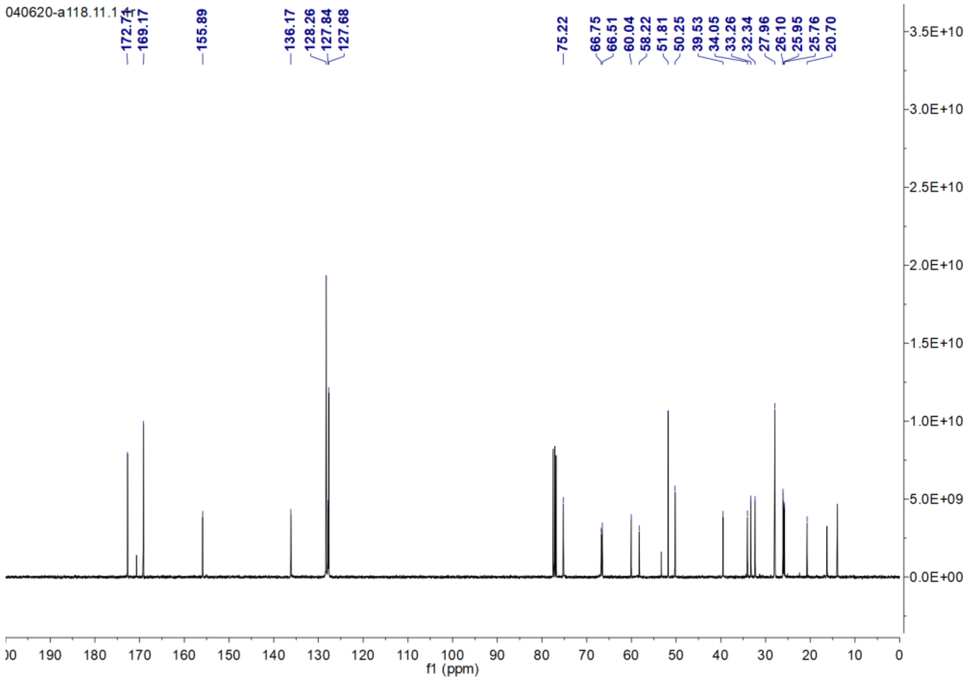

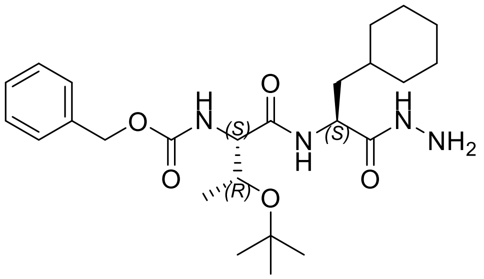

**85d**

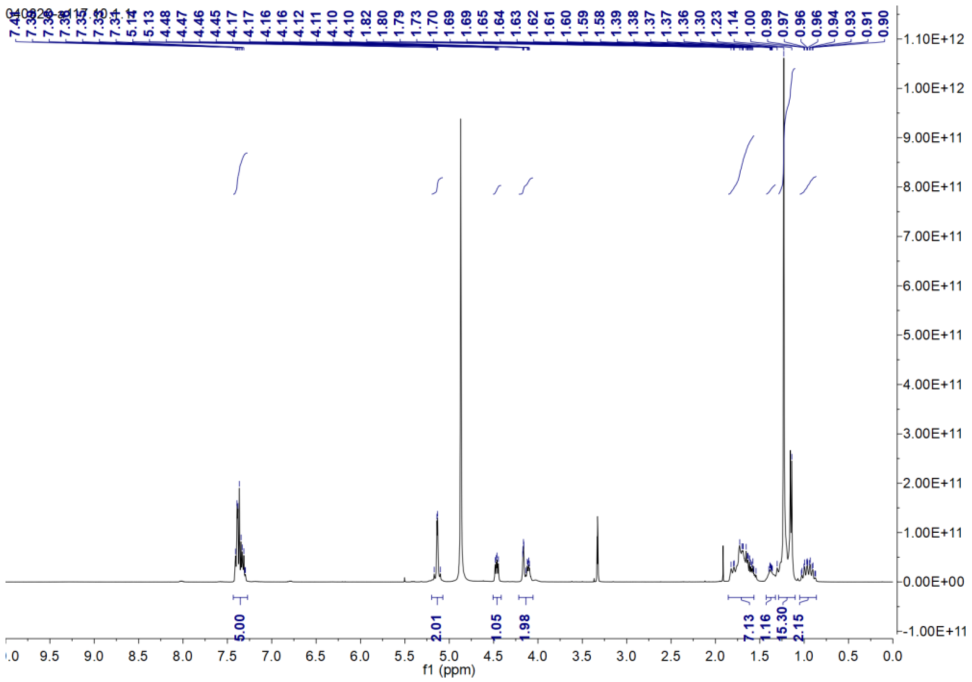
040820-a117.10.1.1
5.00
2.01
1.05
1.98
7.13
1.16
15.30
2.15
f1 (ppm)

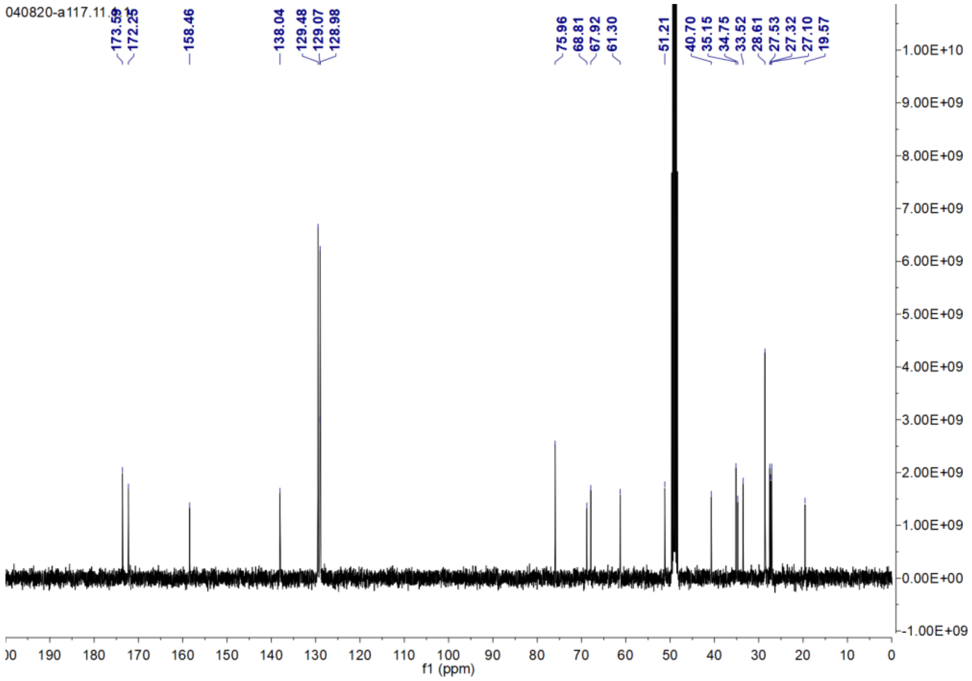
040820-a117.11.1.1
173.59
172.25
158.46
138.04
129.48
129.07
128.98
75.96
68.81
67.92
61.30
51.21
40.70
35.15
34.75
33.52
28.61
27.53
27.32
27.10
19.57
f1 (ppm)

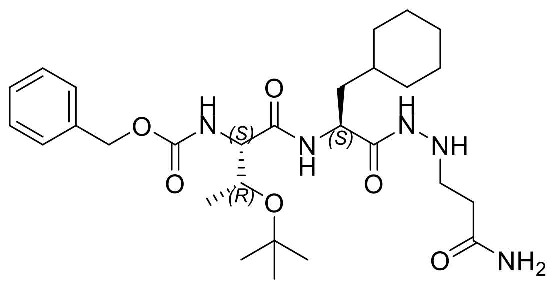

**85e**

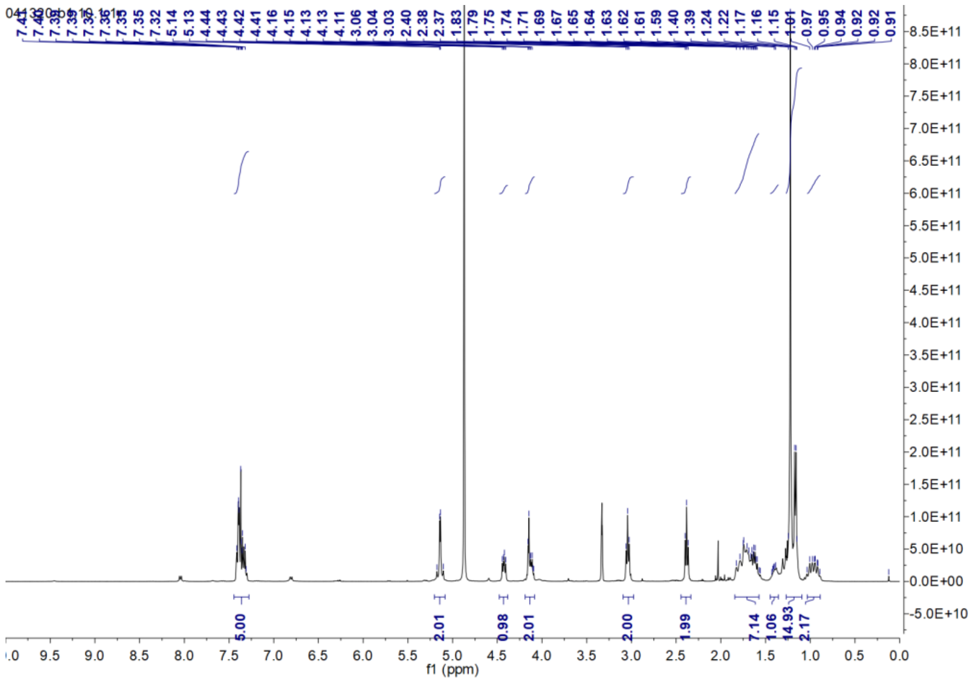

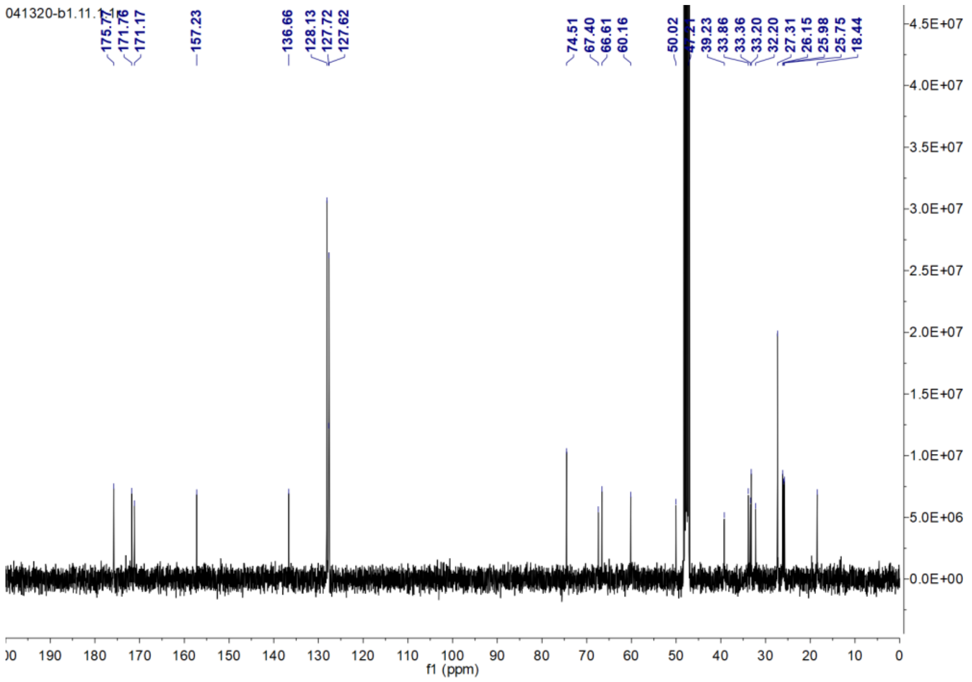

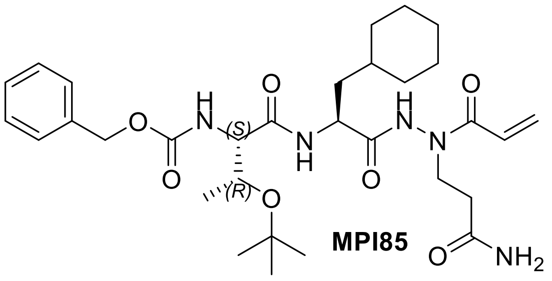

**MPI85**

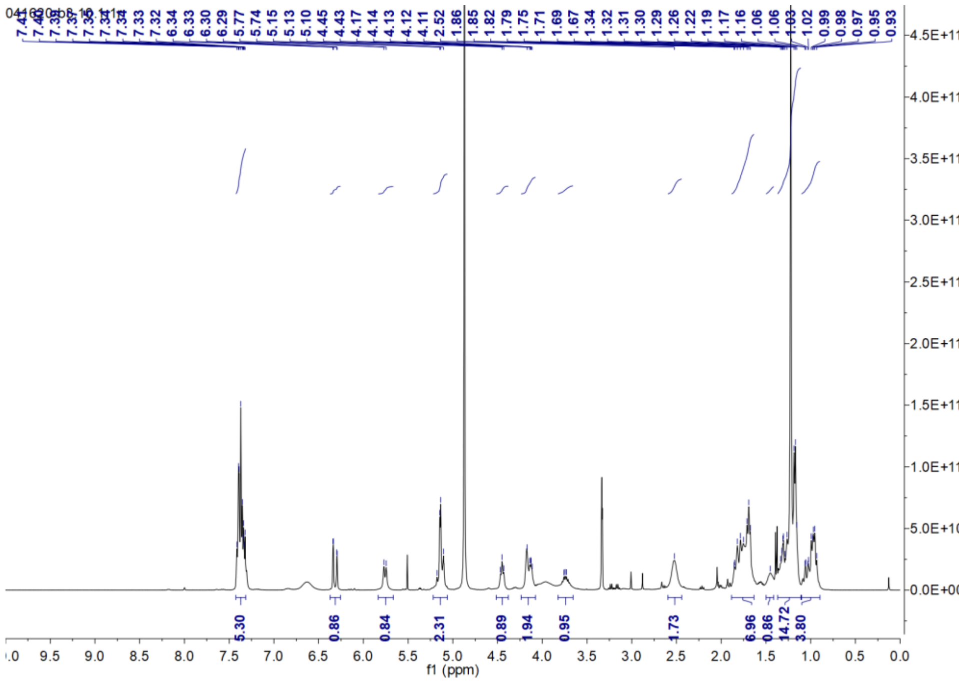
041620-b3.10.1.1r
7.41 7.40 7.38 7.37 7.36 7.35 7.34 7.33 7.32 6.34 6.33 6.30 6.29 5.77 5.74 5.15 5.13 5.10 4.45 4.43 4.17 4.14 4.13 4.12 4.11 2.52 1.86 1.85 1.82 1.79 1.75 1.71 1.69 1.67 1.34 1.32 1.31 1.30 1.29 1.26 1.22 1.19 1.17 1.16 1.06 1.06 1.03 1.02 0.99 0.98 0.97 0.95 0.93
5.30 0.86 0.84 2.31 0.89 1.94 0.95 1.73 6.96 0.86 14.72 3.80
f1 (ppm)

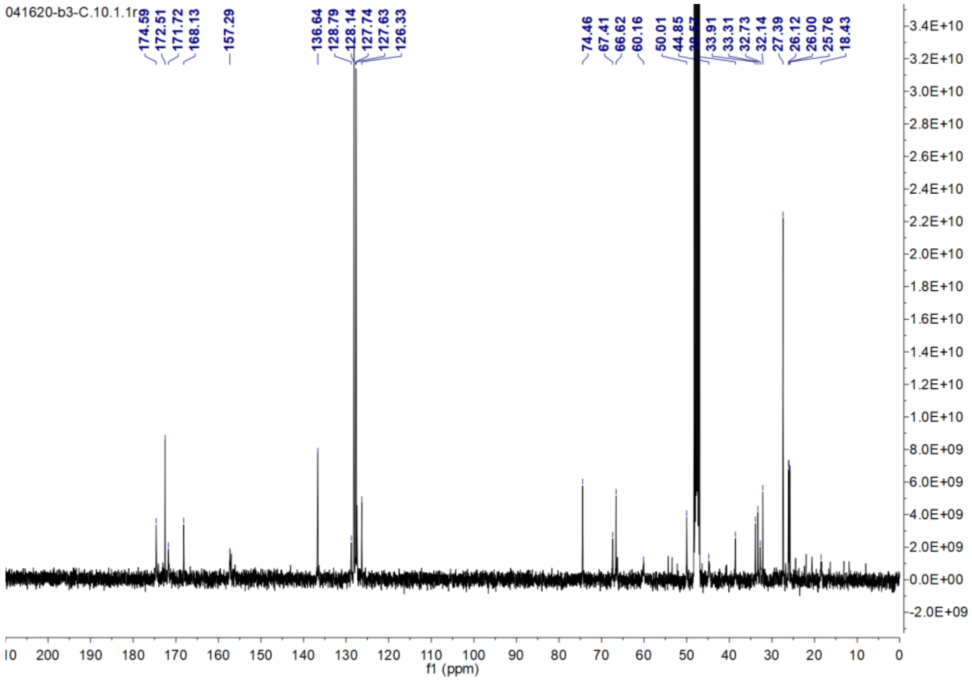
041620-b3-C.10.1.1r
174.59 172.51 171.72 168.13 157.29 136.64 128.79 128.14 127.74 127.63 126.33 74.46 67.41 66.62 60.16 50.01 44.85 33.91 33.31 32.73 32.14 27.39 26.12 26.00 25.76 18.43
f1 (ppm)

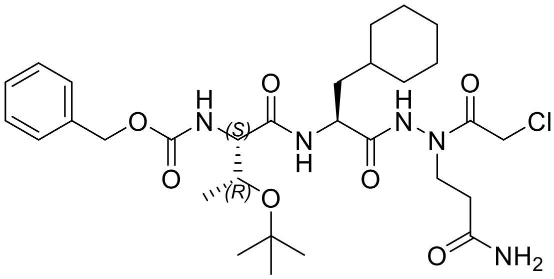

**MPI86**

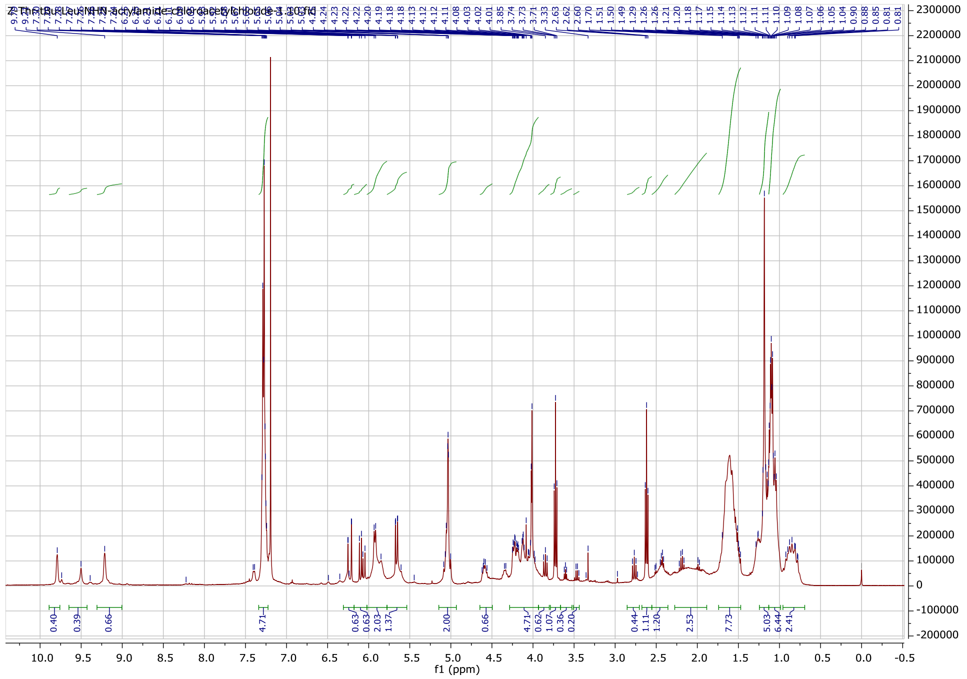

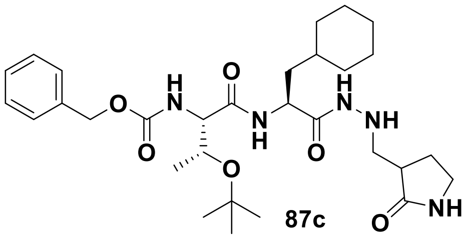

**87c**

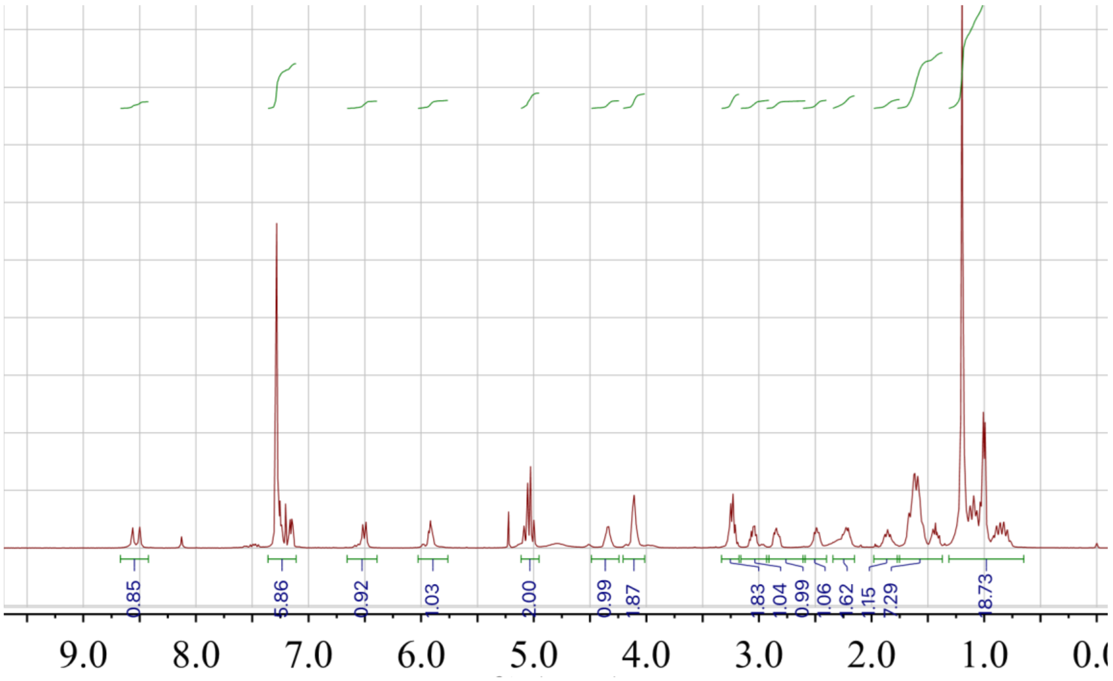

0.85
5.86
0.92
1.03
2.00
0.99
1.87
1.83
1.04
0.99
1.06
1.62
1.15
7.29
18.73
9.0
8.0
7.0
6.0
5.0
4.0
3.0
2.0
1.0

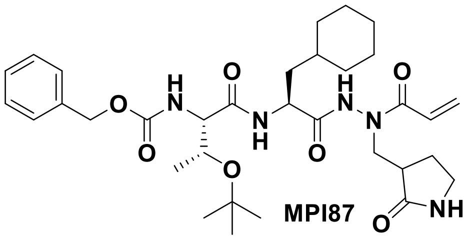

MPI87

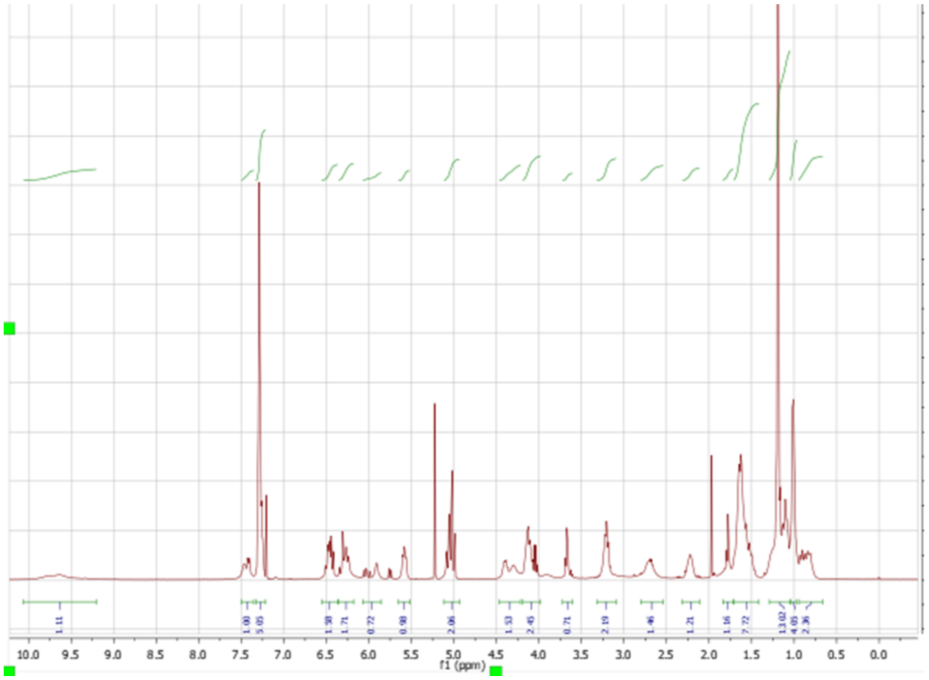
1.11
1.00
5.05
1.58
1.71
0.72
0.98
2.06
1.53
2.45
0.71
2.19
1.46
1.21
1.16
7.72
13.02
4.05
2.36
10.0 9.5 9.0 8.5 8.0 7.5 7.0 6.5 6.0 5.5 5.0 4.5 4.0 3.5 3.0 2.5 2.0 1.5 1.0 0.5 0.0
f1 (ppm)

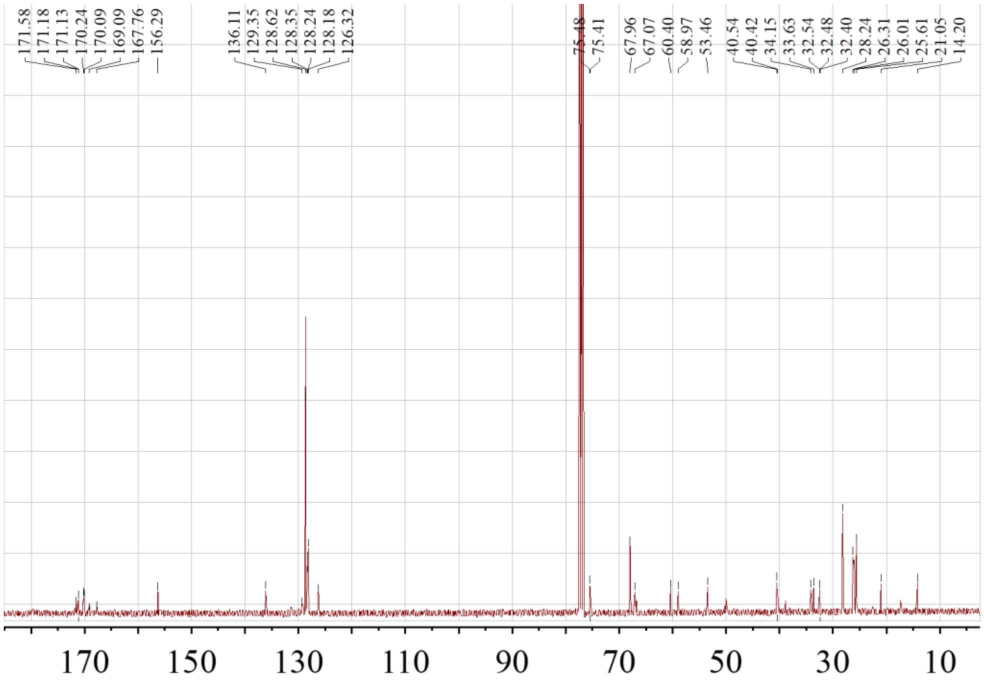
171.58
171.18
171.13
170.24
170.09
169.09
167.76
156.29
136.11
129.35
128.62
128.35
128.24
128.18
126.32
75.48
75.41
67.96
67.07
60.40
58.97
53.46
40.54
40.42
34.15
33.63
32.54
32.48
32.40
28.24
26.31
26.01
25.61
21.05
14.20
170
150
130
110
90
70
50
30
10

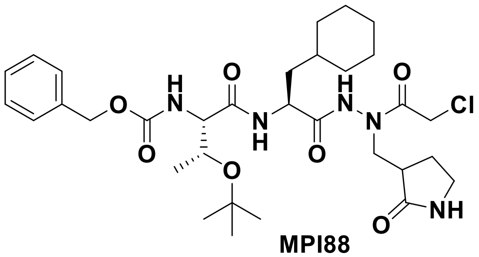

**MPI88**

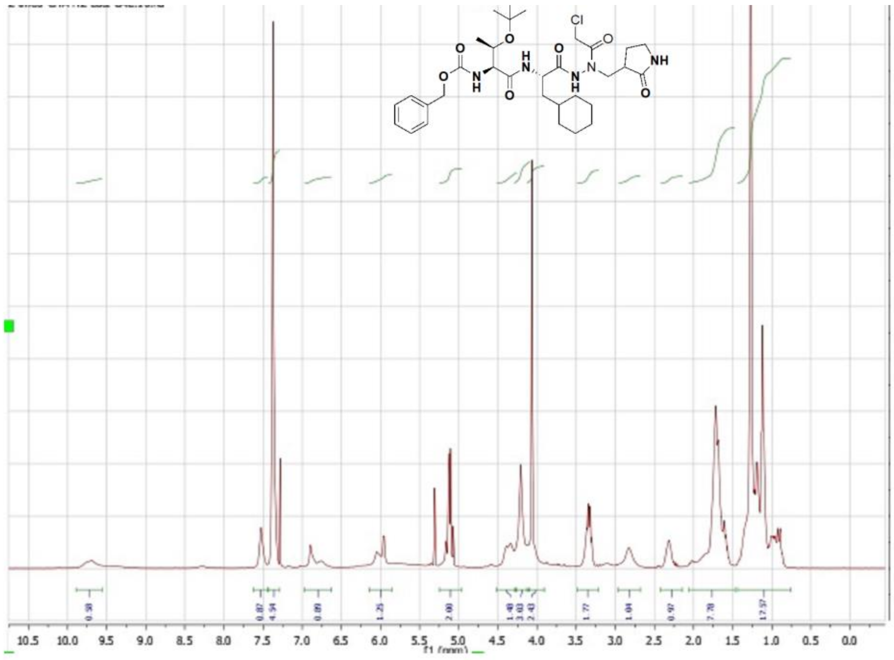
10.5 10.0 9.5 9.0 8.5 8.0 7.5 7.0 6.5 6.0 5.5 5.0 4.5 4.0 3.5 3.0 2.5 2.0 1.5 1.0 0.5 0.0
f1 (ppm)

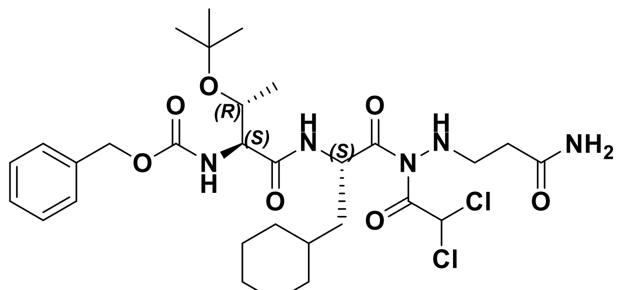

**MPI89**

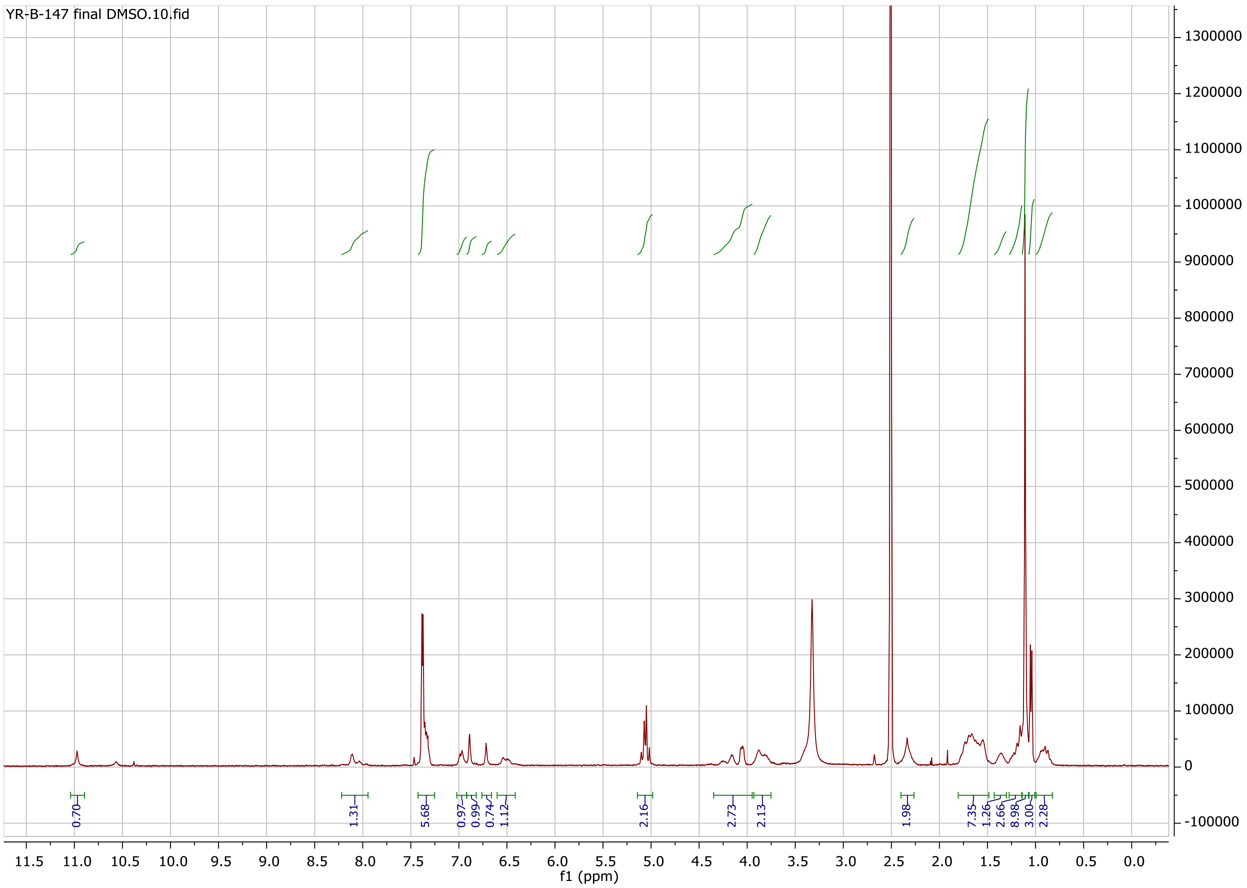

YR-B-147 final DMSO.10.fid
0.70
1.31
5.68
0.97
0.99
0.74
1.12
2.16
2.73
2.13
1.98
7.35
1.26
2.66
8.98
3.00
2.28
11.5 11.0 10.5 10.0 9.5 9.0 8.5 8.0 7.5 7.0 6.5 6.0 5.5 5.0 4.5 4.0 3.5 3.0 2.5 2.0 1.5 1.0 0.5 0.0
f1 (ppm)
1300000
1200000
1100000
1000000
900000
800000
700000
600000
500000
400000
300000
200000
100000
0
-100000

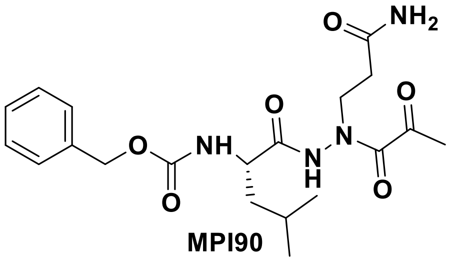

**MPI90**

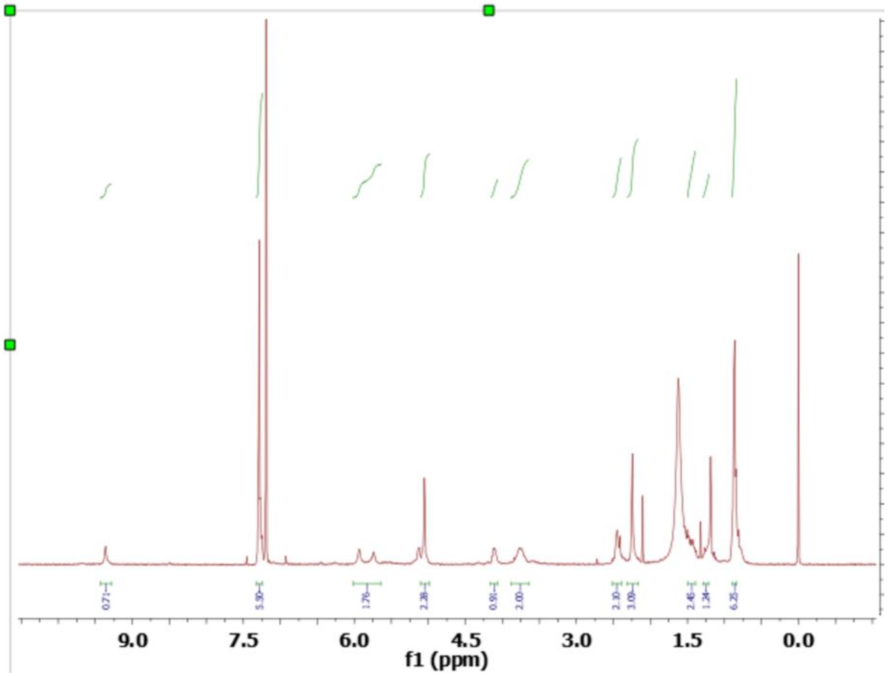
0.71
5.30
1.76
2.38
0.91
2.00
2.30
3.08
2.45
1.24
6.25
9.0
7.5
6.0
4.5
3.0
1.5
0.0
f1 (ppm)

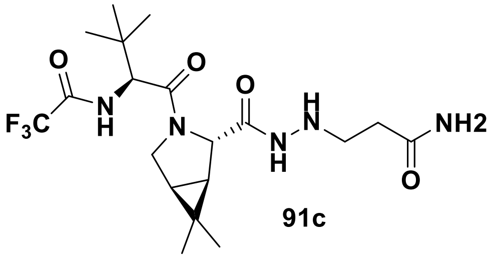
O
O
O
H
N
F3C
N
H
N
N
H
NH2
O

**91c**

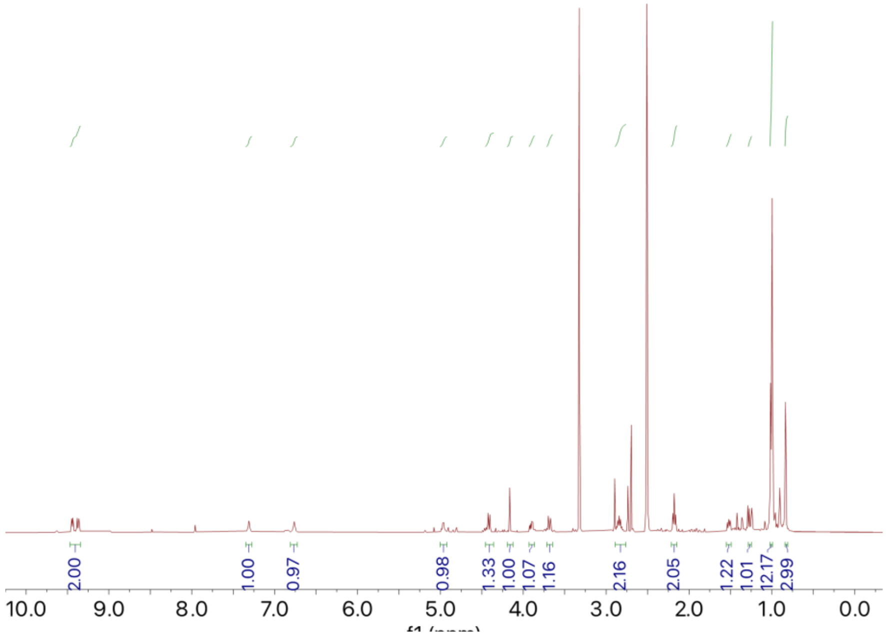
2.00
1.00
0.97
0.98
1.33
1.00
1.07
1.16
2.16
2.05
1.22
1.01
12.17
2.99
10.0
9.0
8.0
7.0
6.0
5.0
4.0
3.0
2.0
1.0
0.0

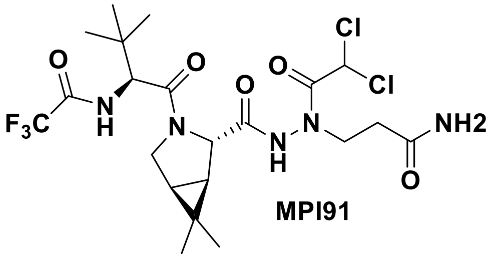

O
O
O
O
Cl
Cl
NH2
F3C
N
H
N
N
H
N
O


MPI91

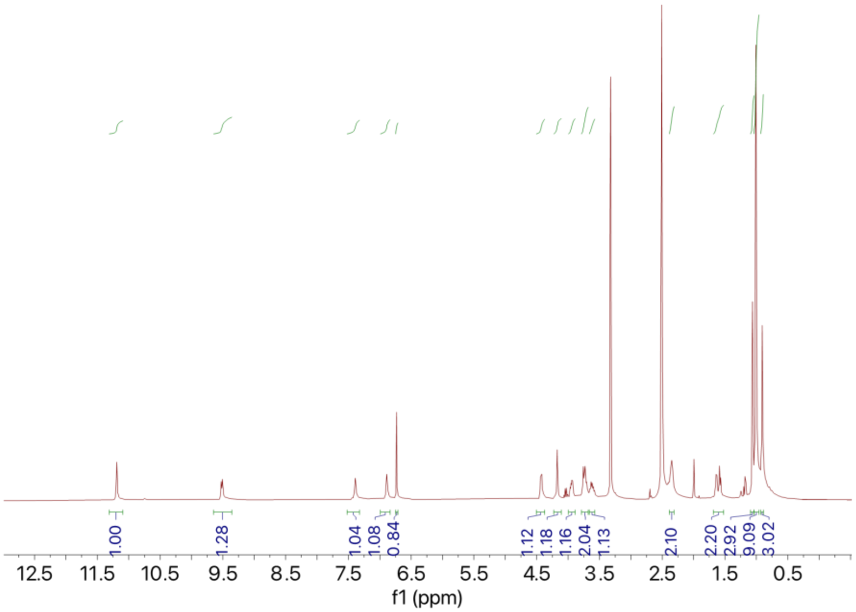

1.00
1.28
1.04
1.08
0.84
1.12
1.18
1.16
2.04
1.13
2.10
2.20
2.92
9.09
3.02
12.5
11.5
10.5
9.5
8.5
7.5
6.5
5.5
4.5
3.5
2.5
1.5
0.5
f1 (ppm)

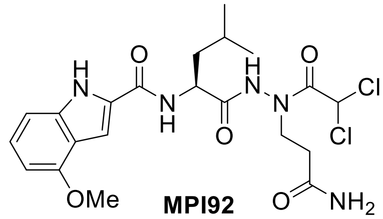

**MPI92**

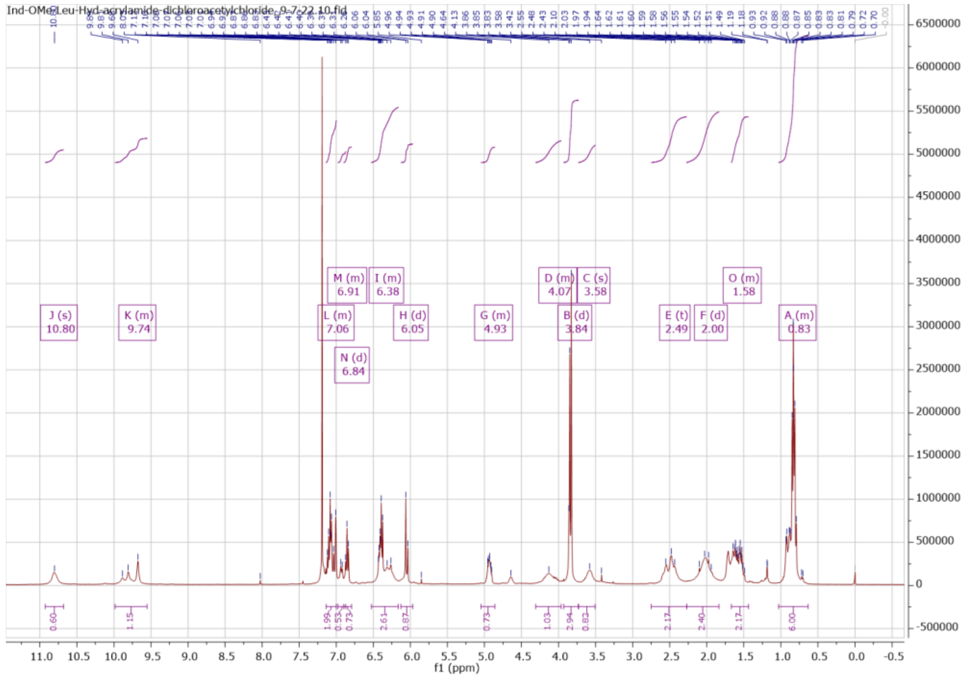
Ind-OMe-Leu-Hyd-acrylamide-dichloroacetylchloride_9-7-22.10.fid
J (s)
10.80
K (m)
9.74
M (m)
6.91
I (m)
6.38
L (m)
7.06
H (d)
6.05
N (d)
6.84
G (m)
4.93
D (m)
4.07
C (s)
3.58
B (d)
3.84
E (t)
2.49
F (d)
2.00
O (m)
1.58
A (m)
0.83
f1 (ppm)

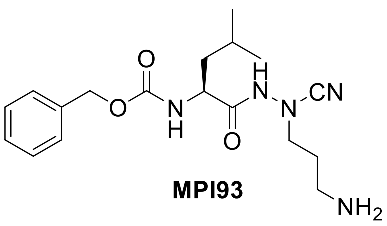

**MPI93**

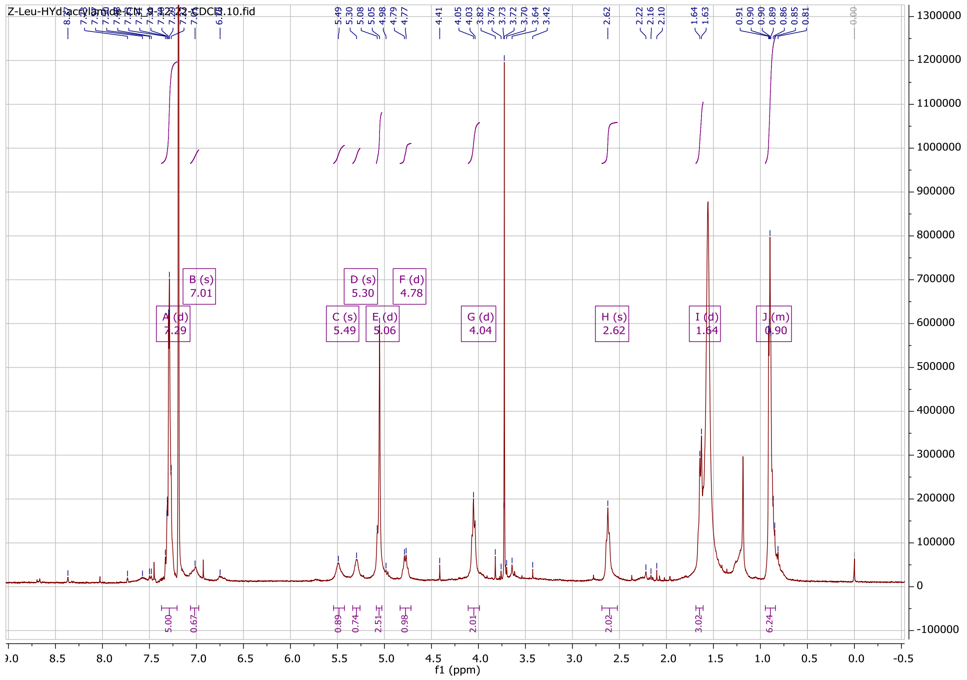

Z-Leu-HYd-acrylamide-CN_9-12-22-CDCl3.10.fid
f1 (ppm)


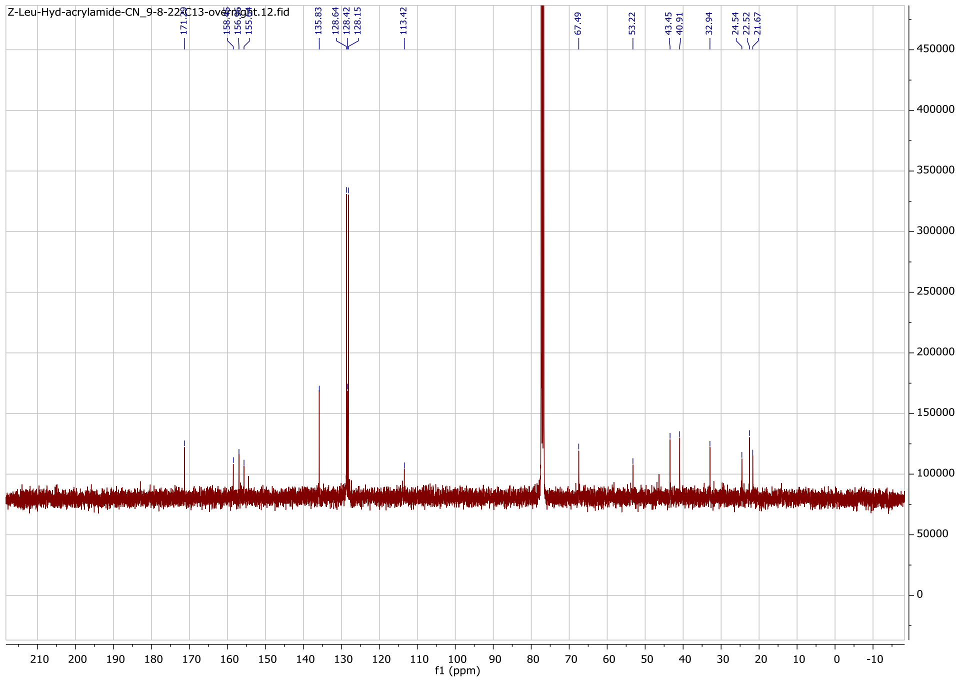

Z-Leu-Hyd-acrylamide-CN_9-8-22-C13-overnight.12.fid
f1 (ppm)

### LC-MS data of selected inhibitors

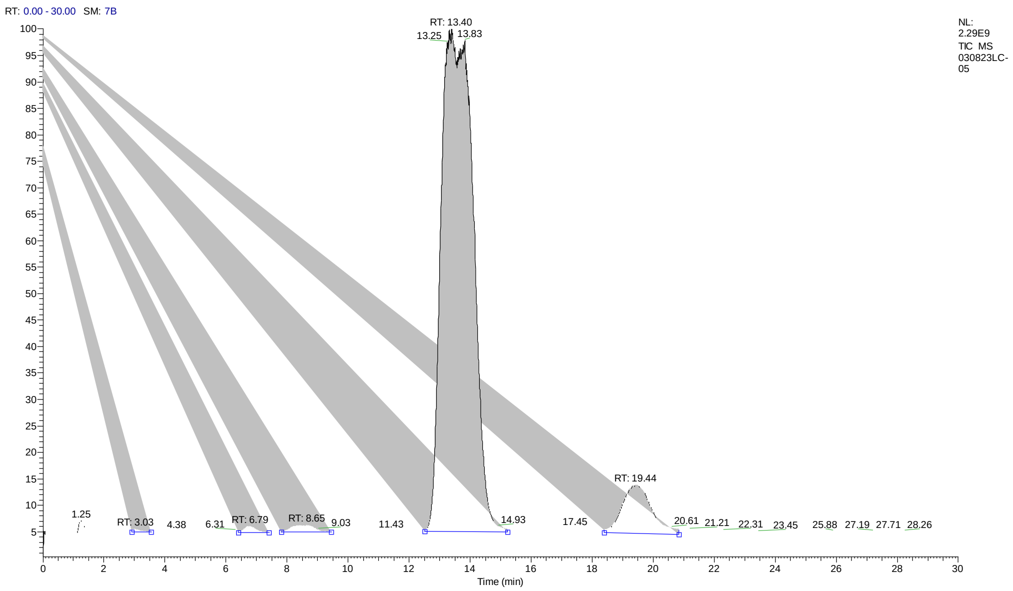


PEAK
LIST
030823LC-05.raw
RT: 0.00 - 30.00
Number of detected peaks: 5

| Apex RT | Start RT | End RT | Area | %Area | **%Purity** |
|---|---|---|---|---|---|
| 3.03 | 2.82 | 3.51 | 2.3E+08 | 0.13 | |
| 6.79 | 6.09 | 7.43 | 8.05E+08 | 0.47 | |
| 8.65 | 7.8 | 9.25 | 1.56E+09 | 0.91 | |
| 13.4 | 12.5 | 15.23 | 1.56E+11 | 91.16 | **91%** |
| 19.44 | 18.5 | 20.64 | 1.26E+10 | 7.32 | |

### MPI88

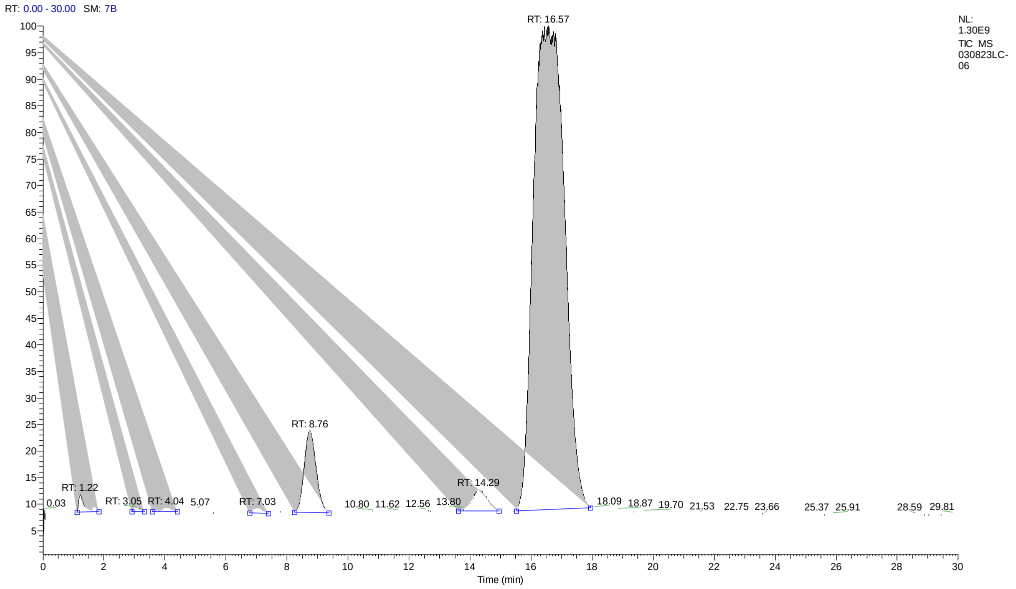


PEAK
LIST
030823LC-06.raw
RT: 0.00 - 30.00
Number of detected peaks: 7

| Apex RT | Start RT | End RT | Area | %Area | **%Purity** |
|---|---|---|---|---|---|
| 3.05 | 2.93 | 3.42 | 1.62E+08 | 0.18 | |
| 4.04 | 3.58 | 4.43 | 2.39E+08 | 0.26 | |
| 5.07 | 4.81 | 5.57 | 1.3E+08 | 0.14 | |
| 7.03 | 6.76 | 7.49 | 2.44E+08 | 0.27 | |
| 8.76 | 8.23 | 9.37 | 5.4E+09 | 5.89 | |
| 14.29 | 13.59 | 15.14 | 1.93E+09 | 2.1 | |
| 16.57 | 15.52 | 17.93 | 8.36E+10 | 91.16 | **91%** |

### MPI89

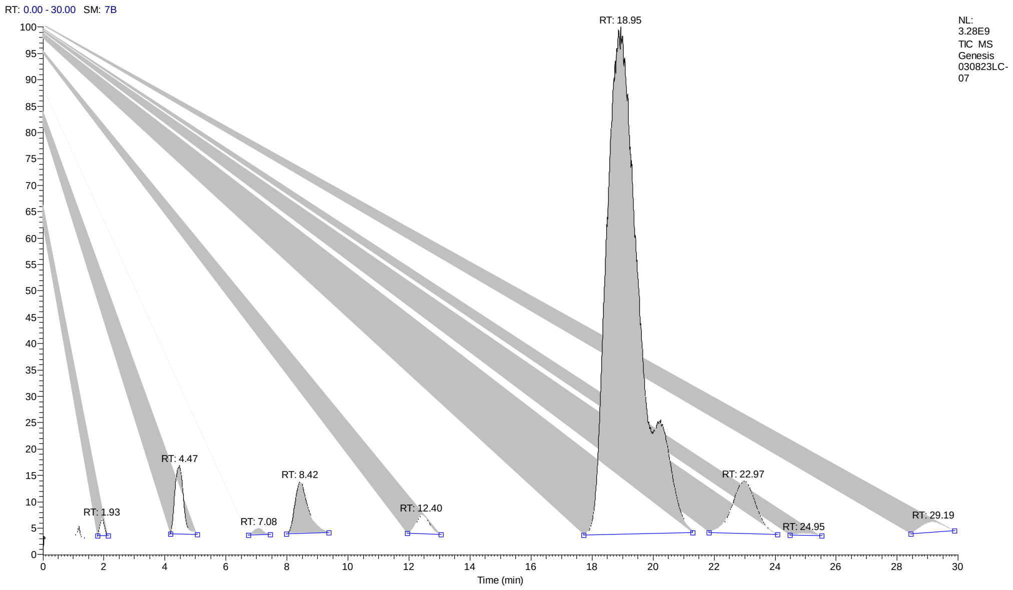


PEAK
LIST
030823LC-07.raw
RT: 0.00 - 30.00
Number of detected peaks:
13

| Apex RT | Start RT | End RT | Area | %Area | **%Purity** |
|---|---|---|---|---|---|
| 3.07 | 2.52 | 3.71 | 1.5E+09 | 0.52 | |
| 4.47 | 4.17 | 5.06 | 8.29E+09 | 2.89 | |
| 5.36 | 5.19 | 5.66 | 2.29E+08 | 0.08 | |
| 6.29 | 5.96 | 6.62 | 2.04E+08 | 0.07 | |
| 7.09 | 6.68 | 7.55 | 1.17E+09 | 0.41 | |
| 8.42 | 7.99 | 9.37 | 9.77E+09 | 3.41 | |
| 9.85 | 9.51 | 10.28 | 4.38E+08 | 0.15 | |
| 11.06 | 10.67 | 11.78 | 6.46E+08 | 0.23 | |
| 12.4 | 11.86 | 13.21 | 3.8E+09 | 1.33 | |
| 14.57 | 13.96 | 15.33 | 4.51E+08 | 0.16 | |
| 18.95 | 17.71 | 21.29 | 2.36E+11 | 82.56 | **83%** |
| 22.97 | 21.82 | 24.08 | 1.78E+10 | 6.21 | |
| 29.19 | 28.08 | 30 | 5.68E+09 | 1.98 | |
